# Climate warming changes synchrony of plants and pollinators

**DOI:** 10.1101/2021.01.10.425984

**Authors:** Jonas Freimuth, Oliver Bossdorf, J. F. Scheepens, Franziska M. Willems

**Affiliations:** Plant Evolutionary Ecology, Institute of Evolution and Ecology, University of Tübingen, Auf der Morgenstelle 5, 72076 Tübingen, Germany; Plant Evolutionary Ecology, Faculty of Biological Sciences, Goethe University Frankfurt, Max-von-Laue-Str. 13, 60438 Frankfurt am Main, Germany; Conservation Biology, University of Marburg, Marburg, Germany

**Author notes:** **Correspondence:** Franziska Willems.

**Keywords:** asynchrony, GBIF, mismatch, phenology, pollination mode

## Abstract

Climate warming changes the phenology of many species. When interacting organisms respond differently, climate change may disrupt their interactions and affect the stability of ecosystems. Here, we used GBIF occurrence records to examine phenology trends in plants and their associated insect pollinators in Germany since the 1980s. We found strong phenological advances in plants, but differences in the extent of shifts among pollinator groups. The temporal trends in plant and insect phenologies were generally associated with interannual temperature variation, and thus likely driven by climate change. When examining the synchrony of species-level plant-pollinator interactions, their temporal trends differed among pollinator groups. Overall, plant-pollinator interactions become more synchronized, mainly since the phenology of plants responded more strongly to climate change than that of the pollinators. However, if the observed trends continue, many interactions may become more asynchronous again in the future. Our study suggests that climate change affects the phenologies of both plants and insects, and that it also influences the synchrony of plant-pollinator interactions.

## Introduction

Phenological events are periodically occurring events in the life cycle of organisms. The timing of these events often depends on environmental factors such as temperature or photoperiod, and it is well known that climate change affects some of these and thus changes the phenologies of many organisms [1,2]. With such phenology shifts, there is increasing concern about possible phenological mismatches between interacting organisms, which could exceed the natural resilience of ecosystems [3,4]. Climate change-induced phenological shifts have been documented extensively for individual species [5], but we still know much less about how these shifts affect ecological interactions. Kharouba et al. [6] recently reviewed 54 published interaction studies across ecosystems and interaction types and found no clear general trend, with about half of the studied interactions becoming more asynchronous but the other half becoming even more synchronized through climate change.

Plant-pollinator systems are among the biotic interactions expected to suffer most from a mismatch of phenological events [7]. Several previous studies have observed mismatches [8,9], but in others plants and pollinators seemed to be able to keep up with each other [10]. An interesting question in this context is also which of the two partners is advancing faster if there is an increasing mismatch. So far, the evidence here is also mixed. For instance Gordo & Sanz [11] found pollinators to advance faster than trees, and Parmesan [5] that butterflies advanced faster than herbaceous plants, but in a study by Kudo & Ida [12] it was the plants – spring ephemerals – that advanced faster than their bee pollinators.

Mismatches of plant-pollinator interactions can have negative consequences for both partners. For the pollinators, this can include lower survival rates, a decreased overall fitness and higher parasite loads [13]. Moreover, mismatches might also impact pollinator demography, the body sizes [8] and frequencies of sexes, and thus population viability [13]. On the plant side, desynchronized pollinator interactions are mainly expected to impact plant fitness and thus long-term population growth and survival. For instance, Kudo & Ida [12] found that seed counts were reduced in early-flowering spring ephemerals after desynchronization with their bee pollinators. However, in another study fly-pollinated plants did not show similar responses [14].

Testing hypotheses about plant-pollinator responses to climate change is not trivial. Since changes in phenology take place on the scale of decades [15], we need long-term data. A possible source of long-term data on plant phenology are herbarium specimens [16,17], which can indicate the day of year that a specific species was flowering in a given location and year. Herbarium data provide unique historical depth, but they need to be treated with caution because of the sampling biases associated with them [18,19]. In recent years the digitization of herbaria as well as other collections and observation data, including on other taxa such as pollinating insects, e.g. from long-term monitoring networks, is creating an increasing number of public data bases that contain vast amounts of natural history data of large spatial and temporal scales [20]. These data are increasingly used for analyses of broad ecological trends and global changes [19,21]. One of the largest and most important hubs of large-scale and long-term ecological data sets is the Global Biodiversity Facility (GBIF), an intergovernmental initiative and public data base that provides access to biodiversity data compiled from various individual sources like academic institutions, government agencies or independent collections [22].

Another matter is finding a measure for changes in phenology. Primack et al. [23] demonstrated that the collection dates of herbarium specimens of a plant species in a year can be used as a proxy for flowering time in that year. The same approach of using occurrence records in natural history collections or other data bases can in principle be used to estimate the activity times of other groups of organisms such as insects ([6] and references therein). For instance, analyses of natural history collections in the UK have demonstrated phenology changes in bees [9] and butterflies [24]. Thus, the occurrences of plants and insects in GBIF may be used to estimate activity shifts of different taxonomic groups, as well as their synchrony. When we use the term ‘activity’ in this paper, we refer to the period in an organism’s life when it can interact with its ecological partner. For plants this is the period of flowering, for insect pollinators, such as bees, flies, beetles, and butterflies, it is the period of flight.

We used data from GBIF to study phenological shifts of plants and insect pollinators in Germany, at the level of taxonomic groups as well as individual species’ interactions. We asked the following questions: (i) Are there long-term trends in the phenology of plants and pollinators? (ii) If yes, are phenology trends related to climate change? (iii) How does climate change affect the synchrony of interactions between individual plant and pollinator species?

## Methods

### Phenology data

We worked with occurrence records of plants and insects available from the GBIF database [25–28]. For the plants, we restricted ourselves to species covered by the BiolFlor database of plant traits [29], because we originally intended to classify plants by their level of pollinator dependence – an idea we later abandoned. For the insects we restricted ourselves to beetles (Coleoptera), flies (Diptera), bees (Hymenoptera) as well as butterflies and moths (Lepidoptera), as these groups contain most insect pollinators [30]. We used the R package *rgbif* [31] to download all available records of the above taxa from GBIF. Our basic criteria for including records were that they originated from Germany, and that their basis of record (as defined in GBIF) was either a living specimen (e.g., a captured insect), a human observation (i.e., an observation of a species made without collecting it), just an observation (i.e., when the exact type of observation was not clear), or a preserved specimen (e.g., an herbarium record or a collected specimen). If names of plant species were not accepted names, we used the R package *taxsize* [32] to check the names against the GBIF backbone taxonomy and determine the actual accepted name.

Prior to the data analyses, we subjected the data to several steps of quality control. First, we removed all records from before 1980 as these turned out to be too inconsistent, with few records per year and large gaps due to consecutive years without records. We also removed the records from 2021 as the year had not been complete at the time of our analysis. Second, we removed all records from the days of year (DOY) 1, 95, 121, 163, 164, 166 and 181, and in particular DOY 365 and 366 from the National Museum of Natural History in Luxembourg because the high numbers of records on these days indicated that either records without collecting date had been assigned these by default, or the dates were used by BioBlitz events where very large numbers of records are taken on a specific day of the year. Including these data would have strongly biased the intra-annual distributions of our records. Finally, we removed some records for which no elevation or temperature data could be obtained (see below).

To ensure reasonable coverage of the studied time interval, we then restricted the records to species which had at least 10 occurrence records in every decade covered (with the year 2020 included in the last decade).

After these data curation steps, we maintained just above 12 million occurrence records that covered altogether 1,764 species, with 11.4 million records of 1,438 plant species, around 590,000 records of 207 species of butterflies and moths, some 76,000 records of 20 bee species and 30,000 records of 22 fly species, and almost 25,000 records of 77 species of beetles (**Table 1**). There were large differences between plants and insects not only in the numbers of records but also in their temporal distribution across the studied period (**Figure S 1**). While plants had relatively even record numbers across years, the insect groups, in particular flies and bees, were strongly underrepresented in the earlier decades, and record numbers increased rapidly in the last 20 years, probably due to the advent of platforms like iNaturalist.org and naturgucker.de, which allow recording of species occurrences by citizen naturalists, and which made up most of the insect occurrence data for Germany in GBIF.

**Table 1.** Overview of the studied taxonomic groups, their numbers of species and total records, ranges of records per species, and the average temporal shifts and temperature sensitivities (± 1 S.E.) of their phenology, the significance level of the slope estimate, and the respective fractions of species that showed a negative slope in their shifts and temperature sensitivities.

| Tax. group | # Species | # Records | # Records/Species |  |  | Temporal shifts |  |  | Temperature sensitivity |  |  |
| --- | --- | --- | --- | --- | --- | --- | --- | --- | --- | --- | --- |
|  |  |  | Min | Median | Max | Days / decade | *P*-value | % Species Advancing | Days / 1°C | *P*-value | % Species Advancing |
| Beetles | 77 | 24,758 | 55 | 196 | 1,879 | $0.9 \pm 0.9$ | 0.340 | 44% | $-1.7 \pm 0.5$ | < 0.001 | 74% |
| Flies | 22 | 30,927 | 166 | 675 | 9,194 | $-5.0 \pm 0.9$ | < 0.001 | 91% | $-3.9 \pm 0.7$ | < 0.001 | 91% |
| Bees | 20 | 76,438 | 91 | 599 | 15,059 | $-2.6 \pm 2.0$ | 0.270 | 65% | $-2.0 \pm 1.3$ | 0.140 | 60% |
| Butterflies | 206 | 589,970 | 69 | 799 | 34,065 | $-3.9 \pm 0.3$ | < 0.001 | 86% | $-1.9 \pm 0.2$ | < 0.001 | 74% |
| Plants | 1,438 | 11,357,981 | 62 | 2,280 | 121,612 | $-7.2 \pm 0.2$ | < 0.001 | 86% | $-5.2 \pm 0.2$ | < 0.001 | 84% |

### Temperature and elevation data, and individual interactions

Besides the main phenology data from GBIF records, we obtained several other data sets required for our analyses. To test for associations with climate, we used temperature data from the Climate Research Unit (CRU, https://crudata.uea.ac.uk) at the University of East Anglia, specifically the Time-Series dataset version 4.05 [33], which contains gridded temperature data from 1901-2020 at a resolution of 0.5° longitude by 0.5° latitude. From this dataset we extracted the monthly mean temperatures and averaged them to obtain the annual mean temperatures at the sites of occurrence records. To be able to control for elevation at the locations of occurrence records, we used elevation data at a 90 m resolution from the NASA Shuttle Radar Topography Mission (SRTM), obtained from the SRTM 90m DEM Digital Elevation Database [34] and accessed through the *raster* package [35] in R.

Finally, we obtained data on individual plant-pollinator interactions from a United Kingdom (UK) database on plant-pollinator interactions [36] hosted by the Centre for Ecology and Hydrology (CEH). This database included all known interactions between plants and flower-visiting bees, butterflies, and hoverflies (but unfortunately neither beetles nor moths nor flies other than hoverflies) in the UK, a country similar to Germany in terms of climate and species composition. While these interaction data are unlikely to represent all possible species interactions in Germany, we could not find any similar data for our study area, and so we used this dataset as a prediction for interactions taking place in Germany. To ensure that interactions reflected pollination, we excluded plants that were known not to depend on insects for pollination. This was the case when a plant was pollinated abiotically or only through selfing, or when it reproduced exclusively vegetatively.

### Data analysis

All data wrangling and analysis was done in R [37]. Before analysing phenology data, we examined patterns of climate change in Germany through a linear model that regressed the annual mean temperature values at the collection sites (= the corresponding 0.5° × 0.5° grid cells) over time.

To understand phenology changes in plants and insects, we first estimated the phenological shifts in each taxonomic group (i.e., plants, beetles, flies, bees, and butterflies/moths) as the slopes of linear regressions linking activity, i.e., the DOY of a record to its year of observation. We estimated taxonomic group-specific phenological shifts in two linear mixed-effect models: one that estimated shifts over time and one that related phenology variation to temperature. Both models included the DOY of a record as the response variable, and the latitude, longitude, and elevation of a record as fixed effects. The temporal-change model additionally included the year of a record as a fixed effect and as a random effect across species; the temperature-change model instead included the annual mean temperature at the site of a record as a fixed effect and as a random effect across species. We used the *lme4* package [38] in R to fit these models, and assessed model fits by visually inspecting the relationships between residuals and fitted values, and between residuals and covariates (**Supplementary diagnostic plot documents**). As the random effects from the models agreed well enough with more complex generalized additive models, we considered our linear model reasonably robust.

After estimating the average taxonomic group-level phenological shifts, we compared these between different insect orders and plants through ANOVAs, using Tukey post-hoc tests for pairwise tests of differences. We assessed the normal distribution of the slopes through boxplots and checked potential outliers for their plausibility by examining the data from which these slopes were estimated. The same procedure was used in all subsequent ANOVAs. We tested for an association between temporal shifts and temperature change by calculating the Pearson correlations between the slopes of time- and temperature-relationships within each taxonomic group. To test whether a taxonomic group had an average temporal shift or average temperature sensitivity different from zero, we performed a one-sample *t*-test on the slopes in each taxonomic group for both time and temperature slopes, adjusting the *P*-values for multiple testing using the false discovery rate [39].

Finally, we analysed the synchrony of plants and pollinators using the data on individual plant-pollinator interactions. For each of the plant-insect pollinator pairs predicted from the UK dataset we calculated the differences of the slopes and intercepts (= value for plant − value for pollinator) for the taxonomic groups estimated in the temporal-change model. From this we obtained a linear equation describing the temporal change in their asynchrony, with positive values in a given year describing asynchrony with pollinators active earlier than plants. For each group, we then tested whether the average shift of asynchrony differed from zero by using a one-sample *t*-test on the slopes for time slopes, and *P*-values adjusted for multiple testing using false discovery rates. Last, we tested for differences in the average shifts between interacting groups (i.e., hoverfly-plant, bee-plant, and butterfly-plant) through ANOVAs of the temporal asynchrony shift data and Tukey post-hoc tests that examined pairwise differences between interaction groups. Together with the information about the shifts of taxonomic groups and individual species, these analyses of plant-pollinator synchrony also allowed to infer which groups were the main agent of change in interactions.

## Results

Across all collection locations, there was a strong overall trend of climate warming. The annual mean temperature increased by 0.49 °C per decade (*F*_1, 8477_ = 2038, *R*^2^_adj_ = 0.19, *P* < 0.001 for the linear model), with a total increase of ~2 °C across the study period 1980-2020 (**Figure S2**).

### Temporal trends in plant and insect phenology

Out of the five studied taxonomic groups, three showed on average significant phenological shifts towards earlier activity: plants, flies, and butterflies/moths. For beetles and bees, the trends were less consistent (**Figure 1A**; **Table 1**). The phenology of plants advanced more strongly than all of the insect pollinators except flies (**Figure 1A**). Among the insects, flies and butterflies/moths advanced most strongly (**Table 1**), with both groups showing distinct temporal shifts from the beetles (among-group differences: ANOVA, *F*_4, 1759_ = 33.44, *P* < 0.001; pairwise differences: Tukey post-hoc with *α* = 0.05). There was generally substantial variation of temporal trends within taxonomic groups: Even in plants, flies, and butterflies/moths where >85% of species advanced their phenology (**Table 1**), there were also some for which the opposite was true (**Figure 1B**). In bees and beetles, the numbers of species with advanced versus delayed phenology were more even.

**Figure 1.**
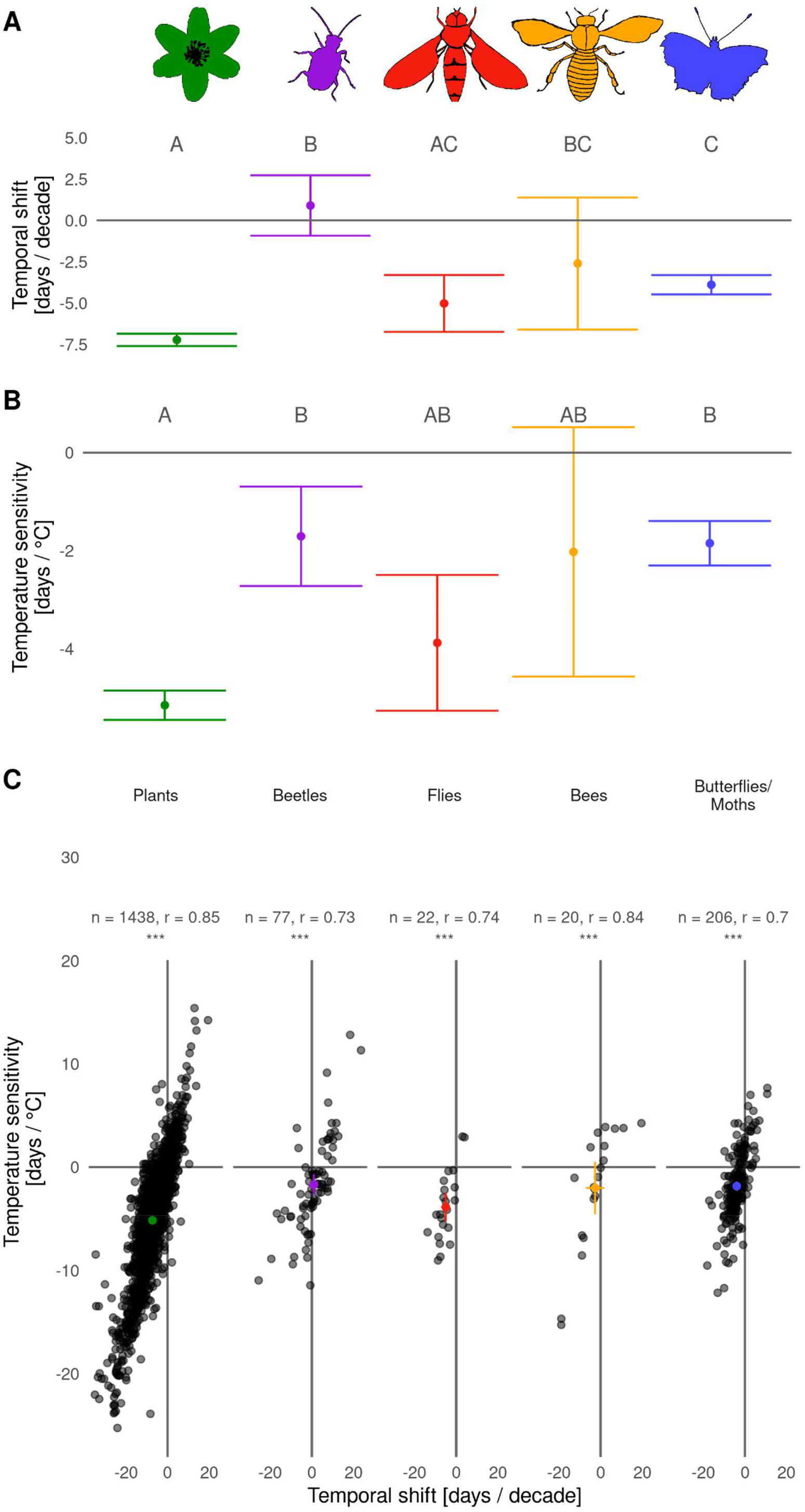
Temporal trends (A), temperature sensitivities (B), and their correlation (C) of the phenology of plants and pollinator groups in Germany from 1980 to 2020. In (A) and (B) the values are groups averages ± 95% confidence intervals, and groups with different letters above are significantly different (Tukey post-hoc tests with α = 0.05). In (C) the coloured dots are the averages of taxonomic groups, with 95% confidence intervals, and the black dots are individual species. The Pearson correlation coefficient (*r*) of each relationship is given above the plot; all correlations are significant at *P*<0.001.

### Temperature sensitivity of plant and insect phenology

When we analysed phenology variation in relation to climate instead of temporal trends, (i.e. testing for climate sensitivities), the results for the different taxonomic groups were similar, but not entirely (**Figure 1**): In addition to the plants, flies and butterflies/moths, also the beetles significantly shifted their phenology with increasing temperatures (**Table 1**), and the only groups for which the average temperature shifts differed significantly were plants and beetles, and plants and butterflies/moths, respectively (**Figure 1B**; among-group differences: ANOVA *F*_4, 1759_ = 22.19, *P* < 0.001; pairwise differences: Tukey post-hoc with *α* = 0.05). In these comparisons, the plants were generally the group with the larger temperature sensitivities. At the level of individual species, the directions and magnitudes of temperature sensitivities were generally strongly correlated with that of the temporal shifts in phenology (**Figure 1C**), i.e., species which strong climate sensitivities were also the ones that displayed large changes over time. In all taxonomic groups the majority of species showed negative temperature sensitivity, i.e., accelerating phenology at higher temperature (**Table 1**), but there was substantial variation in all groups, with a fraction of species showing the opposite responses.

### Synchrony of plant-pollinator interactions

When examining the synchrony of individual plant-pollinator interactions (predicted by the UK dataset), we found that all groups showed negative average shifts, i.e., decreasing the phenological lead of pollinators relative to the plants (**Table 2**). The three examined pollinator groups differed in their magnitudes of synchrony changes (**Figure 2A**), but, except in interactions of plants with flies, they did not become more asynchronous but rather more synchronized during the last decades (**Figure 2B**). The magnitudes of temporal changes were generally greatest for plant-bee interactions, with similar shifts for plant-butterfly and plant-hoverfly interactions (**Table 2**; ANVOA: *F*_2, 4399_ = 363.3_3985_ = 326.8, *P* < 0.001; pairwise differences: Tukey post-hoc with *α* = 0.05). In all three interaction groups, the insects tended to be the earlier partner at the start of the study period. However, since plants advanced their phenology faster than most insect groups, they tended to ‘catch up’ over time, relative to the pollinators (**Figure 2 B**). This is observable both in plant-butterfly and plant-bee interactions. and in the case of plant-hoverfly interactions the plants even ‘overtook’ the insects, with the average asynchrony values for this group becoming more negative during the 1980s, indicating that plants were gradually becoming the earlier partner in these interactions.

**Table 2.** Overview of the plant-pollinator interaction groups, the numbers of individual interactions, as well as plant and pollinator species studied in each, and the observed average temporal changes and temperature sensitivities (± 1 S.E.) of their asynchrony. All estimates in the last two columns are significantly different from zero at *P*<0.001.

| Interaction group | # Interactions | # Species |  | Temporal shifts | Temperature sensitivity |
| --- | --- | --- | --- | --- | --- |
|  |  | Plants | Pollinators | Days / decade | Days / 1 °C |
| Hoverfly - Plant | 1,407 | 219 | 17 | $-3.3 \pm 0.21$ | $-1.81 \pm 0.17$ |
| Bee - Plant | 1,008 | 256 | 12 | $-11.34 \pm 0.39$ | $-5.68 \pm 0.24$ |
| Butterfly - Plant | 1,987 | 260 | 40 | $-2.63 \pm 0.16$ | $-3.09 \pm 0.13$ |

**Figure 2.**
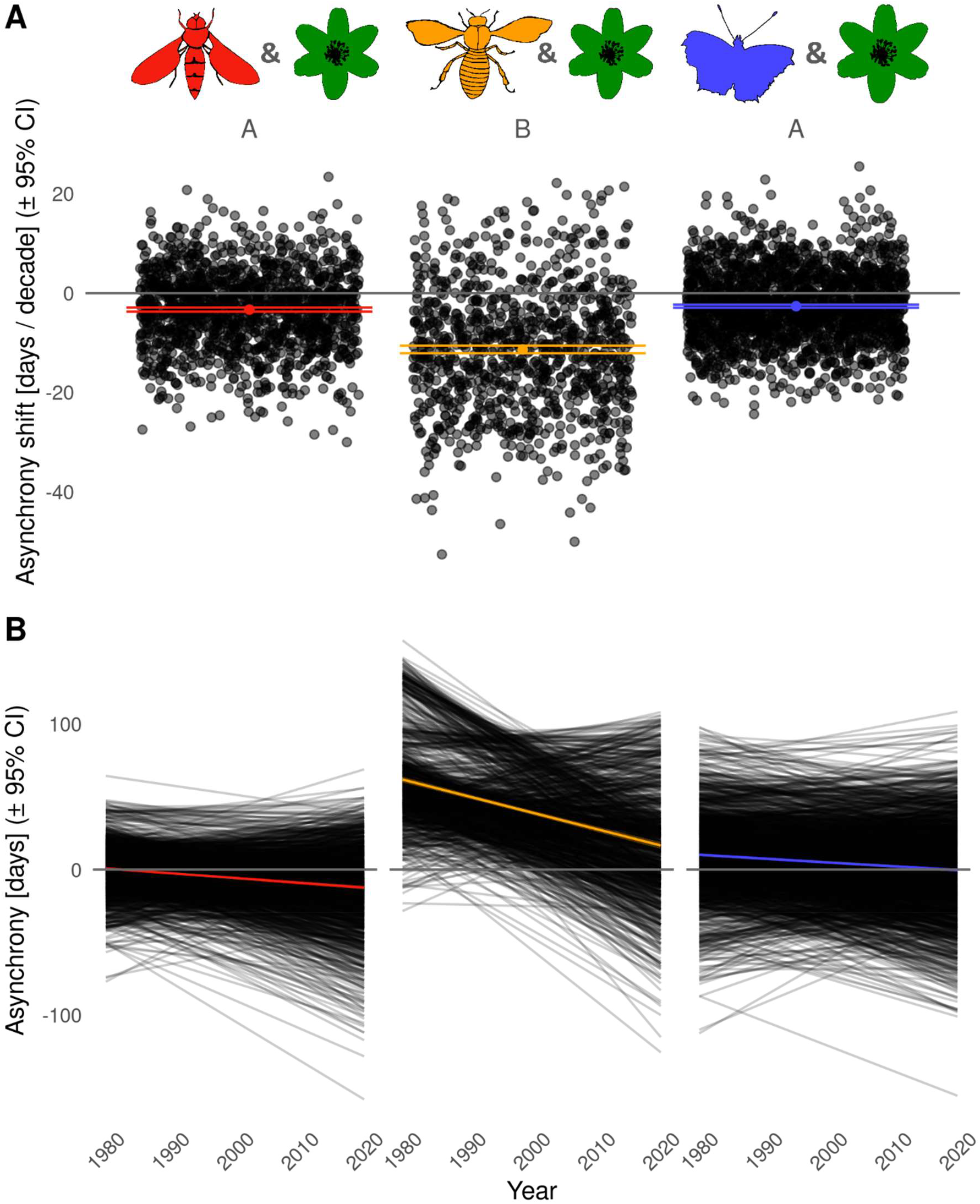
Estimated shifts in asynchrony of individual plant-pollinator interactions (predicted from a UK dataset), separately for different interaction groups. Asynchrony is the difference in the estimated yearly mean DOY of activity between the plant species and the pollinator species in an interaction. (A) Average decadal asynchrony changes of individual interactions (grey dots), and averaged for each group (coloured dots and 95% CI whiskers). Interaction groups with different letters above are significant different (Tukey post-hoc tests with α = 0.05). (B) Asynchrony changes over time (= slopes and intercepts of pollinators subtracted from plant slopes and intercepts), with black lines for individual interactions and the coloured lines for means across interaction groups.

## Discussion

In this study, we took advantage of large collections of occurrence records to examine phenological trends of flowering plants and insect pollinators in Germany. We asked whether phenology changes affected the synchrony of predicted interactions of plants and insects, and whether observed changes in phenology, and variation therein, were related to the different taxonomic groups’ responses to climate warming. Our results showed that the phenological shifts of plants and insects differed, with plants generally shifting most strongly, and substantial variation among the insect groups. These changes took place across a period in which the mean annual temperature increased in Germany. As pollinators had historically often been active before the plants, the observed faster phenology changes of the plants resulted in increased plant-pollinator synchrony in predicted plant interactions with bees and butterflies during the last decades. For interactions between plants and hoverflies, however, we find a trend towards greater asynchrony. At the level of individual species, there was generally a strong correlation between temporal shifts and temperature sensitivities of their phenology, which suggests that the two are causally related. Our results therefore indicate that both the phenologies of different taxonomic groups and the synchrony of plant-pollinator interactions are changing because of ongoing anthropogenic climate change.

### Caveats

When interpreting the results of our study, it is important to consider some caveats of the collections data and occurrence records we used. For instance, the spatial distribution of collections data is usually quite heterogeneous, and this was also true for our data (See diagnostic plots 7 & 8 in both **supplementary diagnostic plot documents**). We attempted to correct for spatial heterogeneity in our analyses by including latitude, longitude, and altitude in the statistical models. Moreover, our data are not only spatially but also temporally heterogenous. While the plant data are well-distributed across decades, the pollinator data are sparser during earlier decades, making slope estimation more sensitive to outliers, particularly for individual species. For entire taxonomic groups, however, these effects should cancel out each other.

Another problem is that not all of the recorded plants were in bloom at the time of recording or not all insects were adults. Particularly in the GBIF category “human observation” a fraction of records is of plants in a vegetative state. In a preliminary inspection of a subset of 23 early-flowering species we found only about 75% of the records with images to be of flowering species. Because of the large number of plant records, and the lack of images for many, it is impossible to evaluate this problem for our entire data set. However, we don’t expect any systematic trends in such ‘inactive’ specimens over time, so they should not have biased our analyses but rather increased the overall noise in our data.

The data we used allowed us to estimate average phenology/activity, but it did not allow to disentangle different aspects of phenology such as changes in first activity versus peak activty, or the duration of phenology. Some of these aspects might change even if peak activity remains unchanged, and it is possible that we have missed some dimensions of phenology variation, and their temporal changes.

### Phenological shifts over time

We found that both plants and some insect pollinator groups significantly advanced their phenology. The advances of plants in our data are stronger than those reported in most previous publications. For example, Fitter & Fitter [40] compared the first flowering dates of hundreds of plant species in England between 1954-1990 and 1991-2000, and they found an average advancement of −4.5 ± 0.8 (mean ± 1.96 * SE / 95% CI, same on all errors given below) days. A more recent long-term analysis by CaraDonna et al. [41] of phenology changes in subalpine meadow plants in the Rocky Mountains found an average advancement of peak flowering of 2.5 ± 0.2 days per decade from 1974 to 2012 for individual plant species, and 5.3 ± 1.7 days and 3.3 ± 1.6 days advancement of the spring and summer peak floral abundances of the entire plant community. All of these estimates are lower than the −7.2 ± 0.4 days per decade we found over the period from 1980 to 2020. Thus, our data seem to indicate that plants in Germany are shifting their phenology more strongly than in many other regions.

For insects, previous studies are less consistent, with widespread but not universal advances in spring phenology (mostly associated with warming) over the last decades [42]. For instance, a long-term study of butterflies in California showed that their peak flight times advanced on average by −1.7 ± 1 days per decade [43], a weaker trend than what we estimated in our data from Germany (−3.9 ± 0.6 days per decade). For bees, our data suggest an average shift of 2.6 ± 4 days per decade; however, the large confidence interval makes comparisons difficult. In New York, Bartomeus et al. [44] found that bee phenology advanced by −1.9 ± 0.1 days per decade during 1965-2011. This less than what we found in our data, but the discrepancy could be explained by the overall later time period of our study. For flies, there is little previous data on phenology shifts [45], except for Olsen et al. [46] who found an advancement of −6.2 days of the 10^th^ percentile of collection day (their measure of first flight) during 2000-2018. Although this appears similar to the −5.0 days per decade we found, a direct comparison is difficult here because of the different phenology measures. For the last of our studied insect groups, the beetles, we found no significant temporal shift of phenology at all, but also no previous studies except for such on individual pest species like bark beetles or potato beetles. Thus, we cannot judge how common the lack of temporal shifts is that we observed in our beetle data.

### Temperature sensitivities of plants and insects

At the level of individual species, there were strong correlations between the temporal shifts of species phenologies and their temperature sensitivities, i.e., how phenology was associated with interannual temperature variation. This was true for all taxonomic groups, and it strongly suggests a causal link between phenology shifts and climate change, supporting previous studies such as CaraDonna et al. [41] and Song et al. [47]. The taxonomic groups differed in their average temperature sensitivities, and these differences did not completely match the ones observed in temporal shifts. While plants generally tended to be the most temperature-sensitive group, there were also significant, albeit more moderate, temperature sensitivities in all pollinator groups except for the bees. Previous studies on insect phenology in the temperate zone (reviewed in [42]) have shown that increased spring temperatures are indeed often associated with earlier insect emergence, but that this pattern cannot be generalized as easily as for plants, as temperature–phenology relationships of insects are more complex. While many insects are able to plastically respond to warmer temperatures by speeding up their rates of development (and thus potentially emerge earlier), others have been found to not respond, or even to delay their phenology. This can be because insect development depends on other cues such as rainfall [48], because insects require a cold period during their diapause (if climate warming reduces this chilling period, this may even increase the amount of warming required for subsequent emergence; [42]), or because hibernation states are not temperature-sensitive. Fründ et al. [49] showed that bees overwintering in larval stages responded to higher winter temperatures with delayed emergence, while bees overwintering as adults showed advanced emergence (but had greater weight losses during overwintering).

There were some species with negative temperature sensitivities, i.e., delayed phenologies, in our data. This also connects well to some of the findings reviewed by Forrest [50], for instance that during the winter aboveground-nesting bees experience different temperatures than the plants they feed on during the summer. Such microclimate differences between overwintering insects could sometimes explain contrasting climate responses. Furthermore, warming can change the number of generations per year (voltinism; [42]). All the above-mentioned mechanisms can increase variation in the phenological responses of insects to climate warming and may explain why climate change is not always accompanied by phenological advances but might also cause delays – as we observed e.g., in some beetles.

### Changes in plant-pollinator synchrony

When we analysed the synchrony of predicted plant-pollinator interactions, we found clear trends in shifting synchrony, with different magnitudes in the pollinator groups. As the phenology of plants generally advanced faster than that of insects during the last decades, but plants had historically been the later partner in most plant-pollinator interactions, these shifts overall led to greater plant-pollinator synchrony. However, if the observed trends continue, then many of the studied interactions will soon reach absolute synchrony, and after that the interactions may become more asynchronous again, albeit in the opposite direction. For plant-hoverfly interactions this point has already been reached. If interactions will become more asynchronous again in the future, then the resilience of pollinator networks, in particular through pollinator generalism, could buffer some of the impacts of phenological mismatches [8]. However, while generalist pollinators make up the larger part of the interactions in most pollination networks, some plant-pollinator interactions are highly specialized, and these might be the ones suffering most from future mismatches [51]. One idea for future research could therefore be to focus specifically on specialist pollinators, or to compare long-term trends and climate change effects on generalist versus specialist plant-pollinator interactions.

## Supporting information

Supplementary plots

Supplementary diagnostic plots 1

Supplementary diagnostic plots 2

## Acknowledgements

We are grateful to two reviewers of a previous submission, whose thorough and constructive feedback greatly improved our manuscript. Our work has been supported by the DFG Priority Program 1374 “Infrastructure-Biodiversity-Exploratories” (DFG project BO 3241/7-1 to OB).

## Authors contributions

JF and FMW conceived the study, JF and FMW collected the data, JF analysed the data and wrote the first draft of the manuscript, with guidance from FMW. JFS and OB provided input to data analysis and manuscript writing. All authors read and approved the final manuscript.

## Data availability

The R code used to conduct the analysis is available at https://github.com/jonasfreimuth/Phenological-shifts-germany.

