## Supplementary plots for "Climate warming changes synchrony of plants and pollinators"

Supplementary Material - Temperature model diagnostics

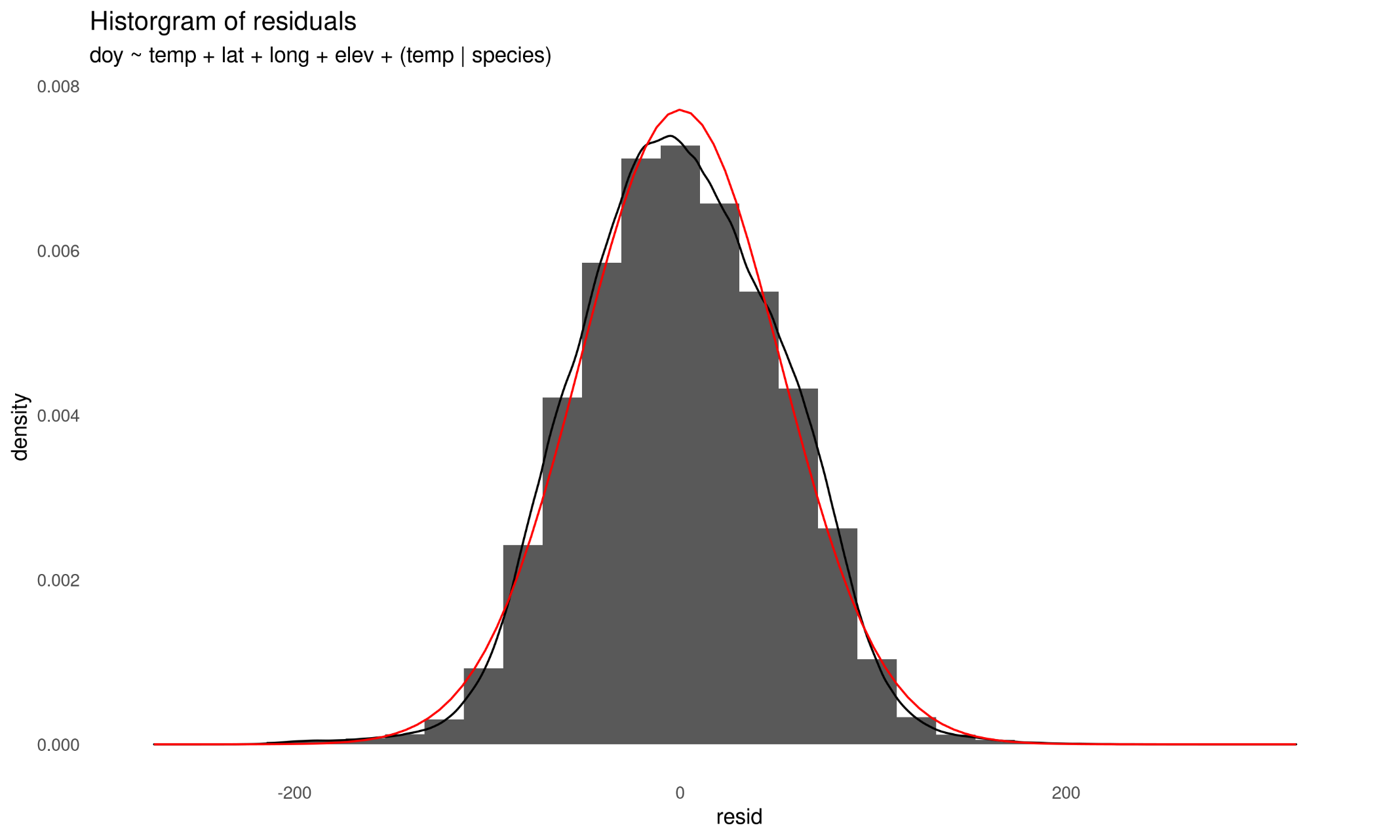

Diagnostic Plot 1 Distribution of residuals of the temperature model. Black line indicates density of residuals, red line indicates a perfect normal distribution.

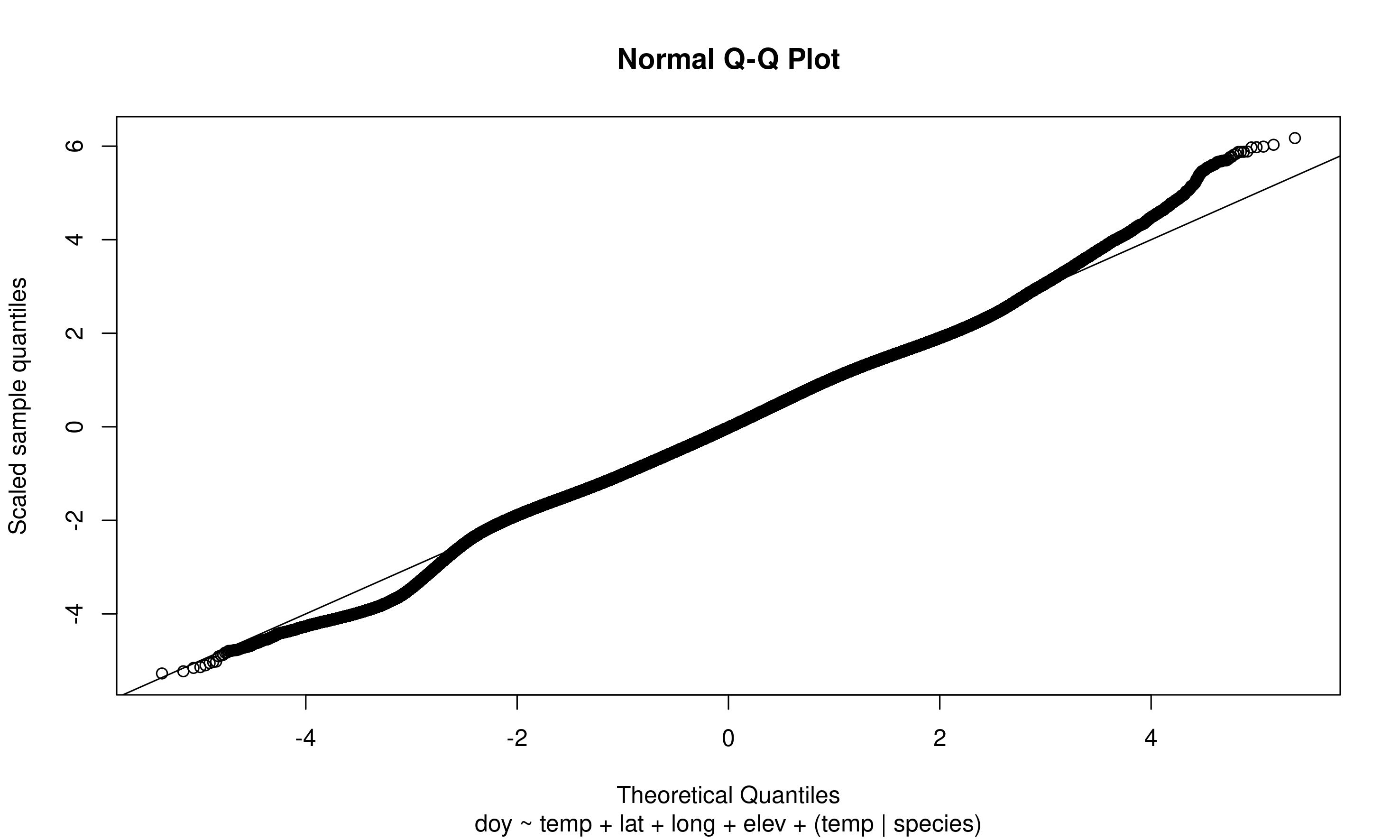

Diagnostic Plot 2 Quantile-Quantile plot of residuals of the temperature model against a normal distribution. Black line indicates perfect fit.

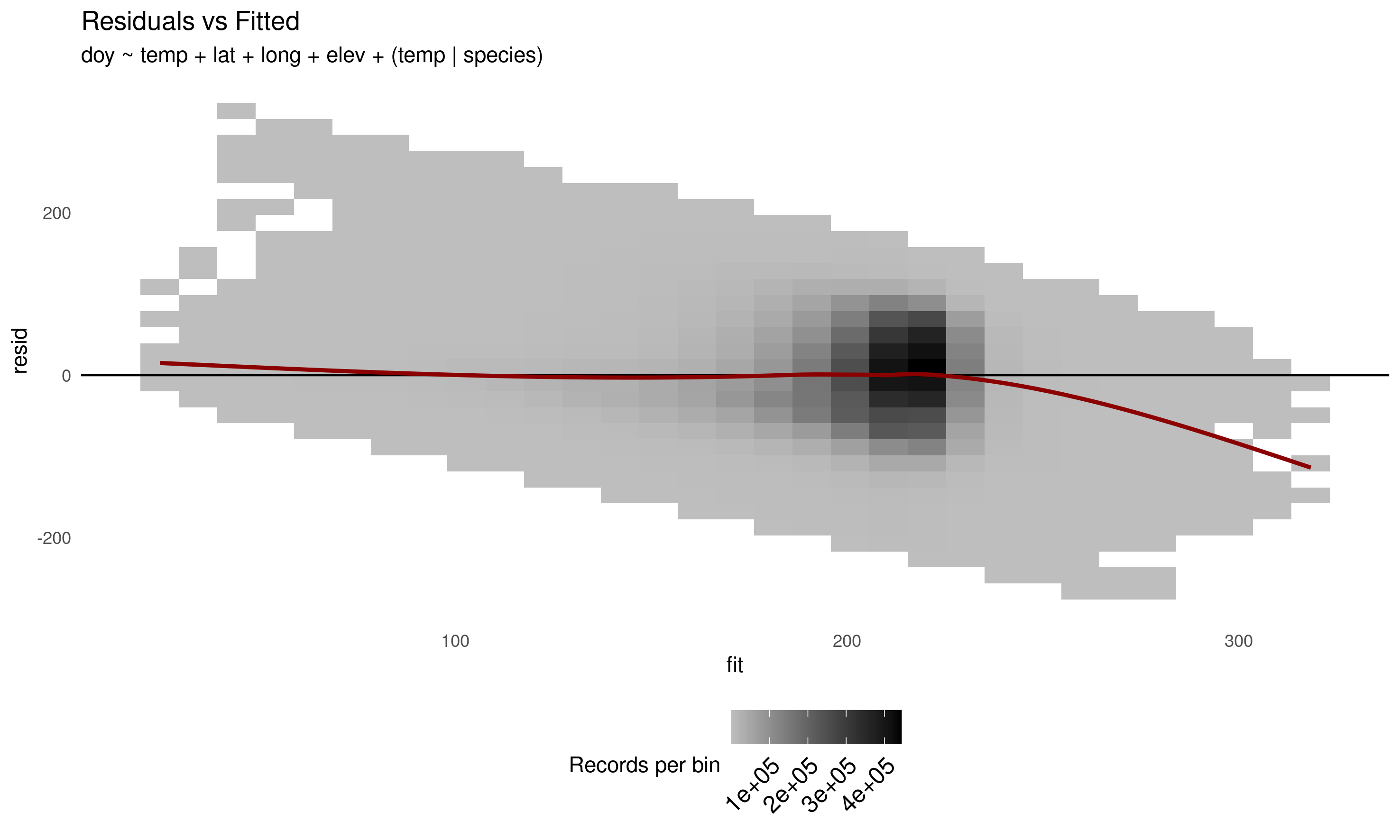

Diagnostic Plot 3 Overall density plot of temperature model residuals vs fitted values. The dark red line indicates a generalized additive model curve fit to the data points. Cell shading indicates point density.

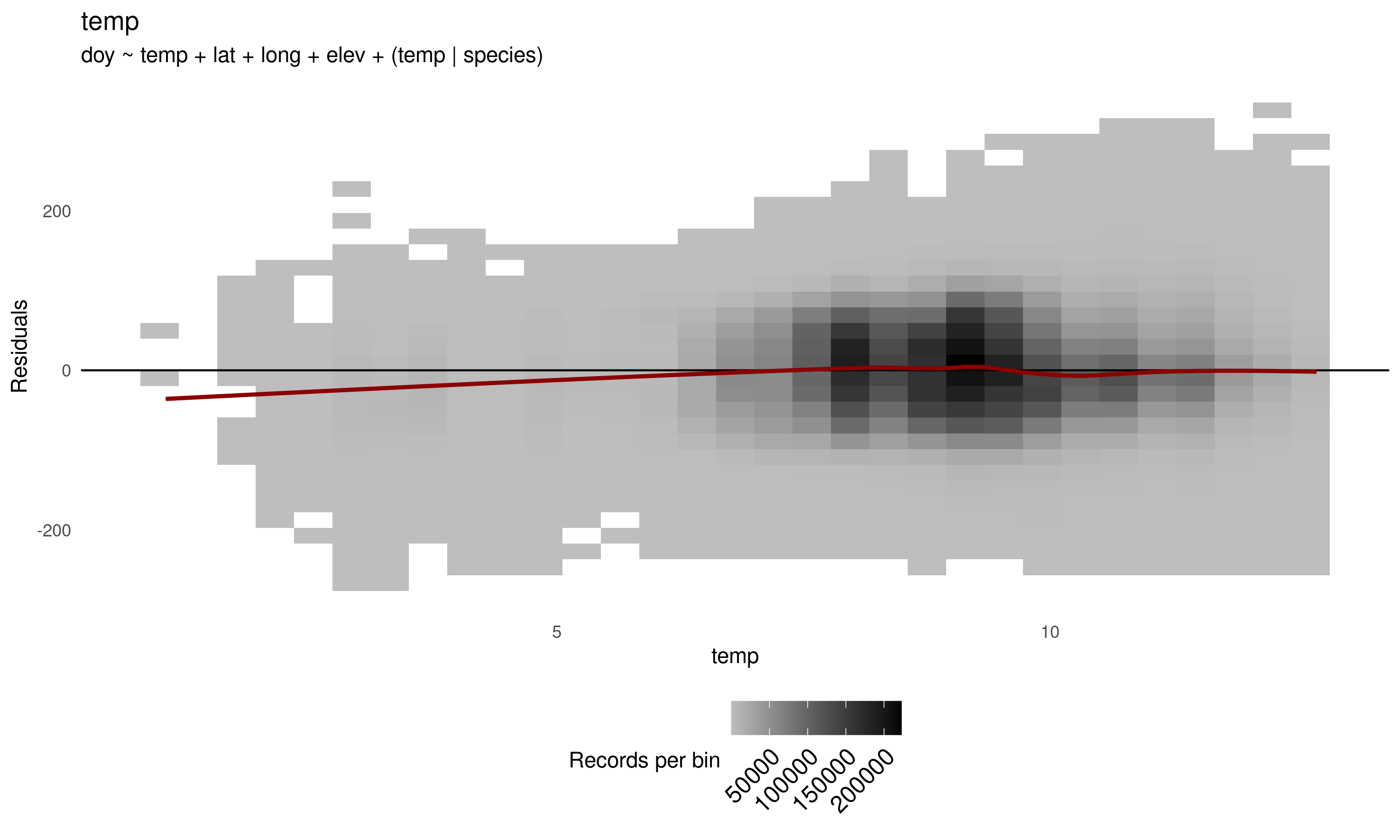

Diagnostic Plot 4 Overall density plot of temperature model residuals vs record yearly mean temperature values in degrees Celsius. The dark red line indicates a generalized additive model curve fit to the data points. Cell shading indicates point density.

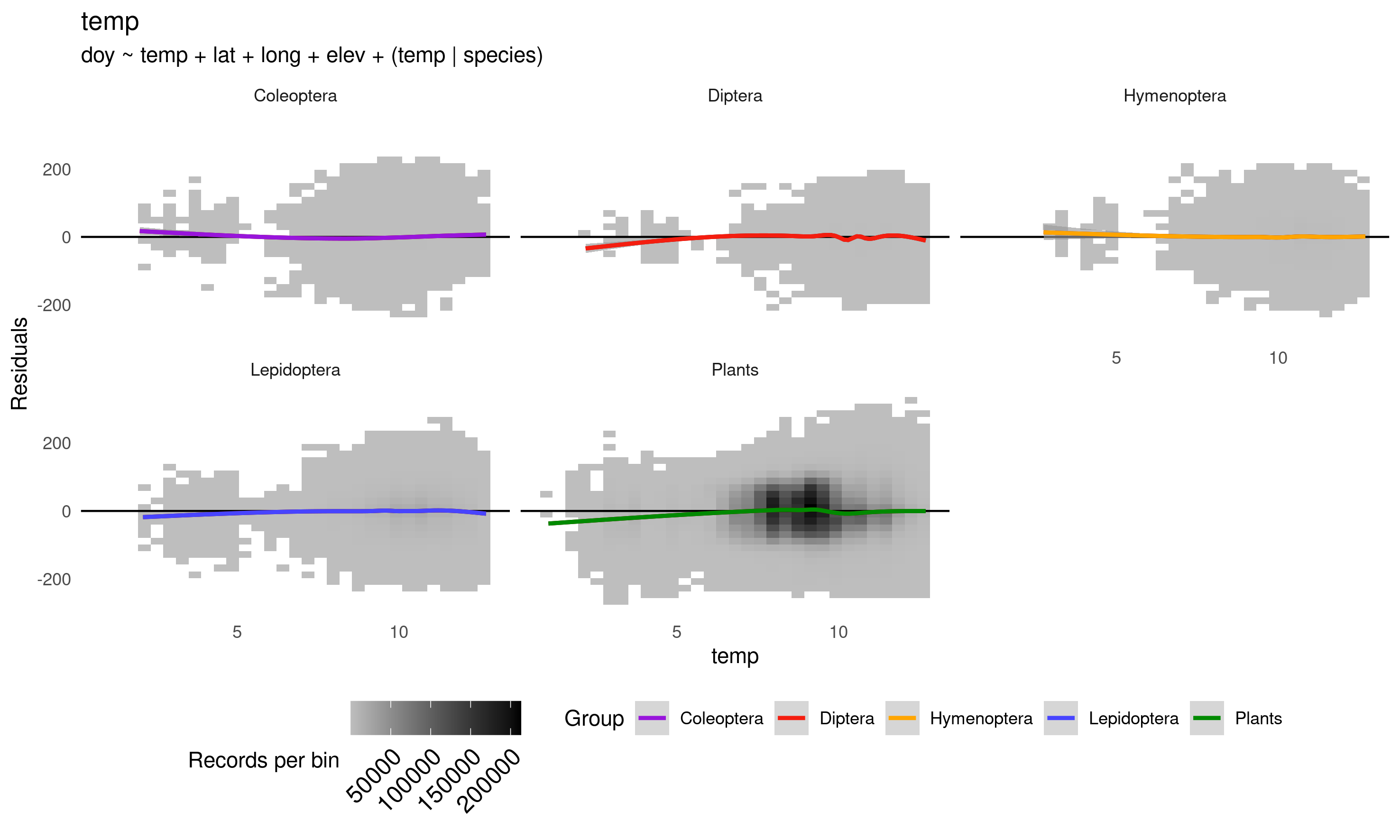

Diagnostic Plot 5 Density plot of temperature model residuals vs record yearly mean temperature values in degrees Celsius by taxonomic groups. The coloured lines indicate a generalized additive model curve fit to the data points. Cell shading indicates point density.

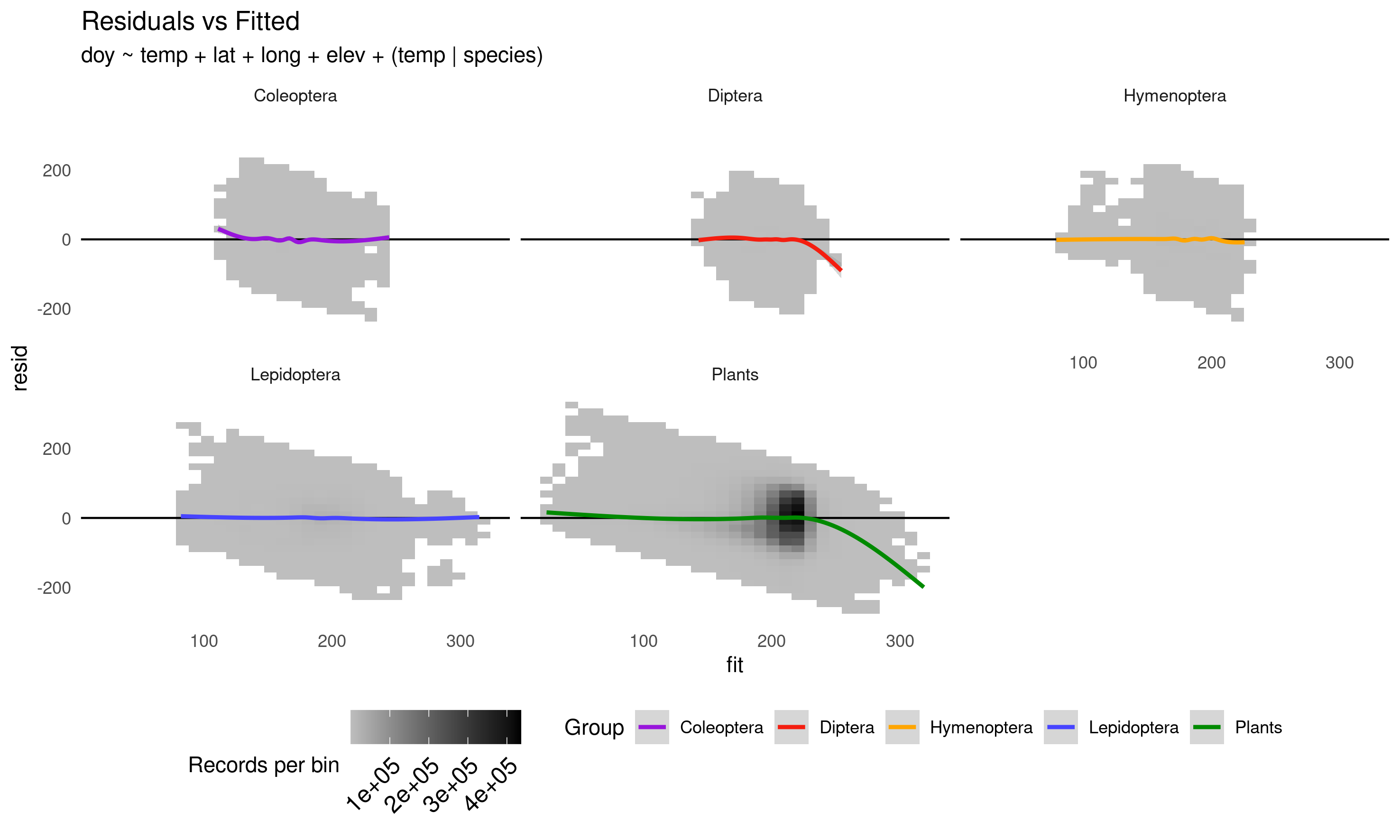

Diagnostic Plot 6 Density plot of temperature model residuals vs fitted values by taxonomic groups. The coloured lines indicate a generalized additive model curve fit to the data points. Cell shading indicates point density.

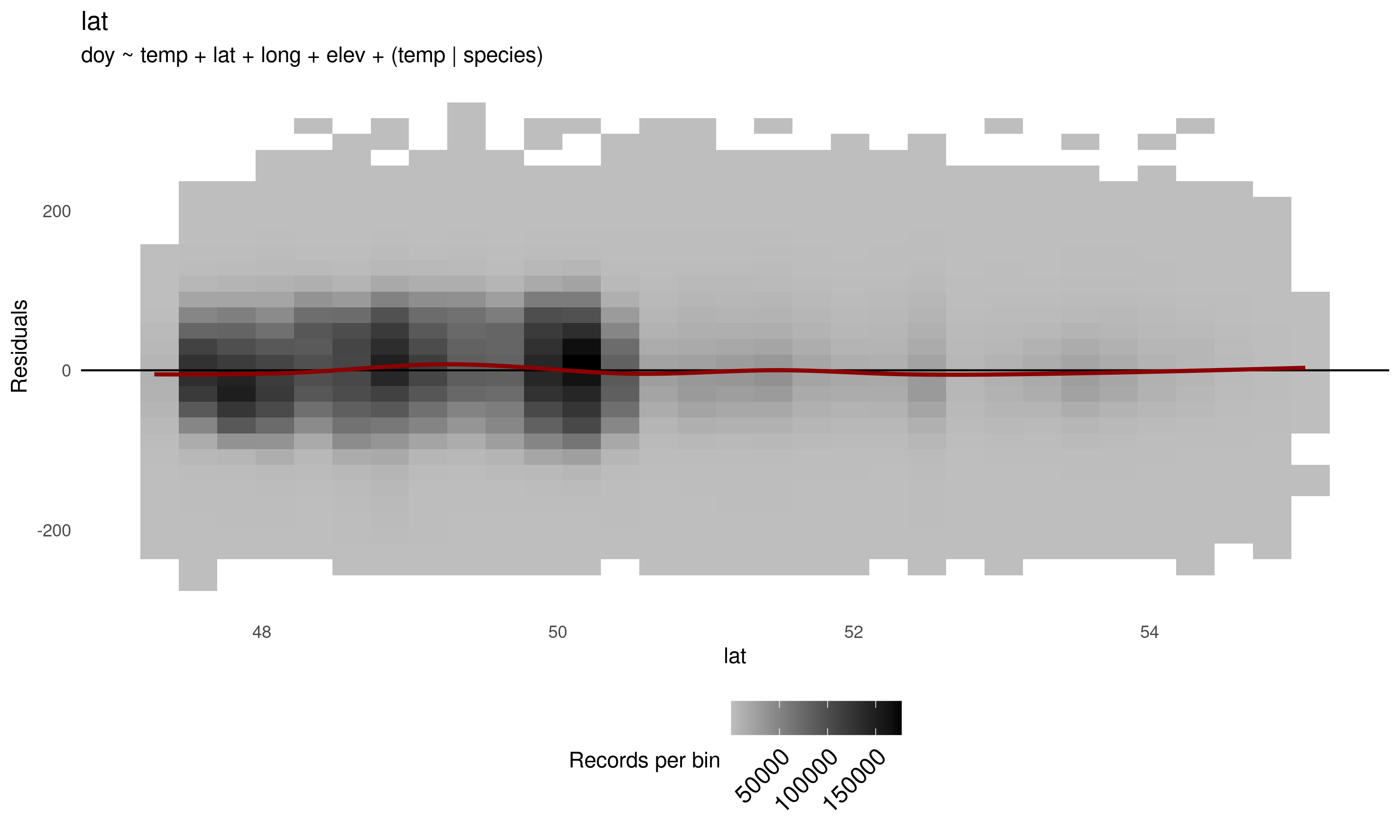

Diagnostic Plot 7 Overall density plot of temperature model residuals vs record latitude values in decimal degrees. The dark red line indicates a generalized additive model curve fit to the data points. Cell shading indicates point density.

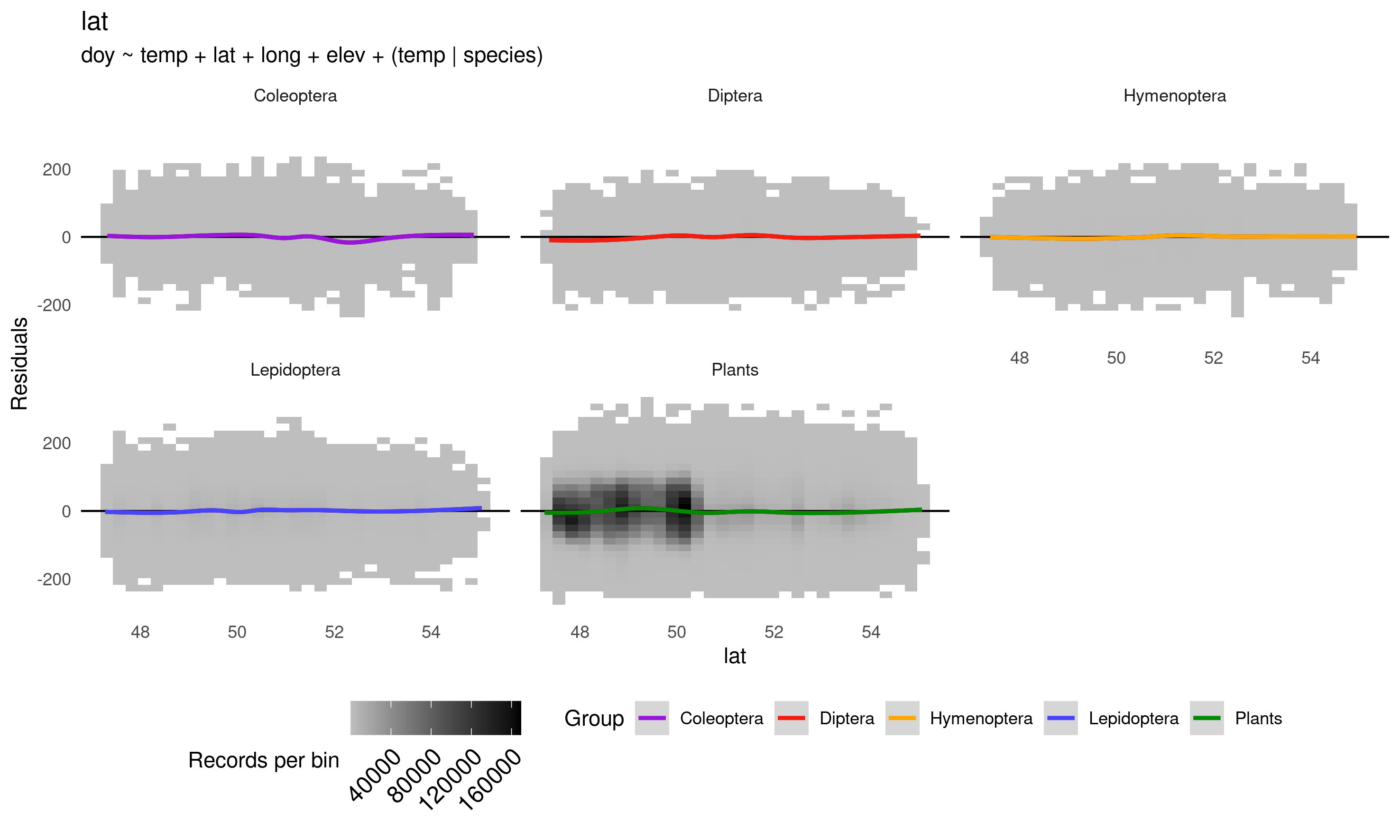

Diagnostic Plot 8 Density plot of temperature model residuals vs record latitude values in decimal degrees by taxonomic groups. The coloured lines indicate a generalized additive model curve fit to the data points. Cell shading indicates point density.

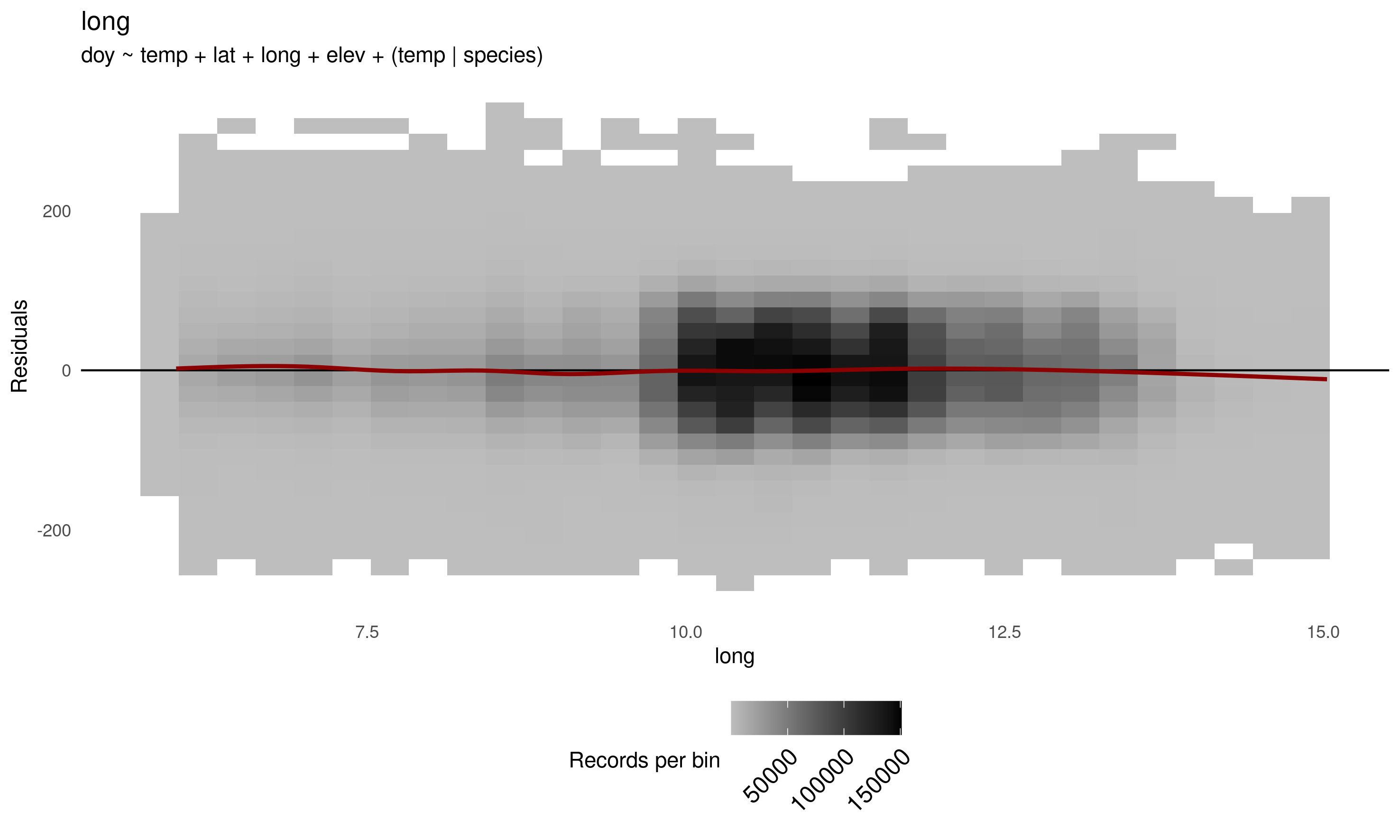

Diagnostic Plot 9 Overall density plot of temperature model residuals vs record longitude values in decimal degrees. The dark red line indicates a generalized additive model curve fit to the data points. Cell shading indicates point density.

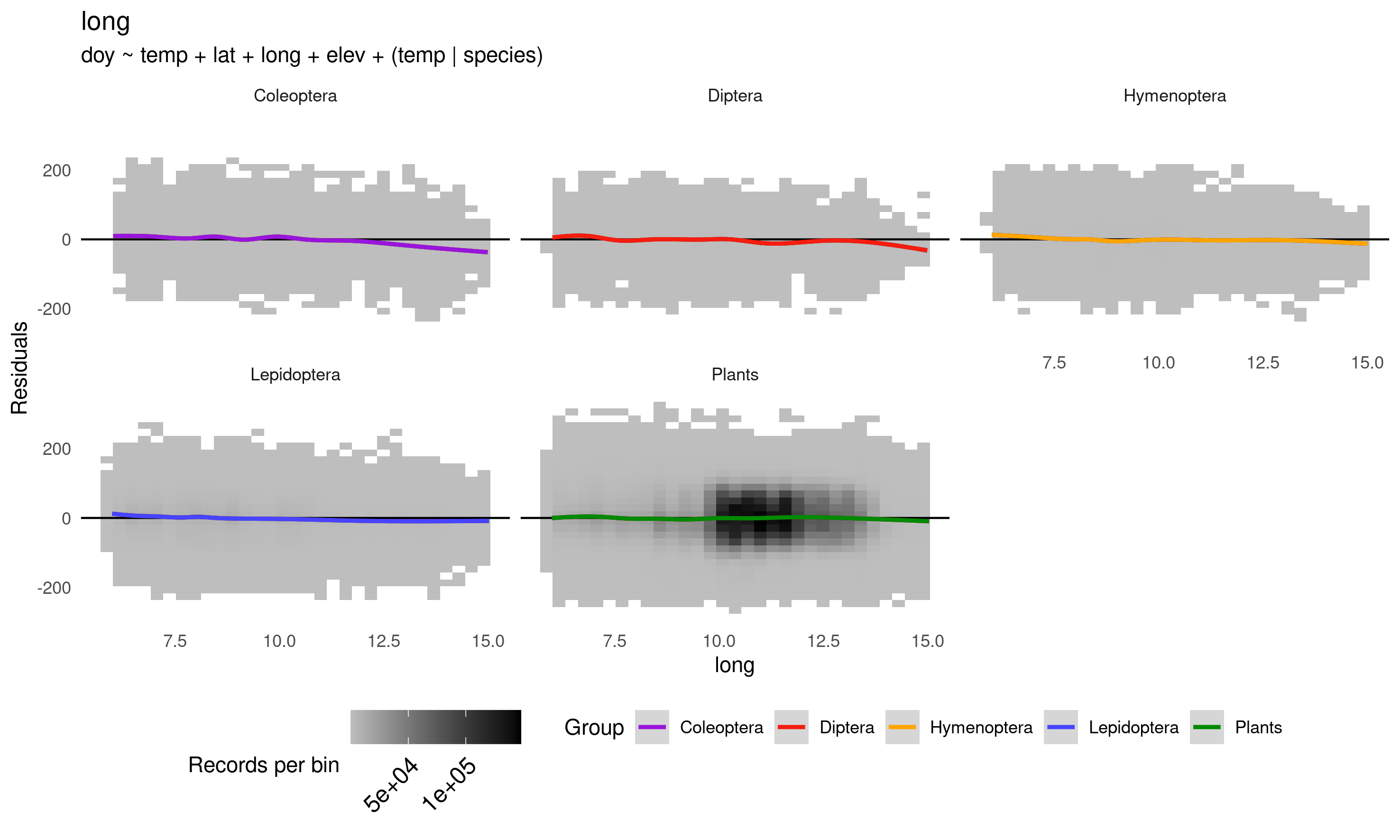

Diagnostic Plot 10 Density plot of temperature model residuals vs record longitude values in decimal degrees by taxonomic groups. The coloured lines indicate a generalized additive model curve fit to the data points. Cell shading indicates point density.

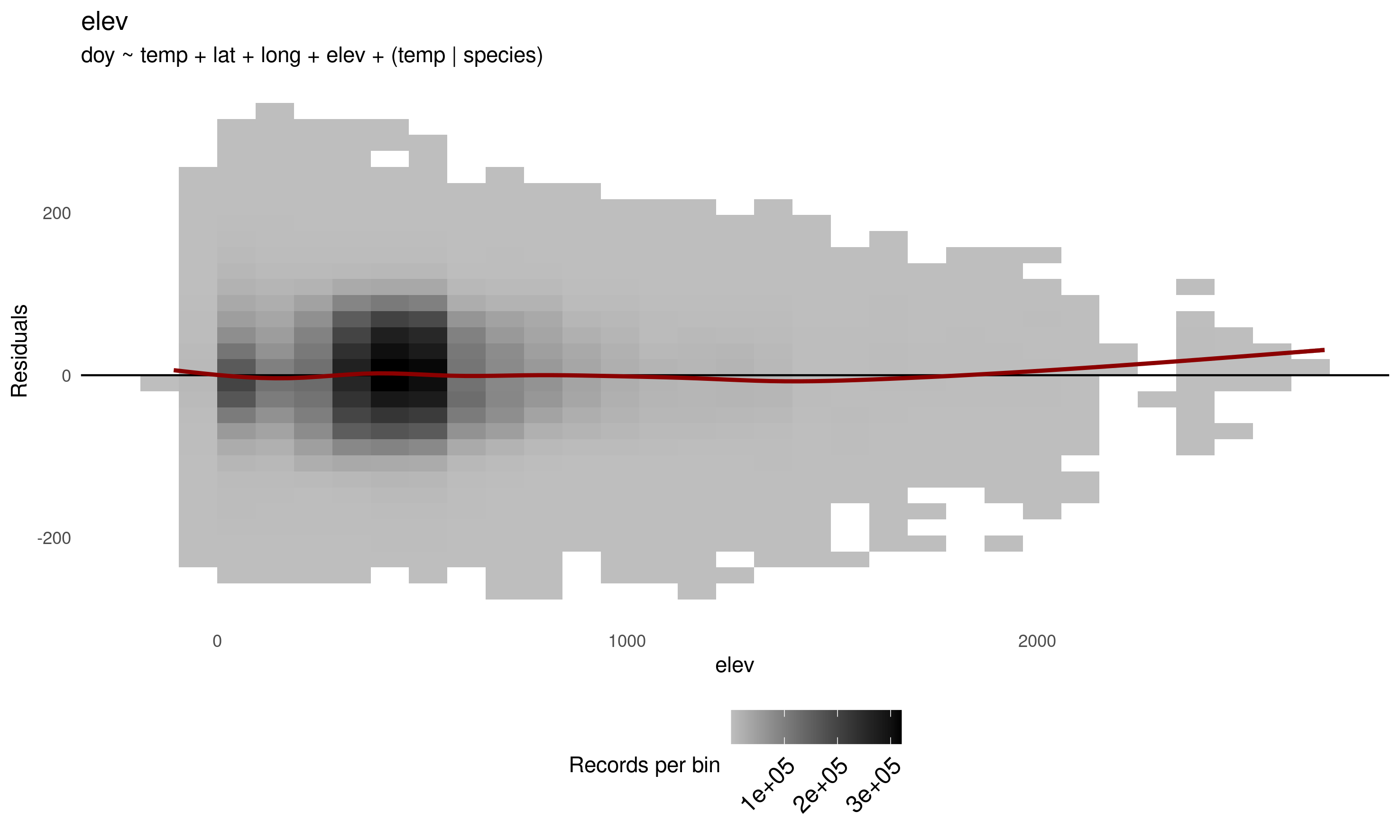

Diagnostic Plot 11 Overall density plot of temperature model residuals vs record elevation values in meters. The dark red line indicates a generalized additive model curve fit to the data points. Cell shading indicates point density.

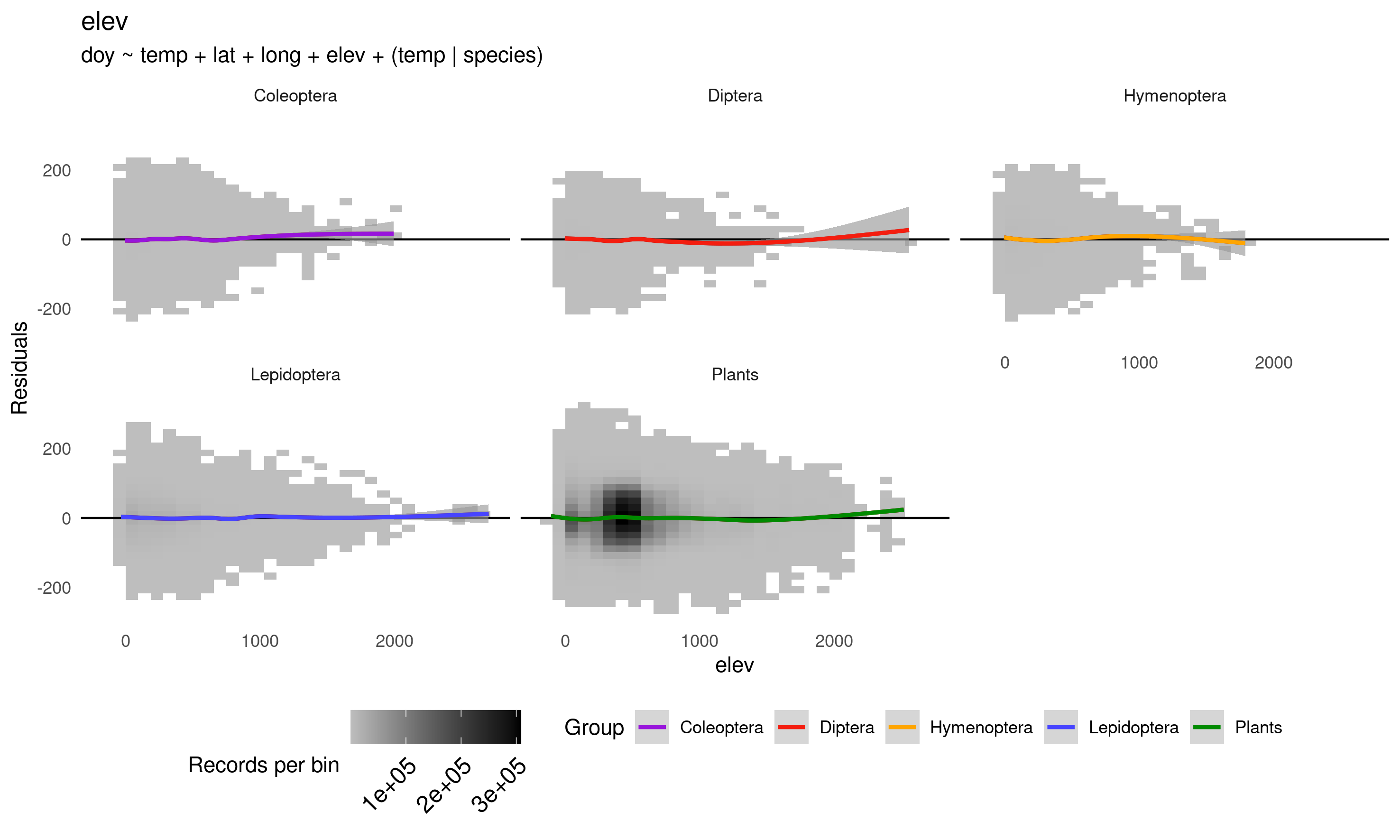

Diagnostic Plot 12 Density plot of temperature model residuals vs record elevation values in meters by taxonomic groups. The coloured lines indicate a generalized additive model curve fit to the data points. Cell shading indicates point density.

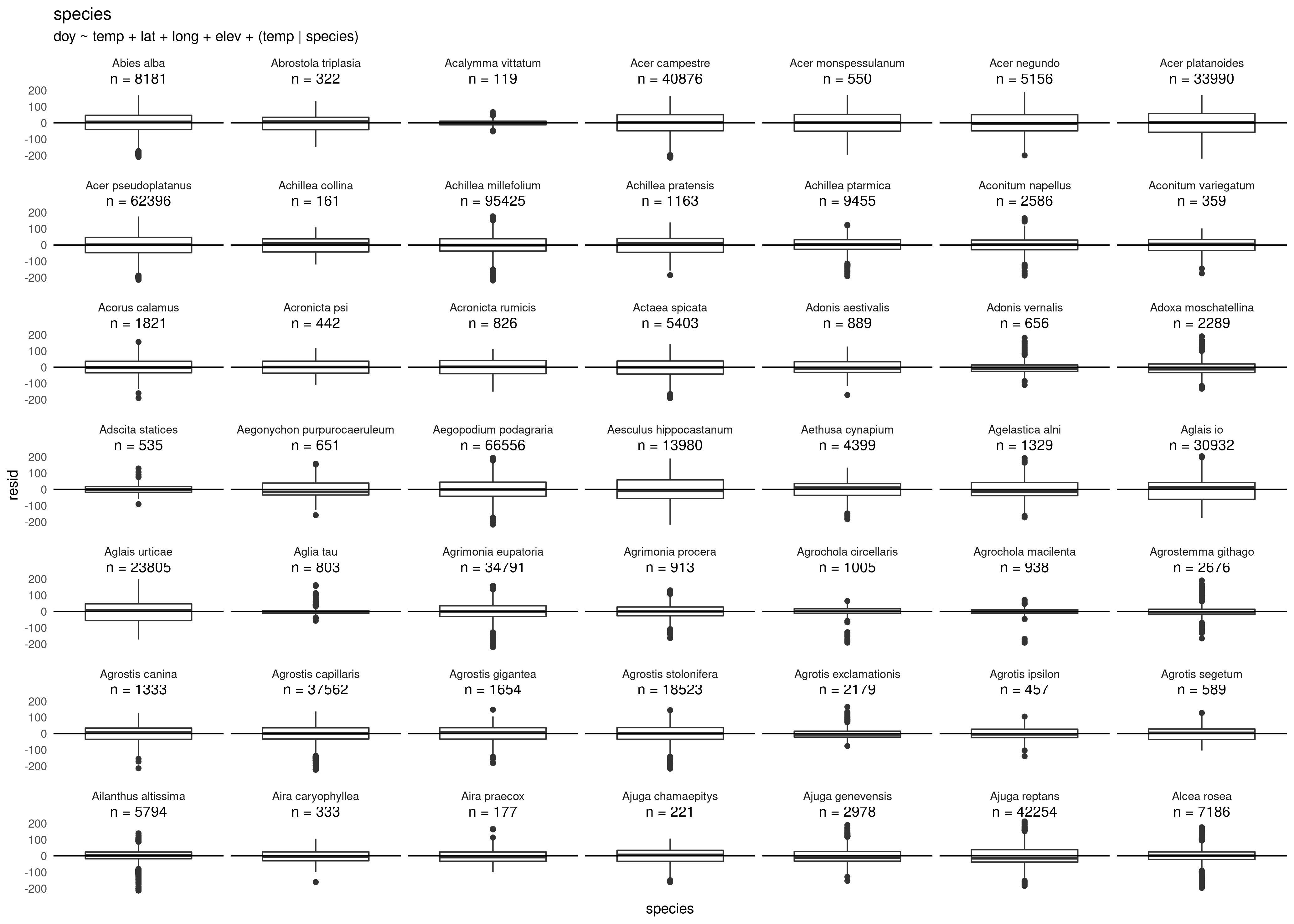

Diagnostic Plot 13 (and following) Boxplots of temperature model residuals by species with species numbers of records.

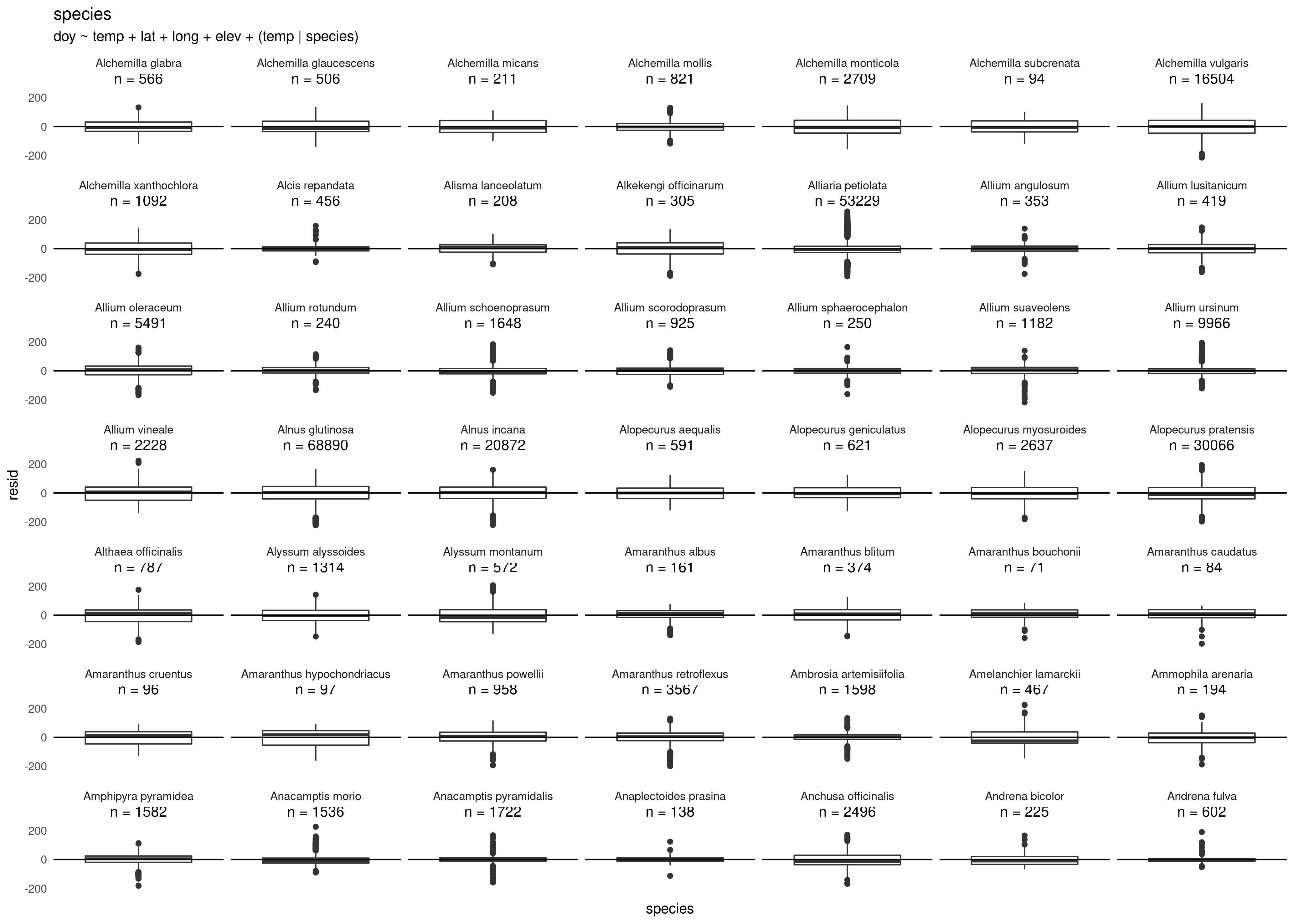

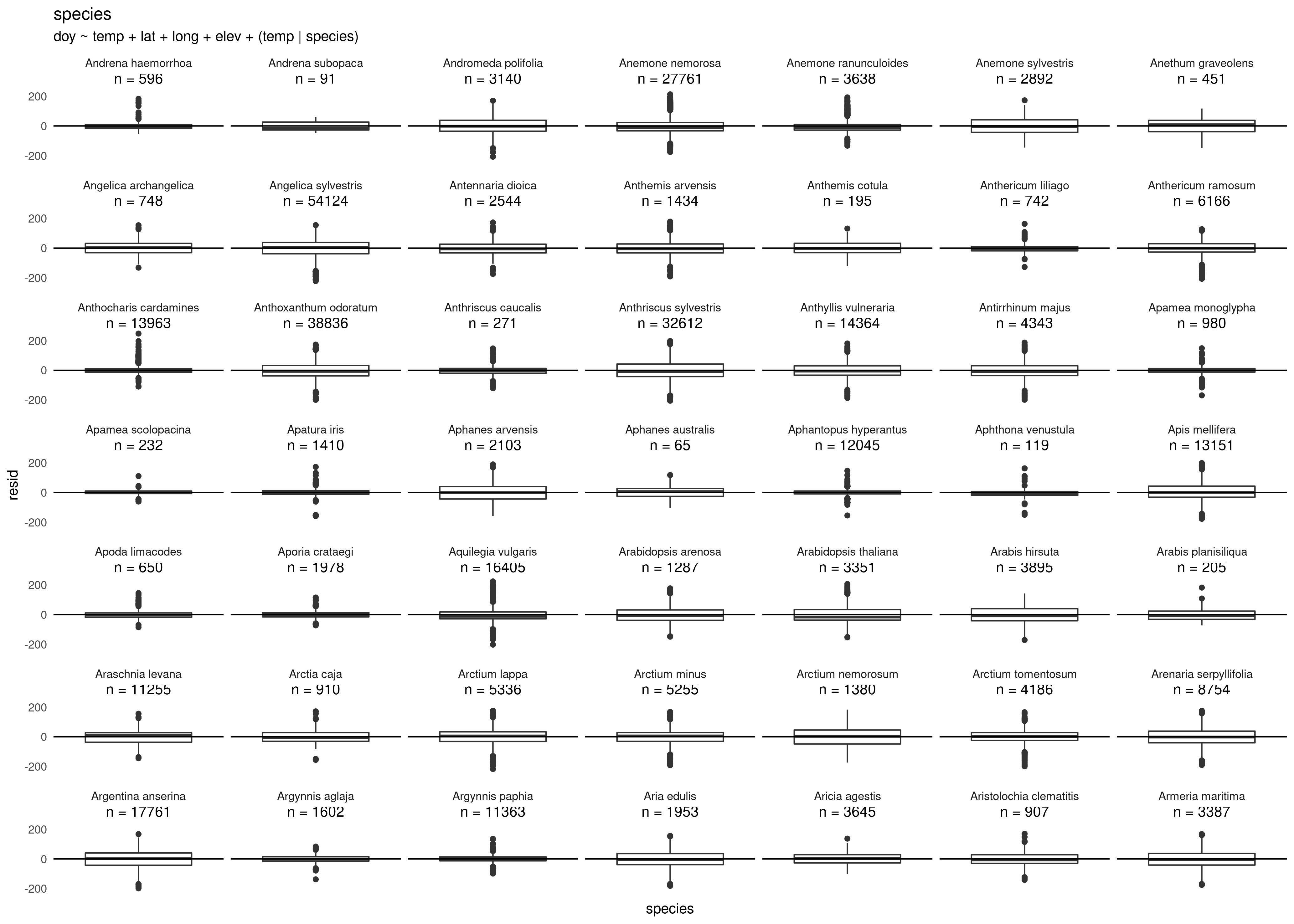

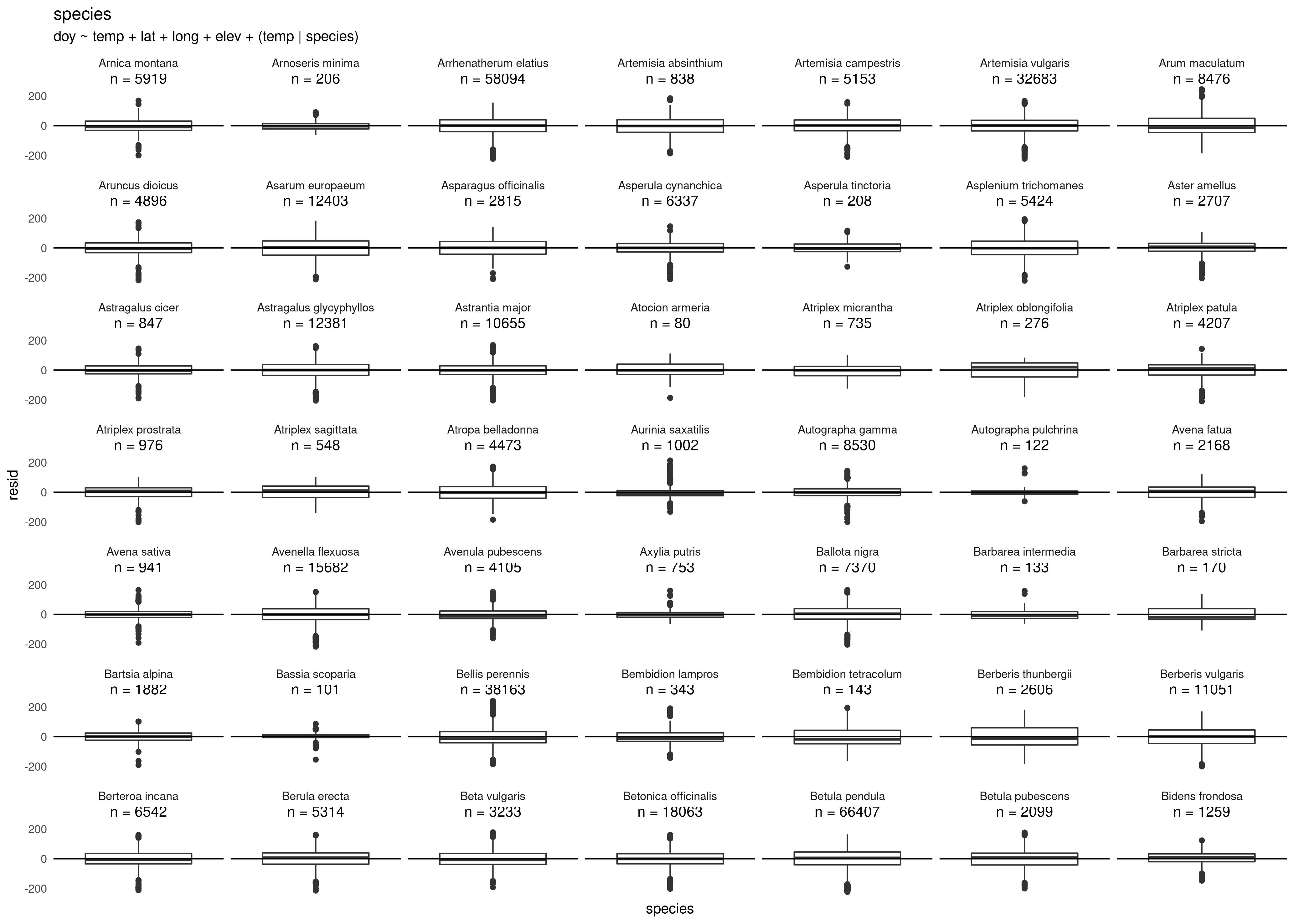

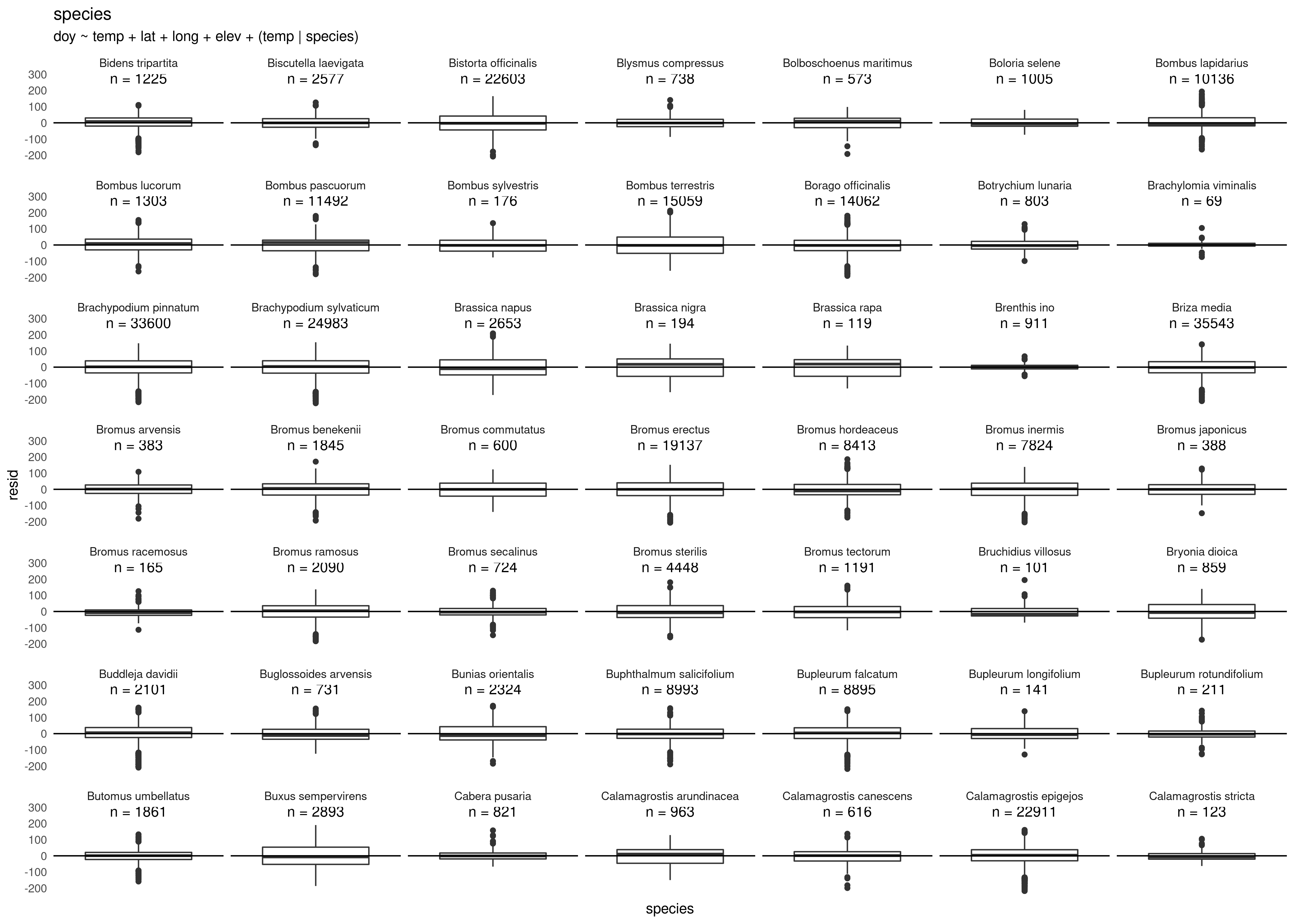

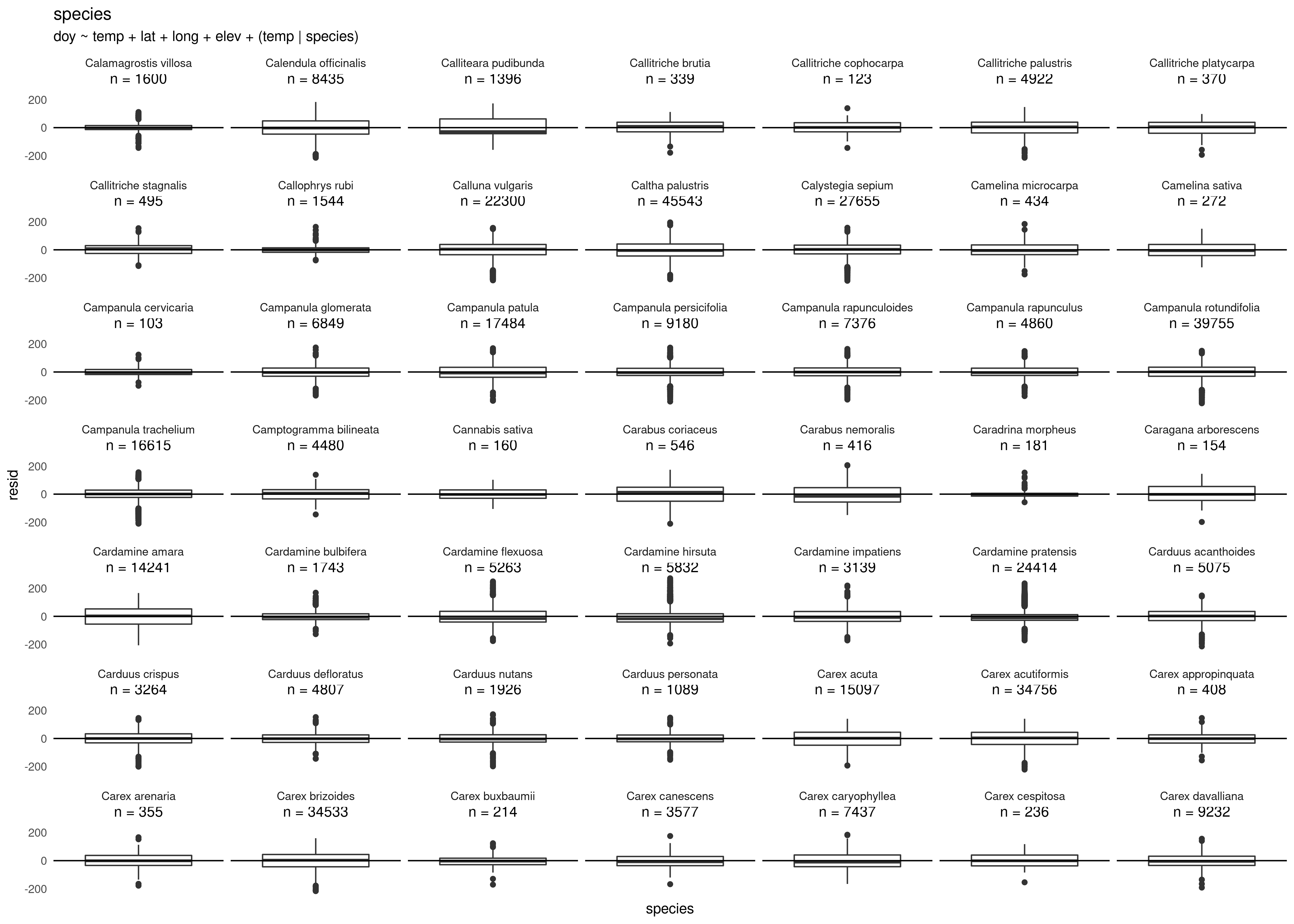

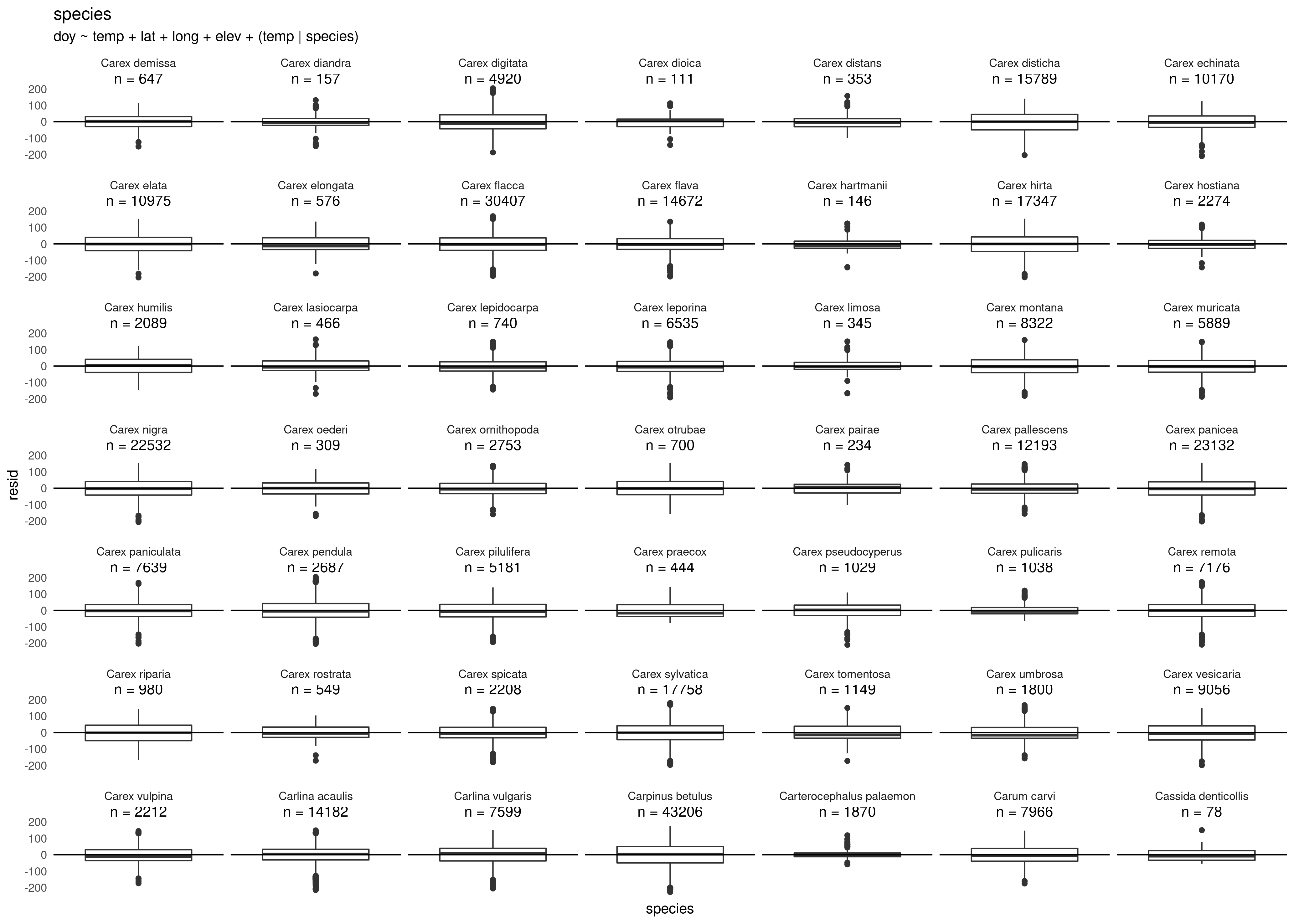

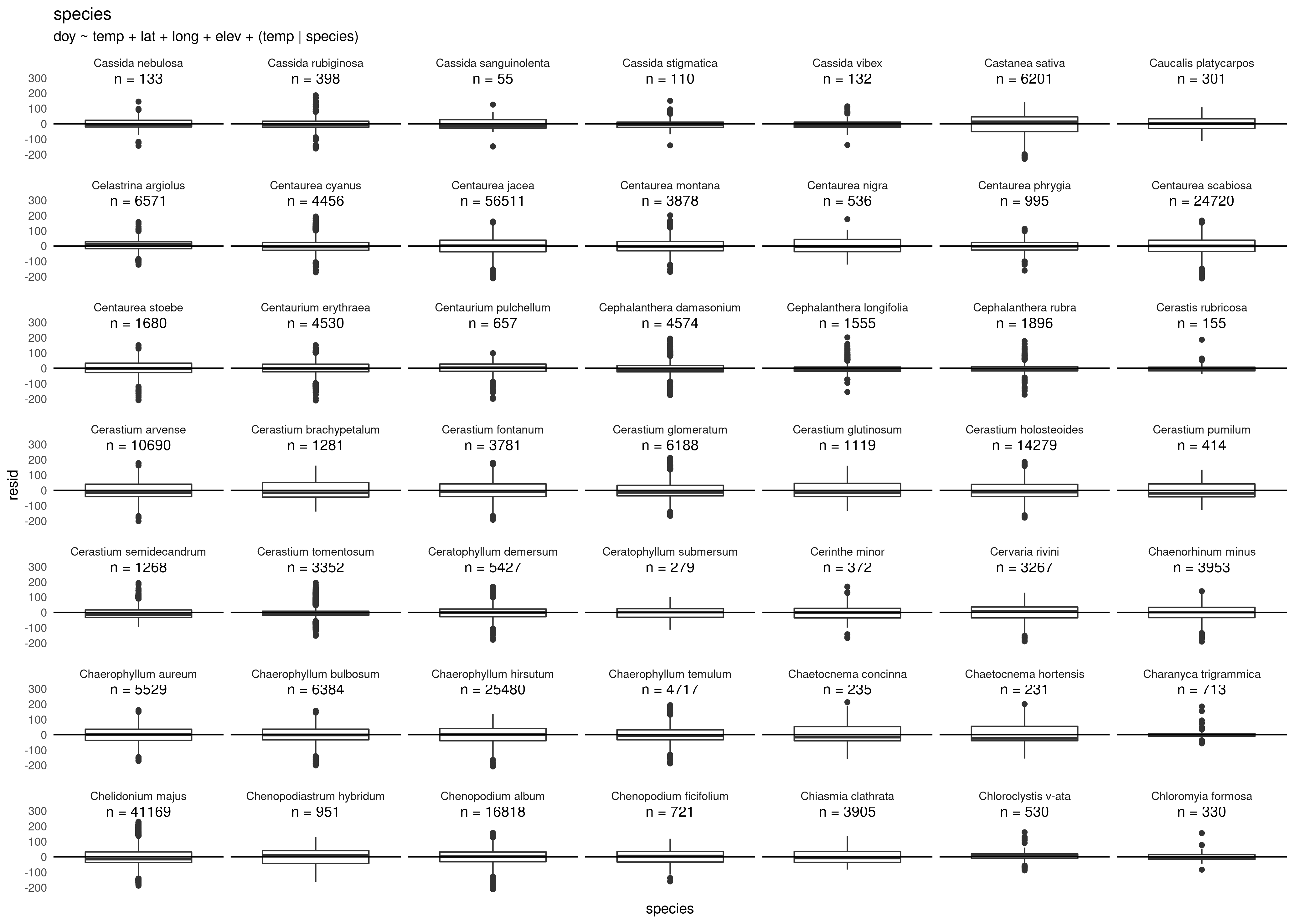

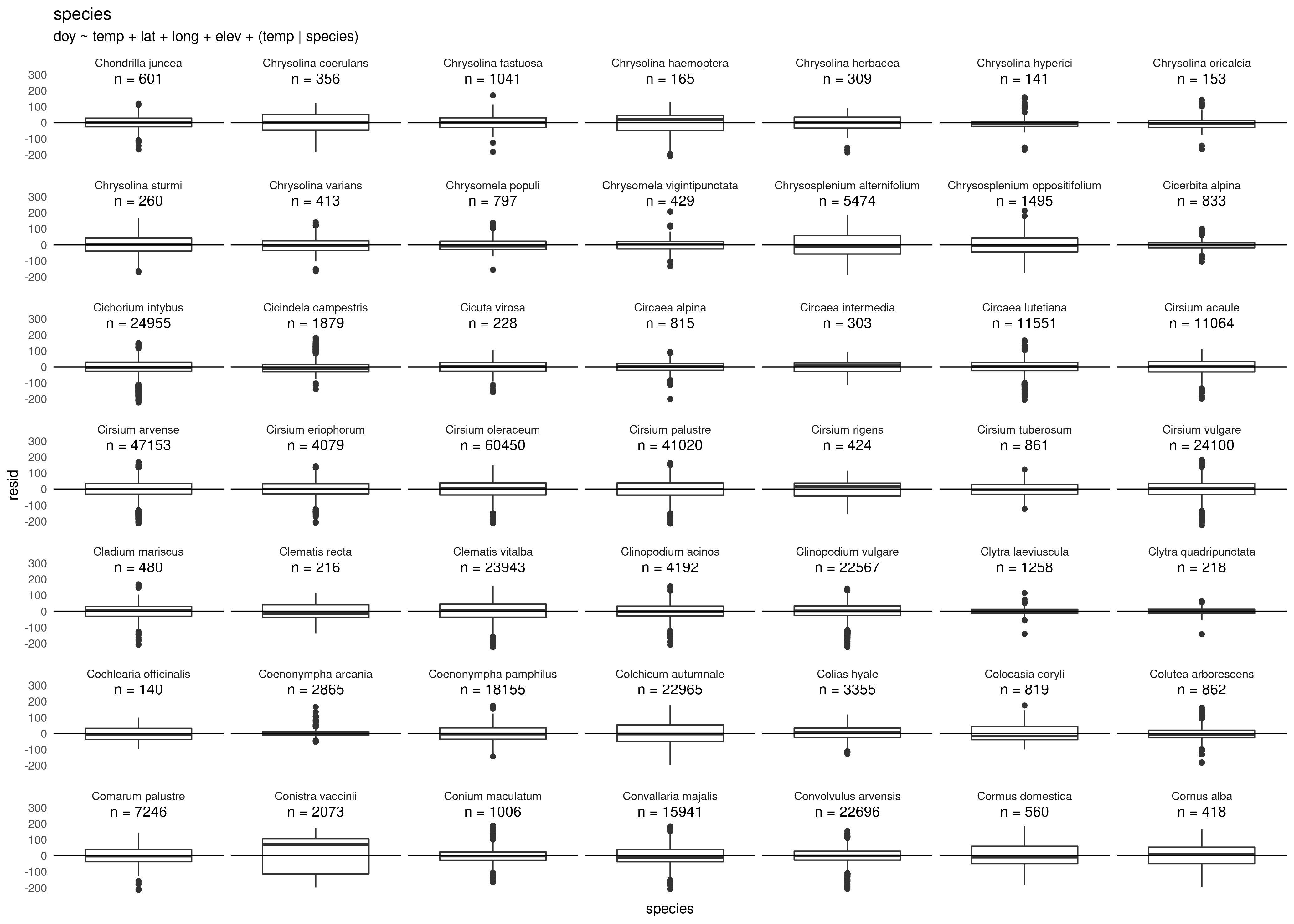

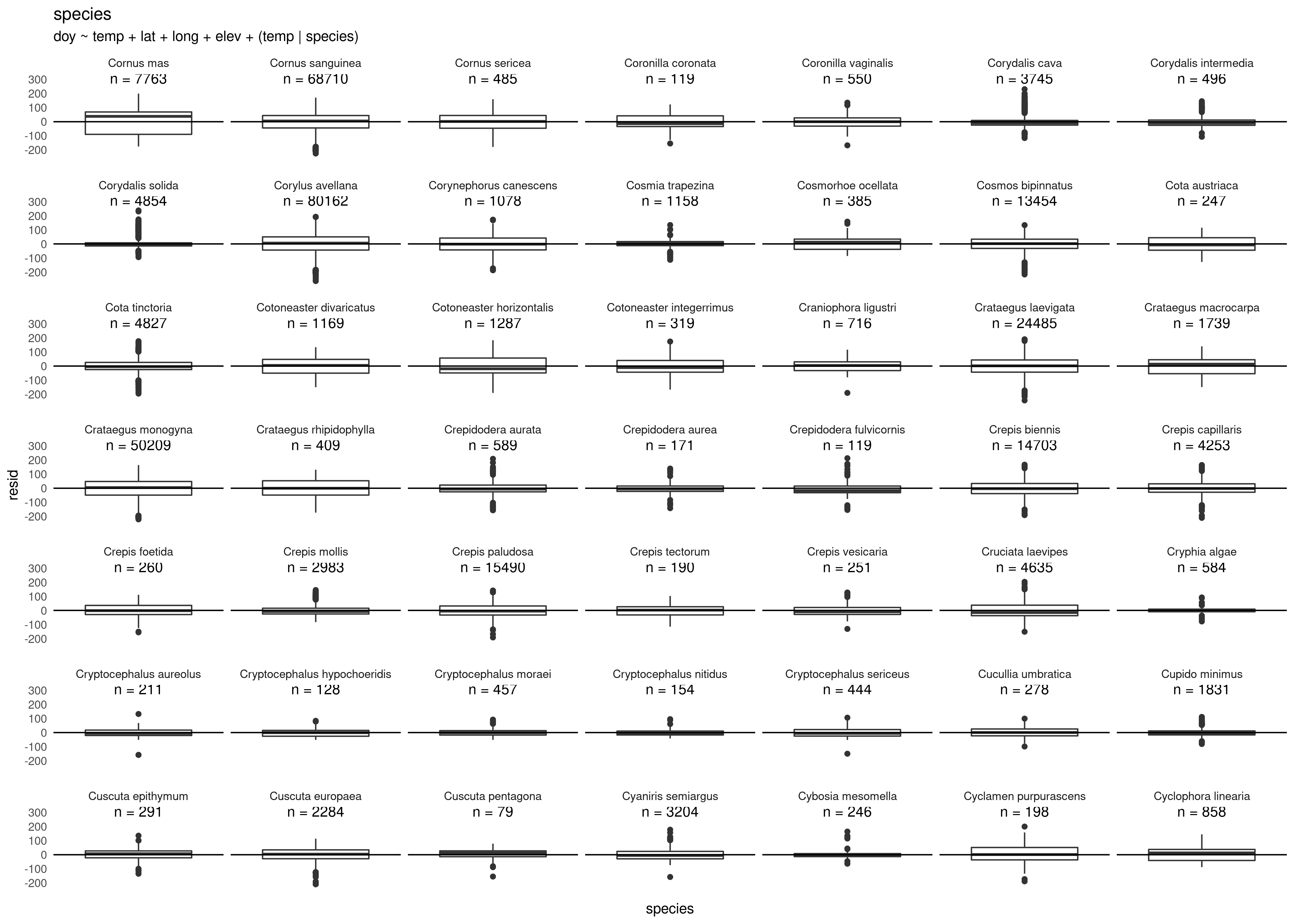

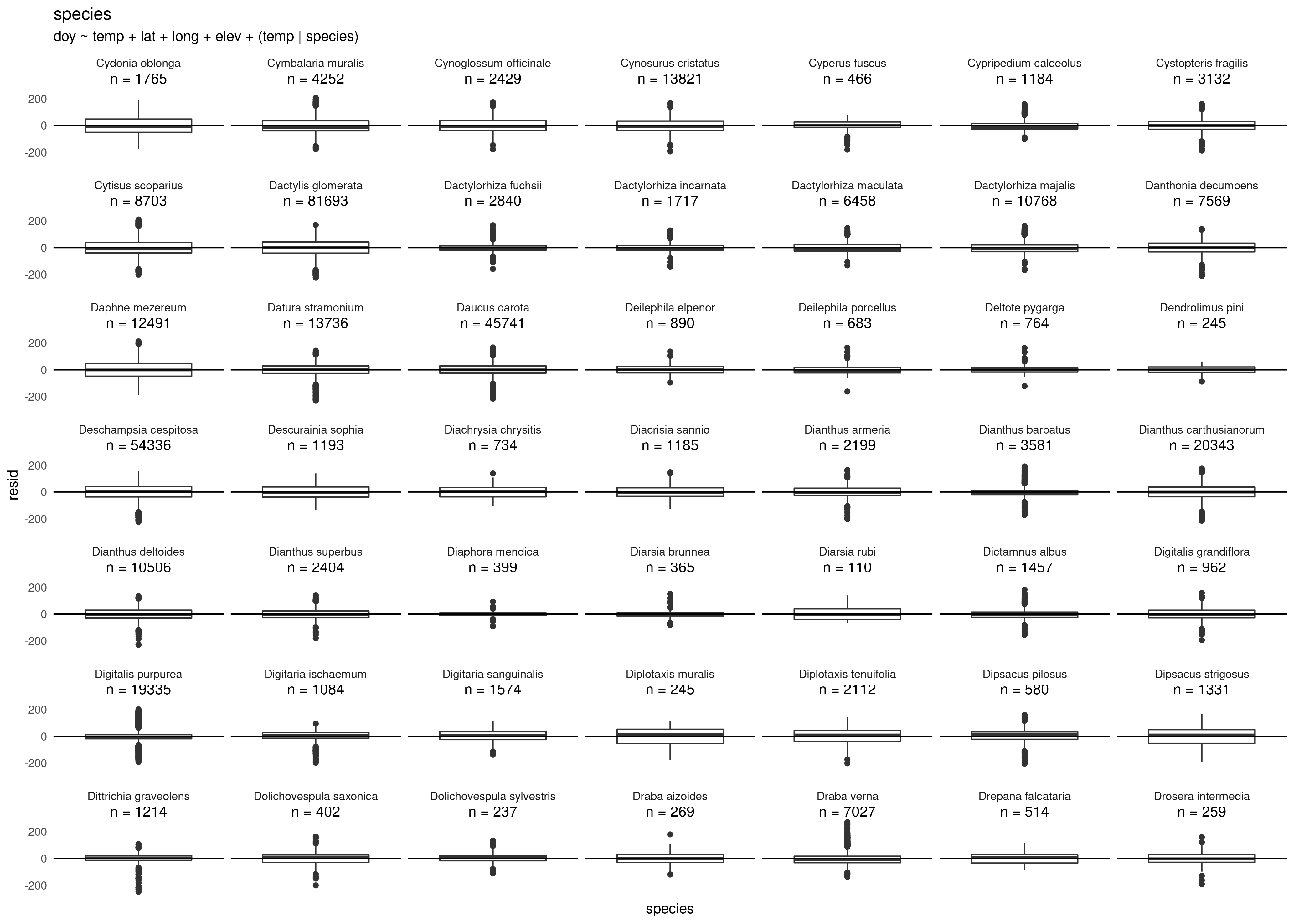

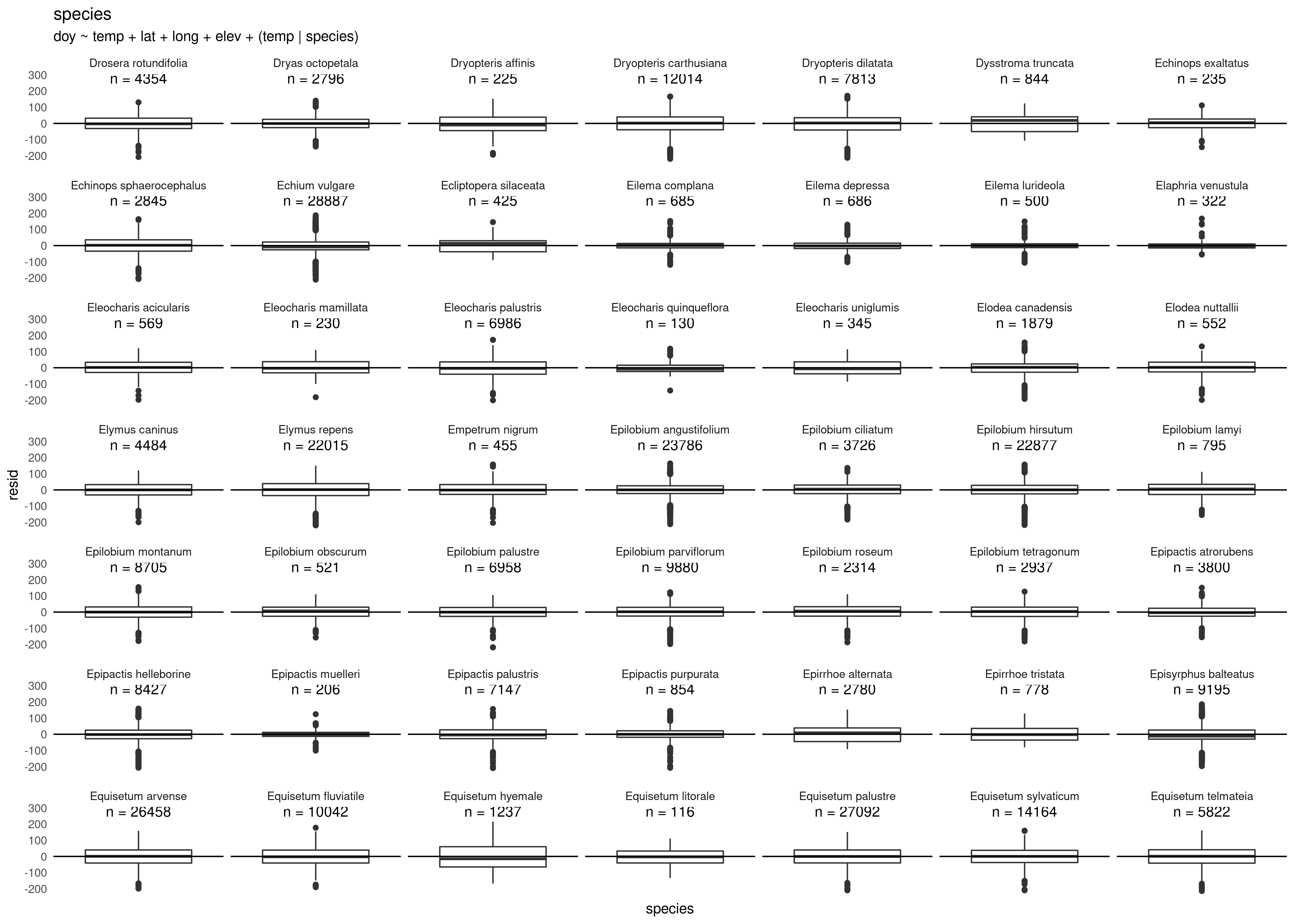

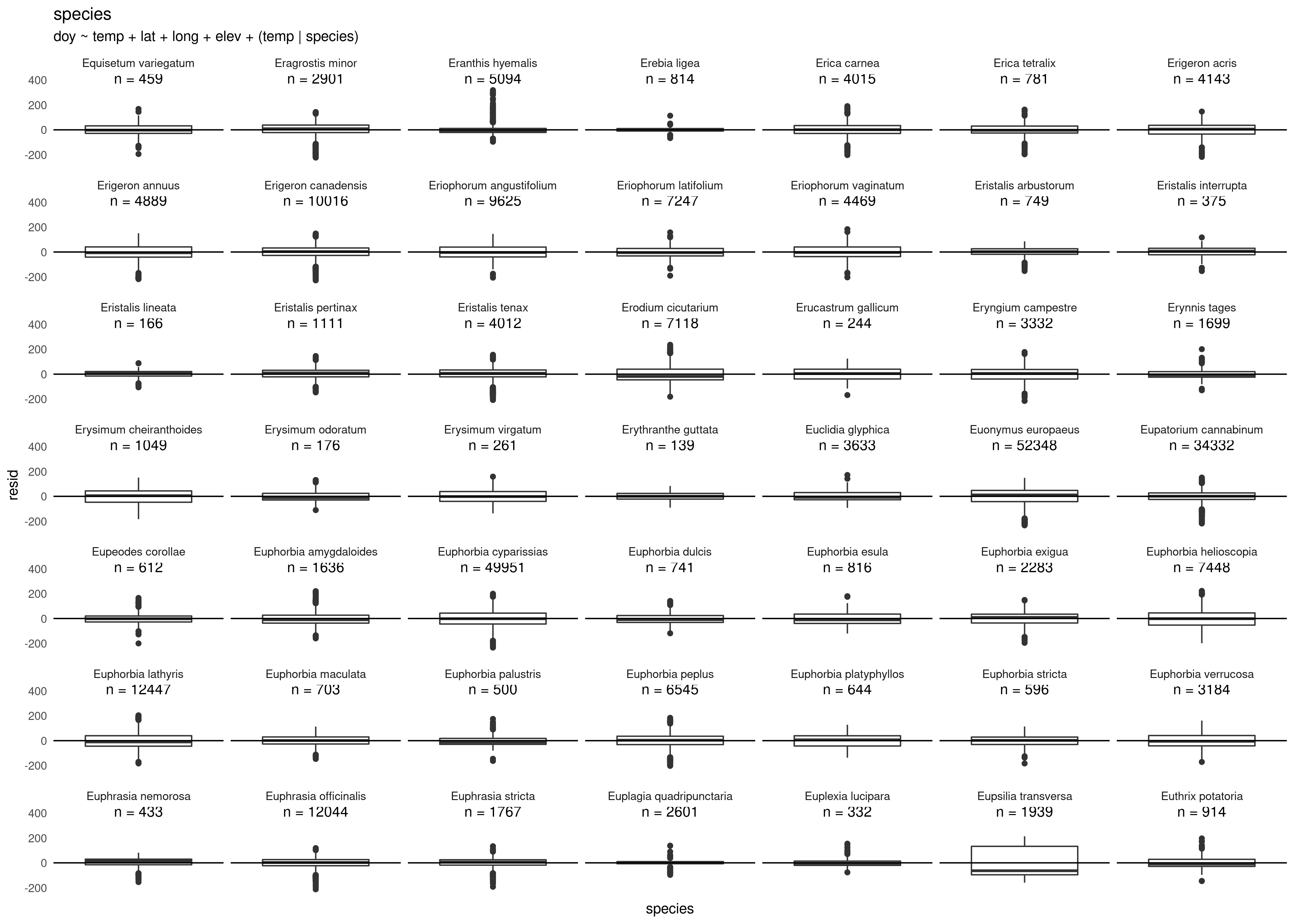

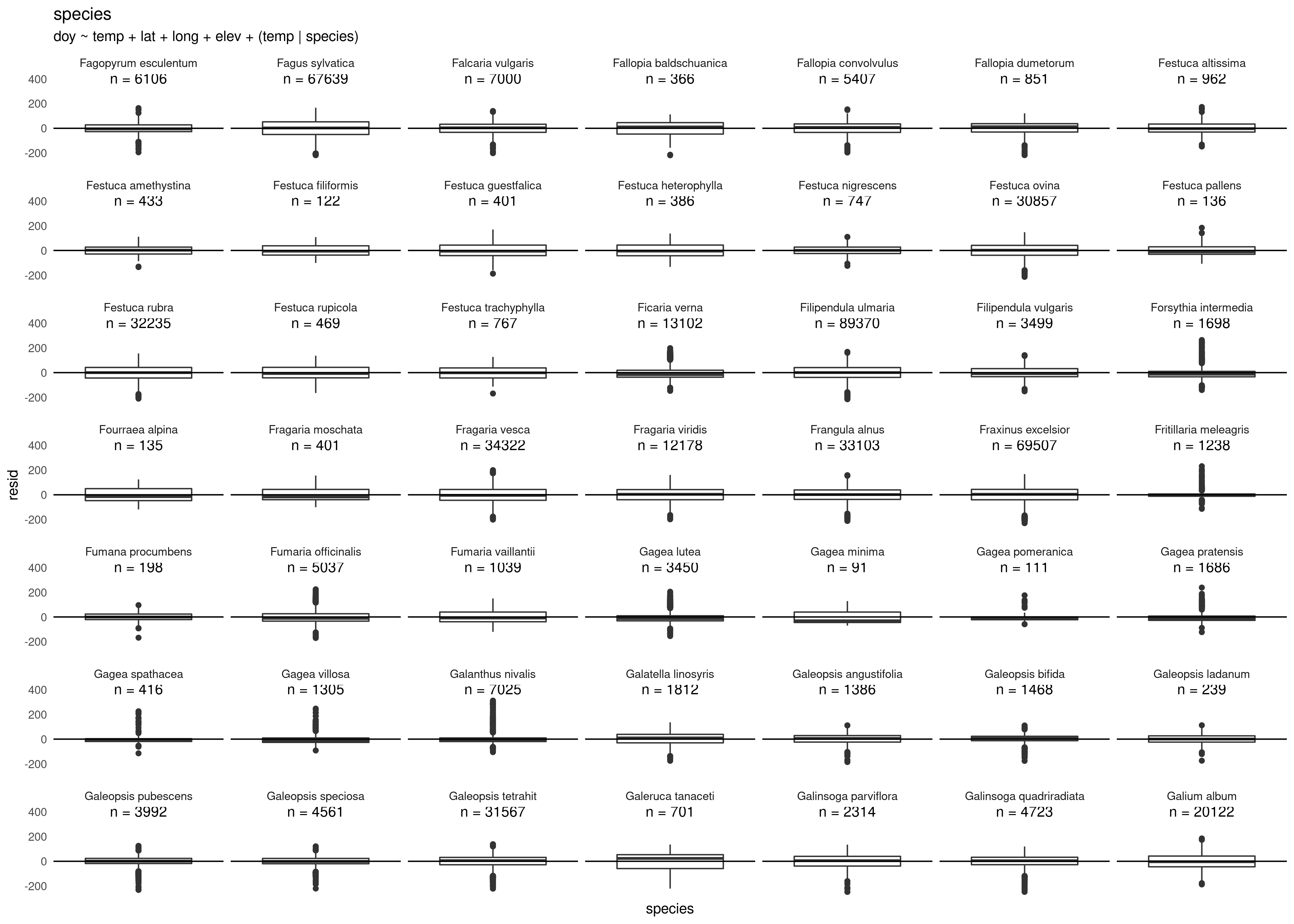

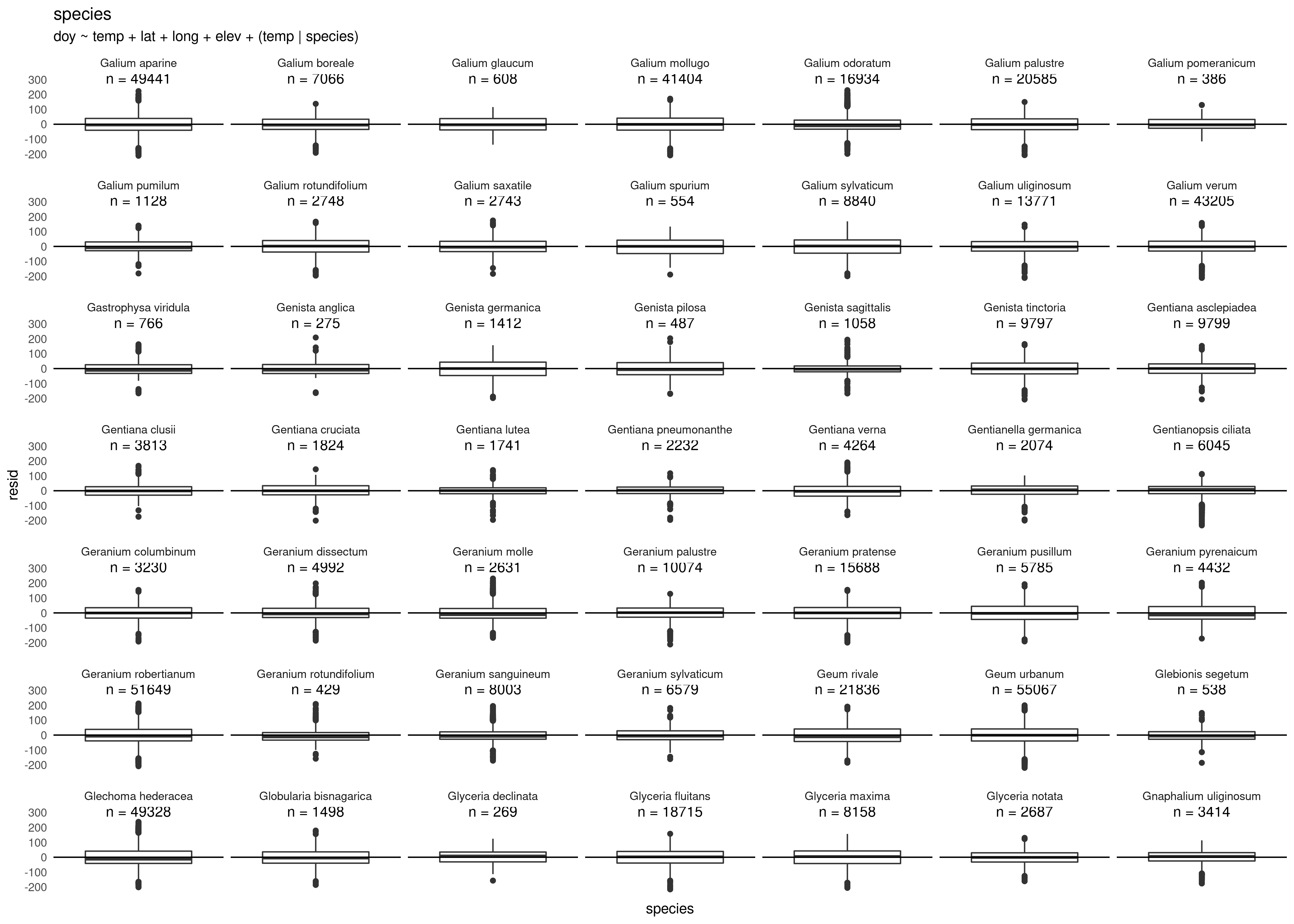

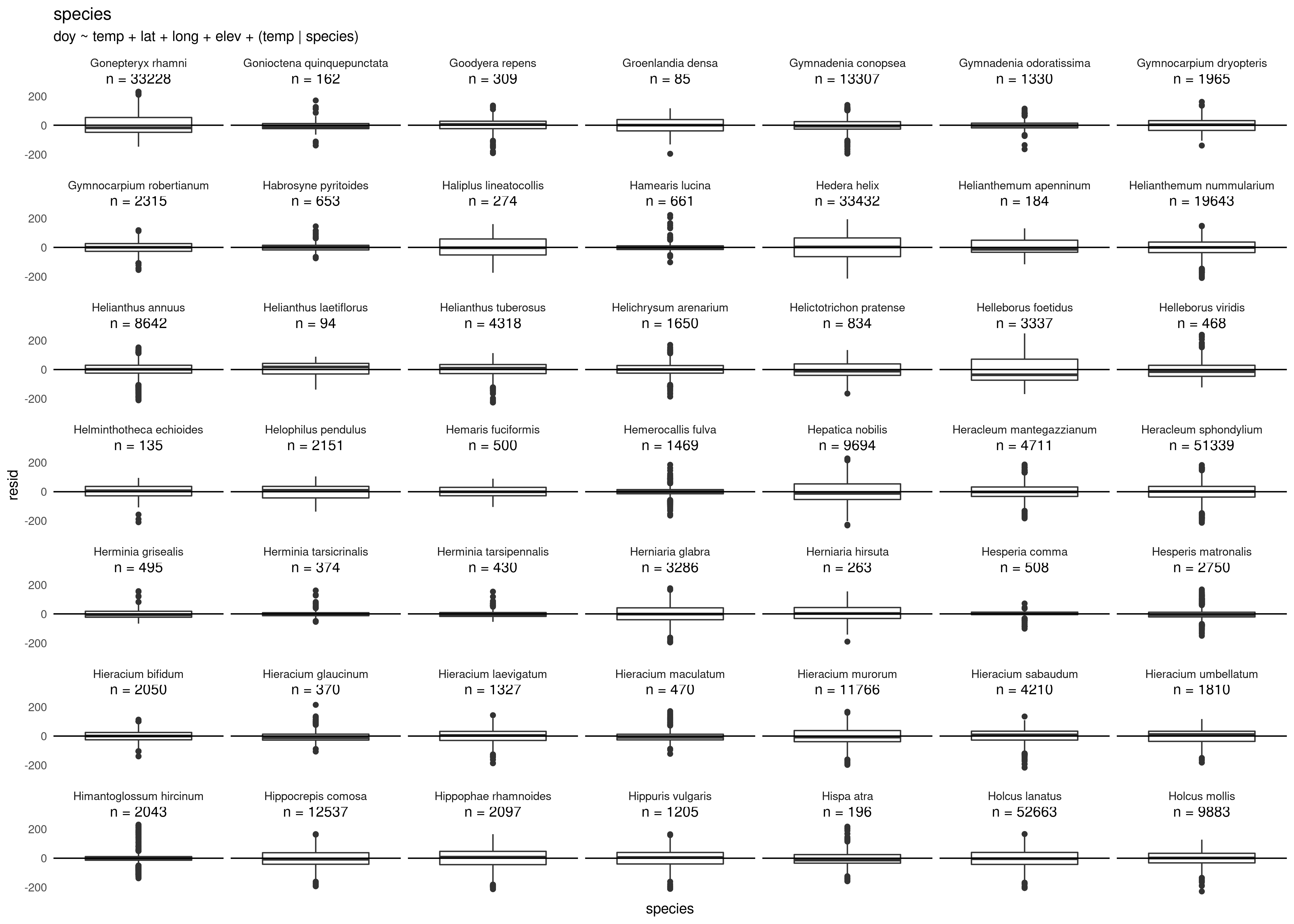

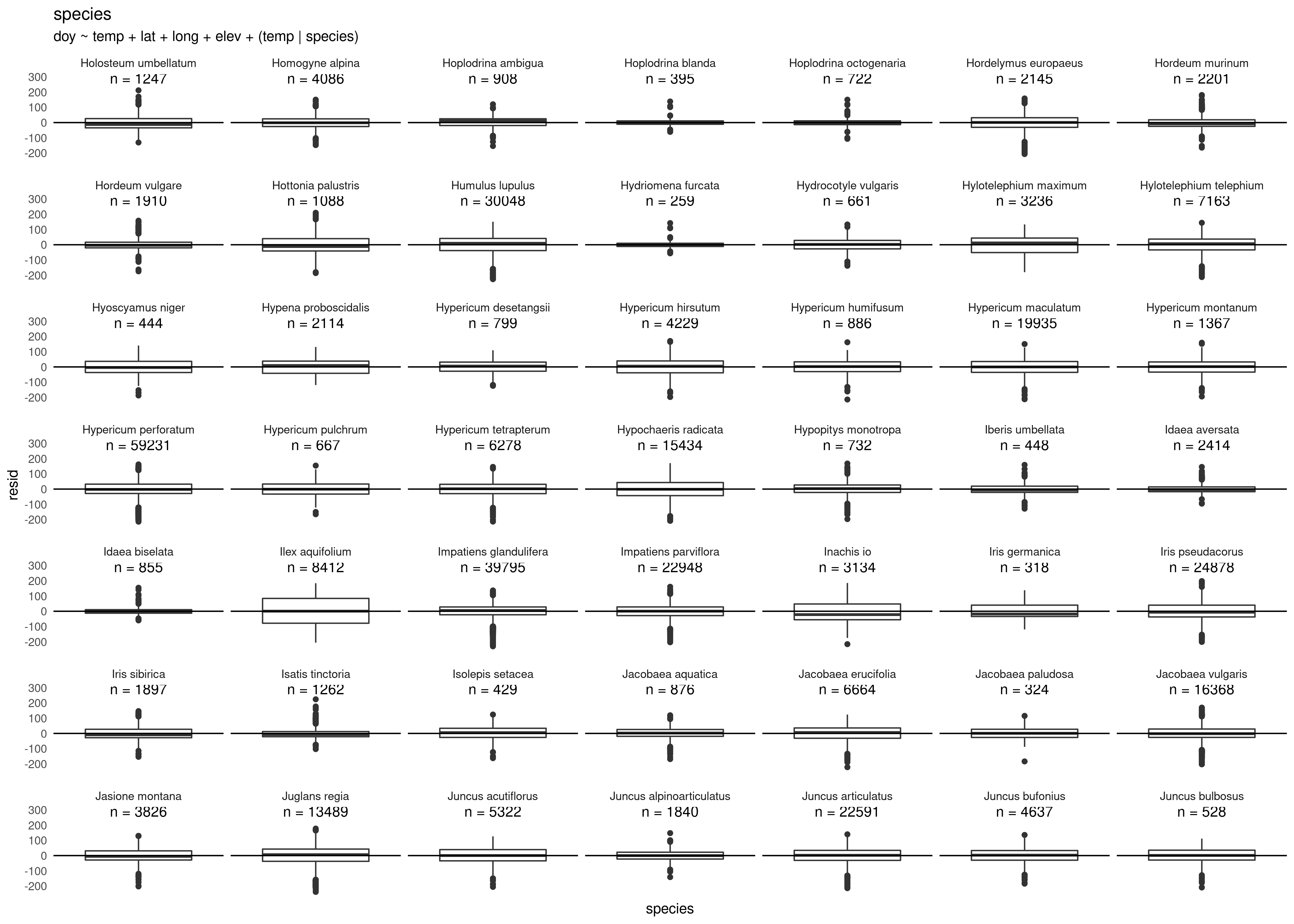

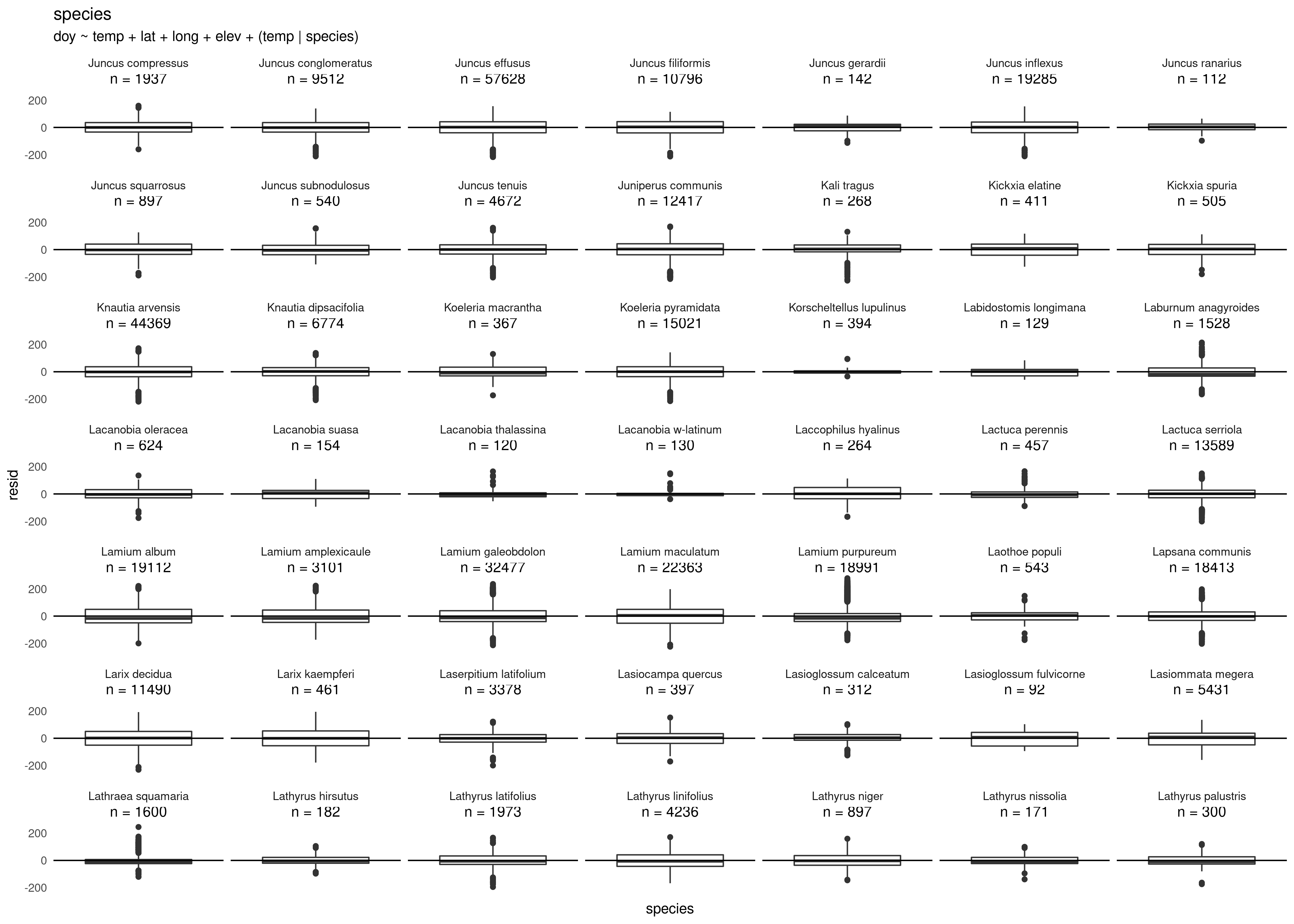

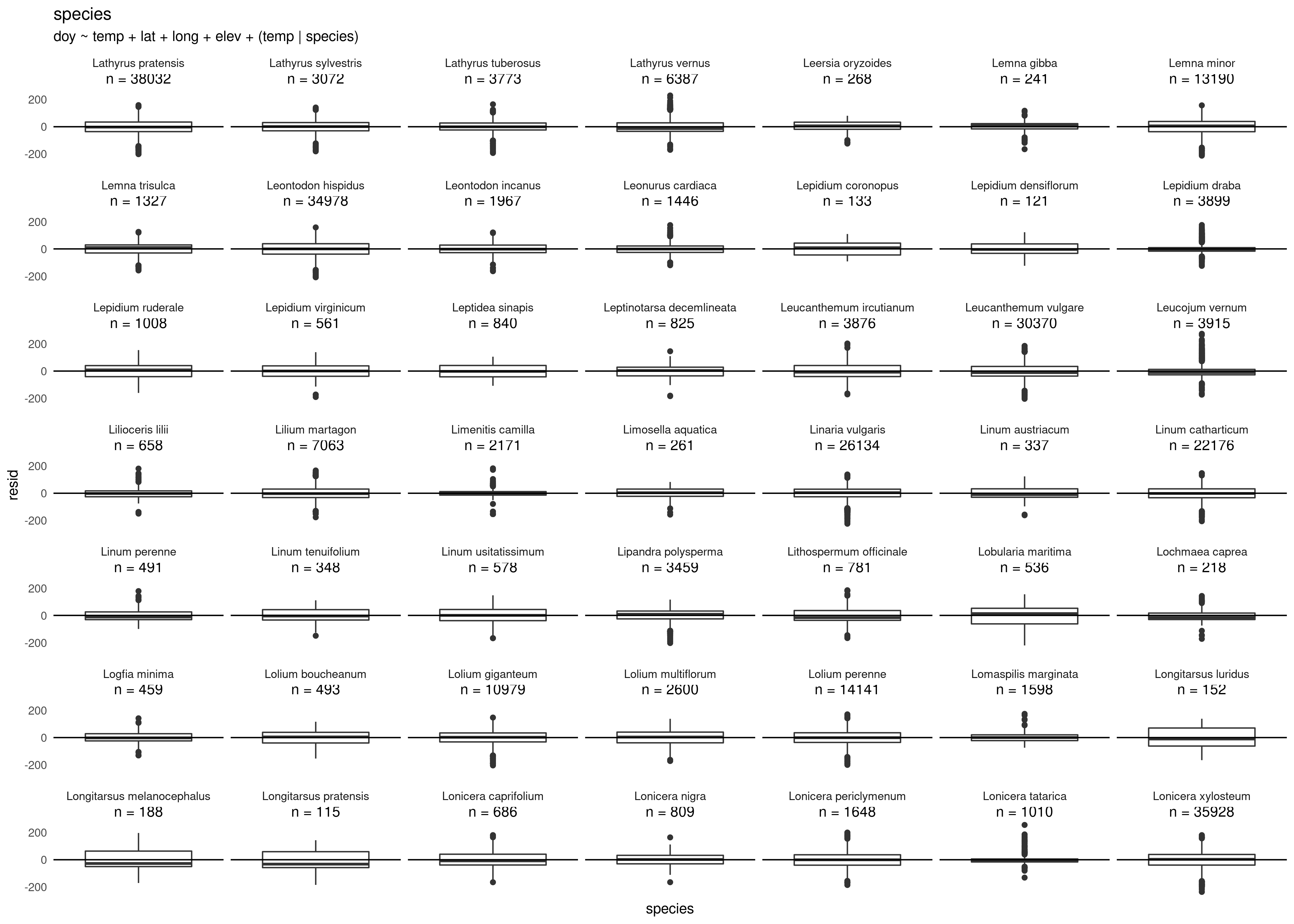

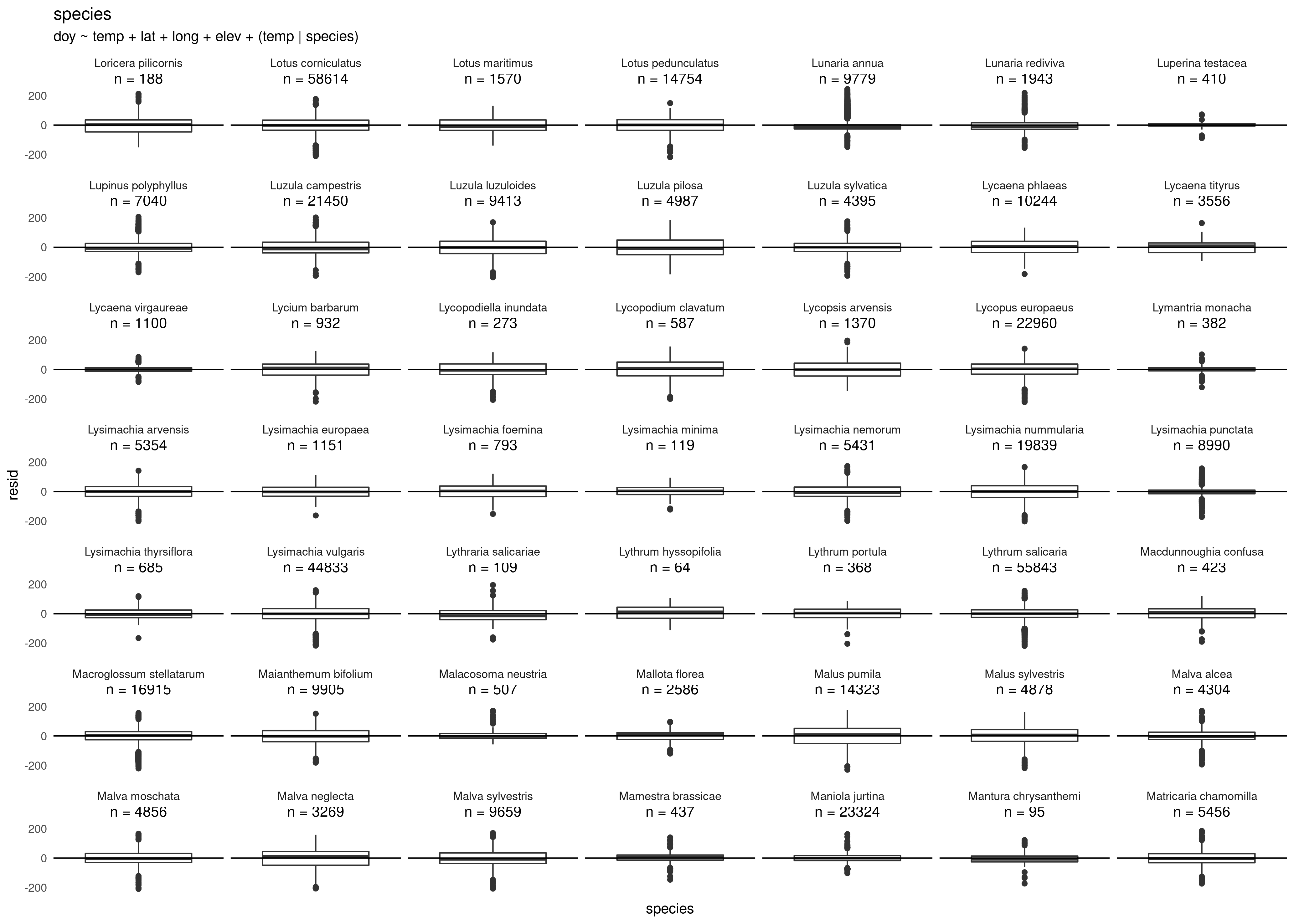

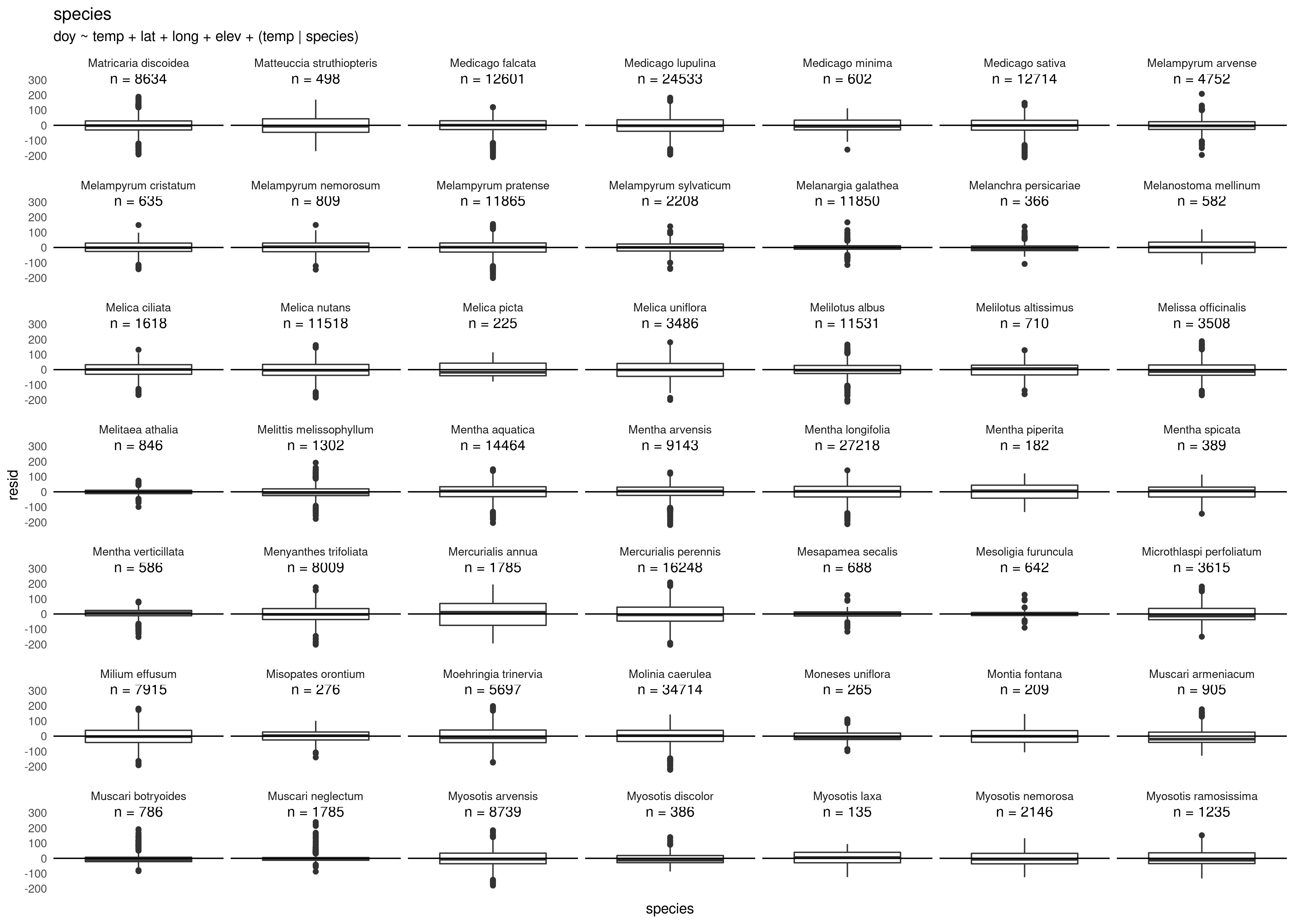

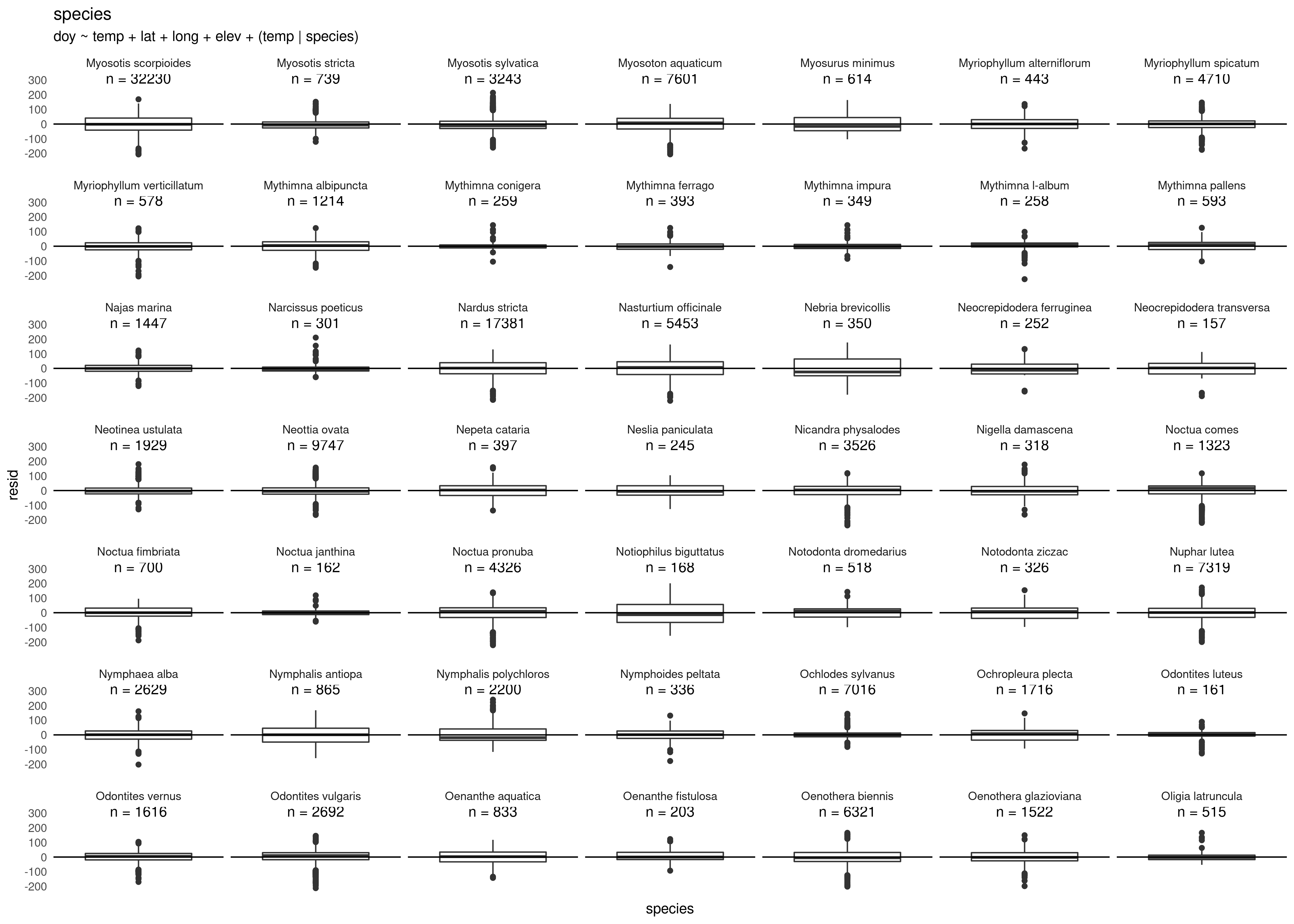

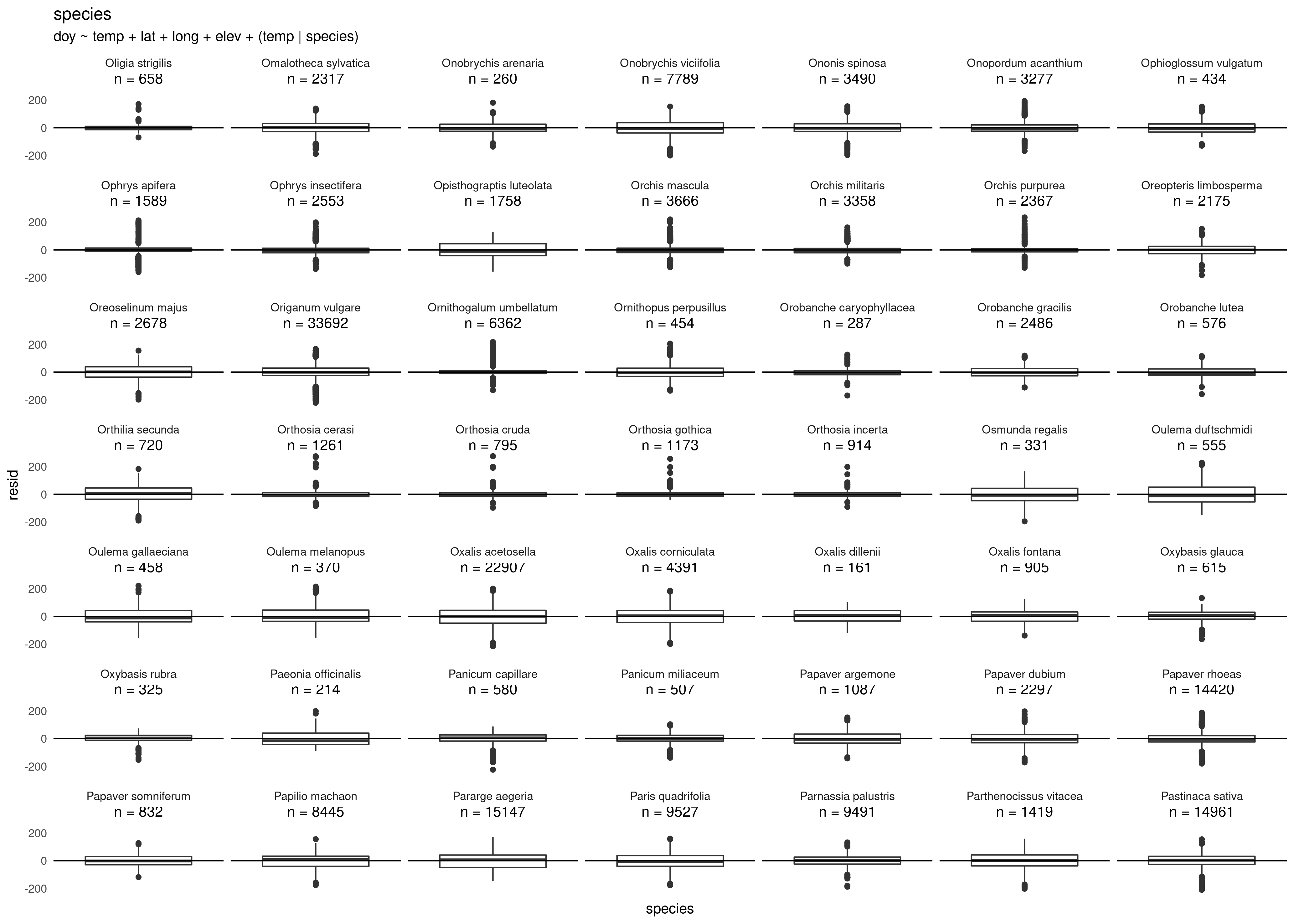

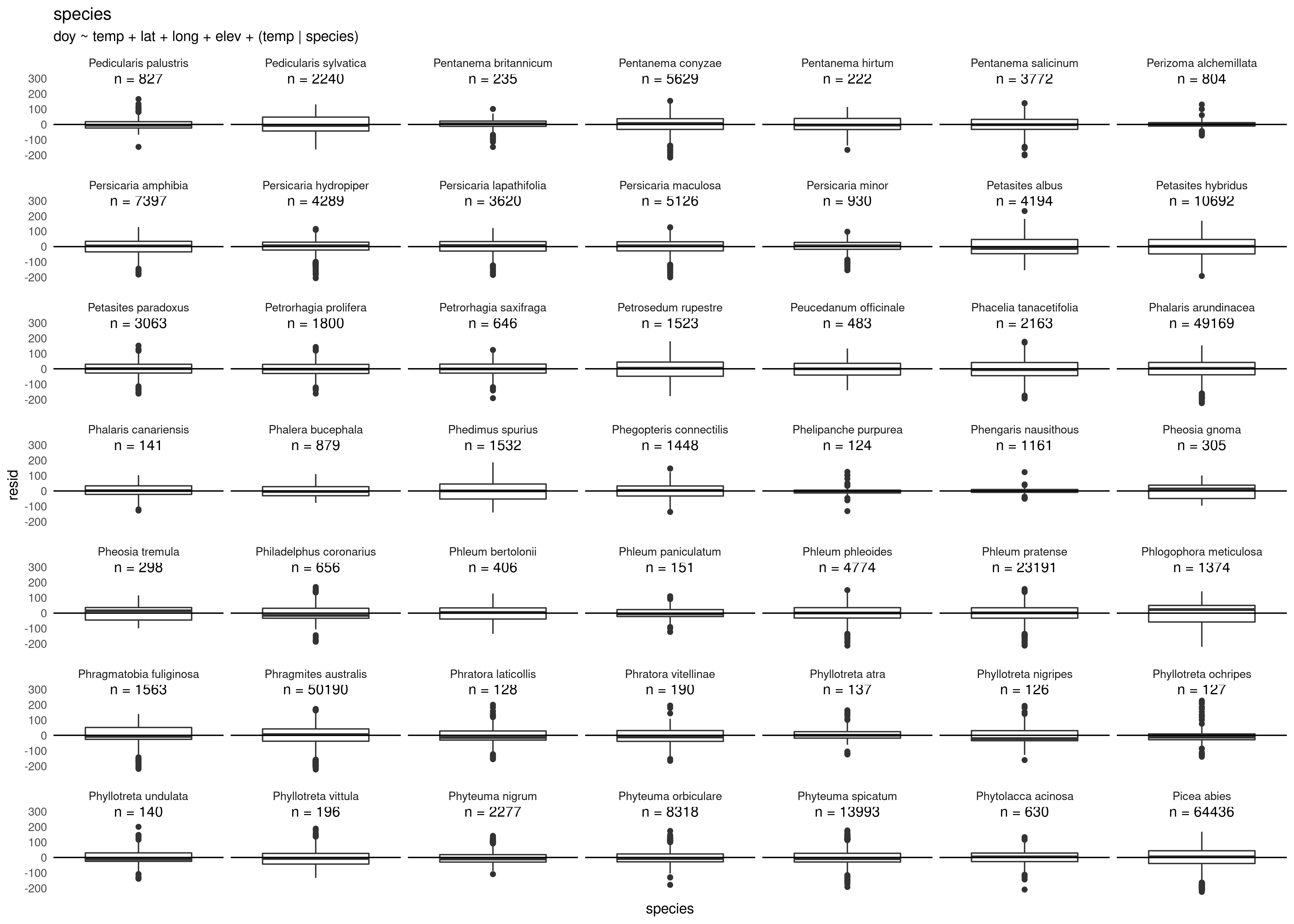

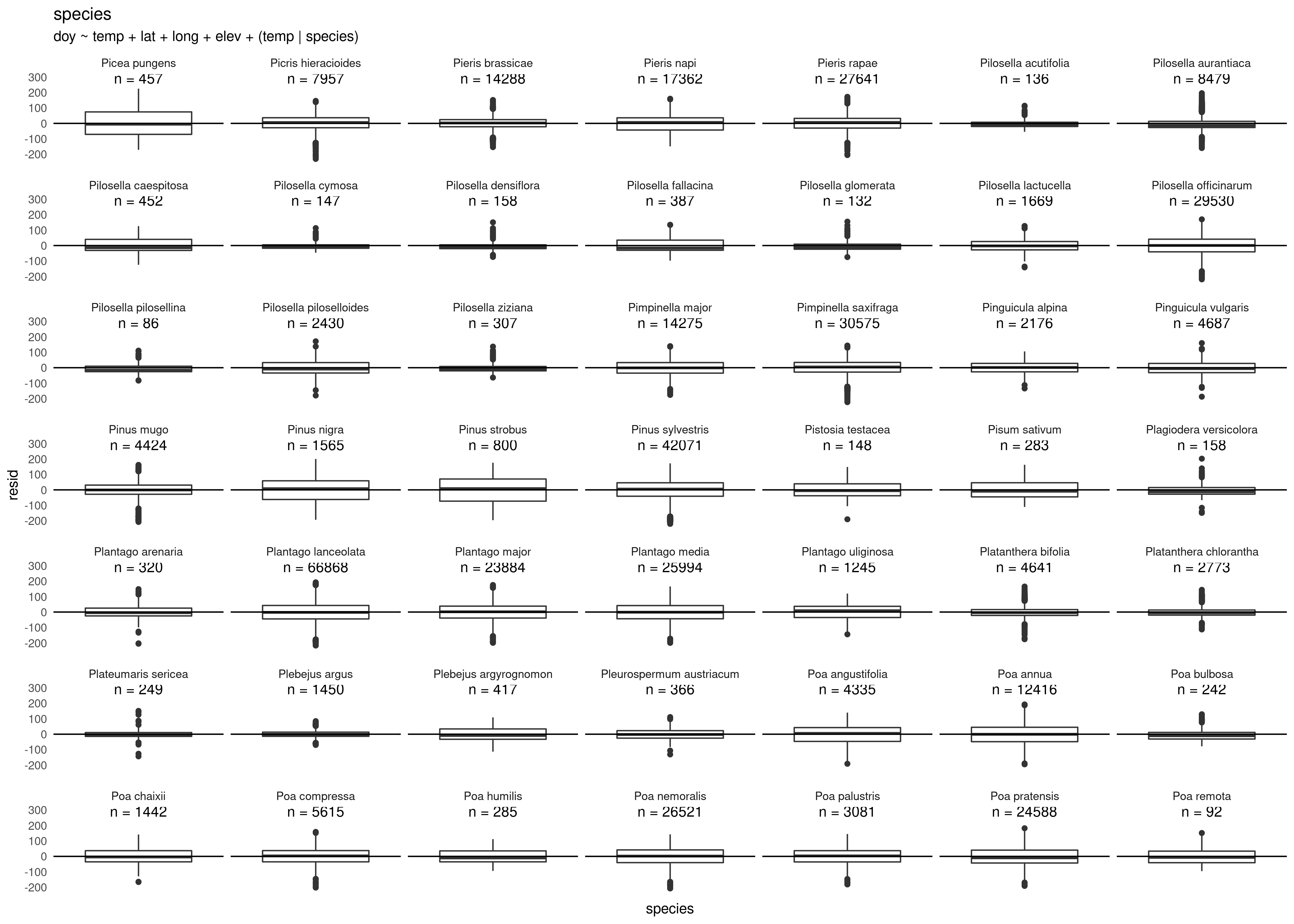

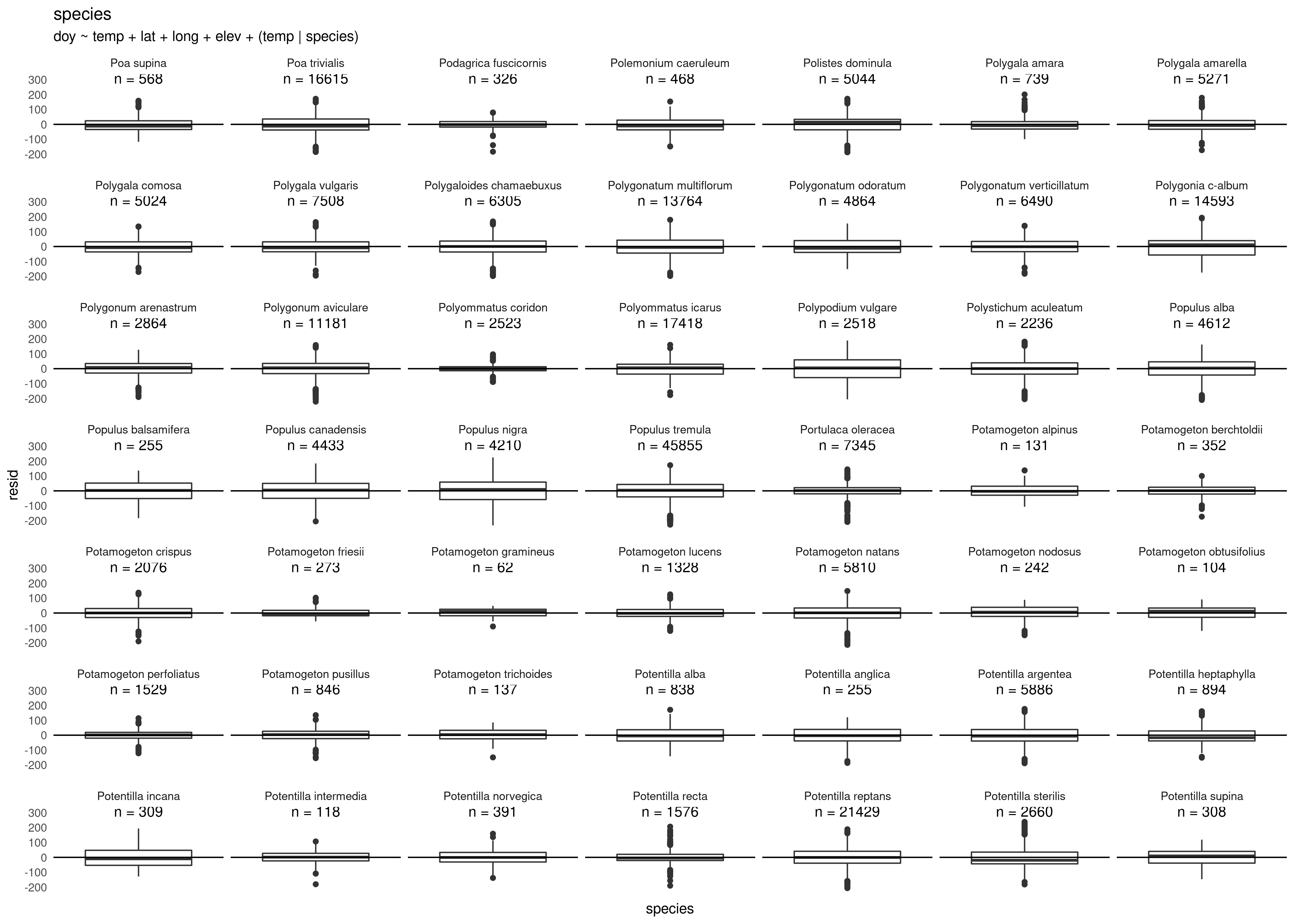

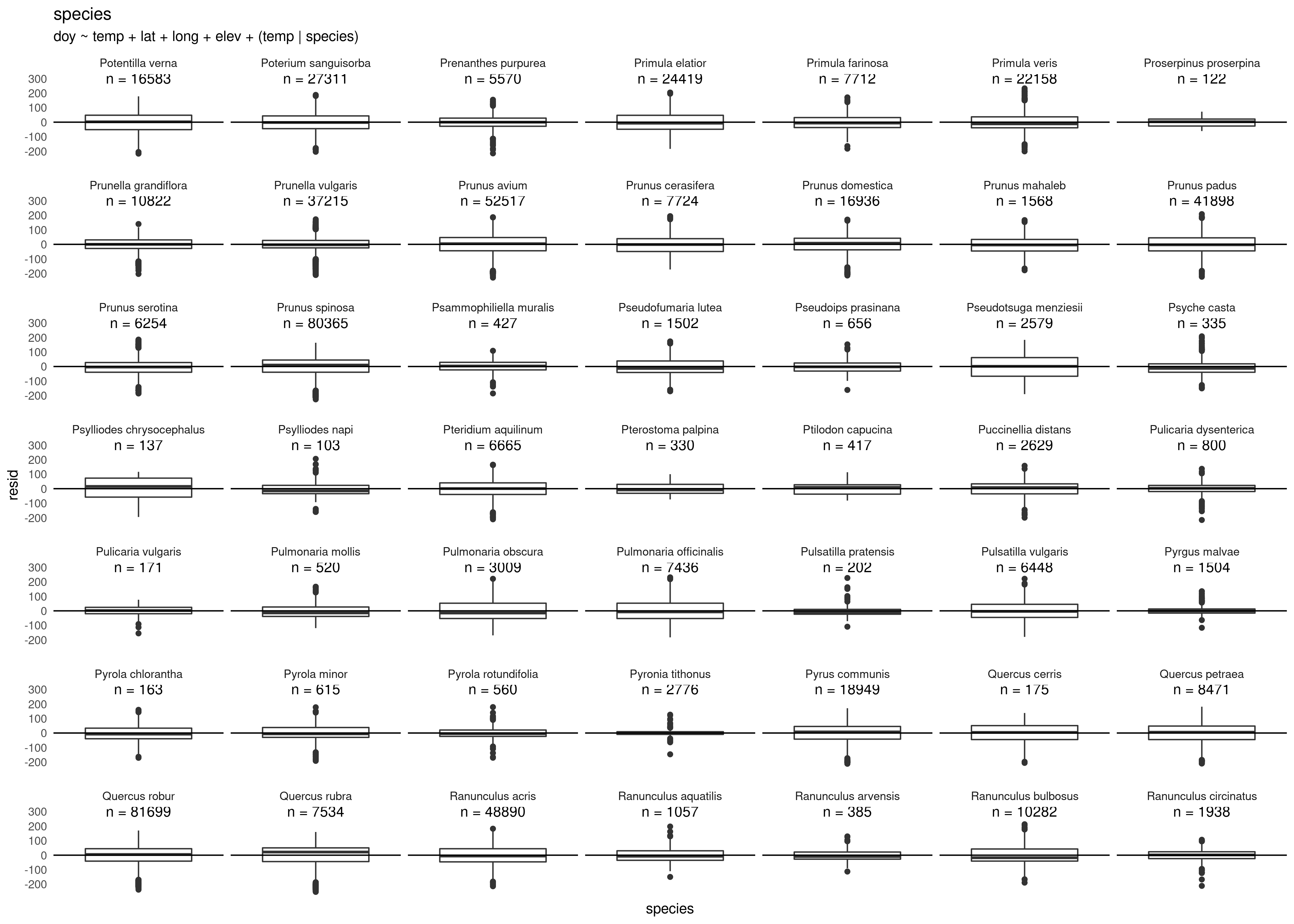

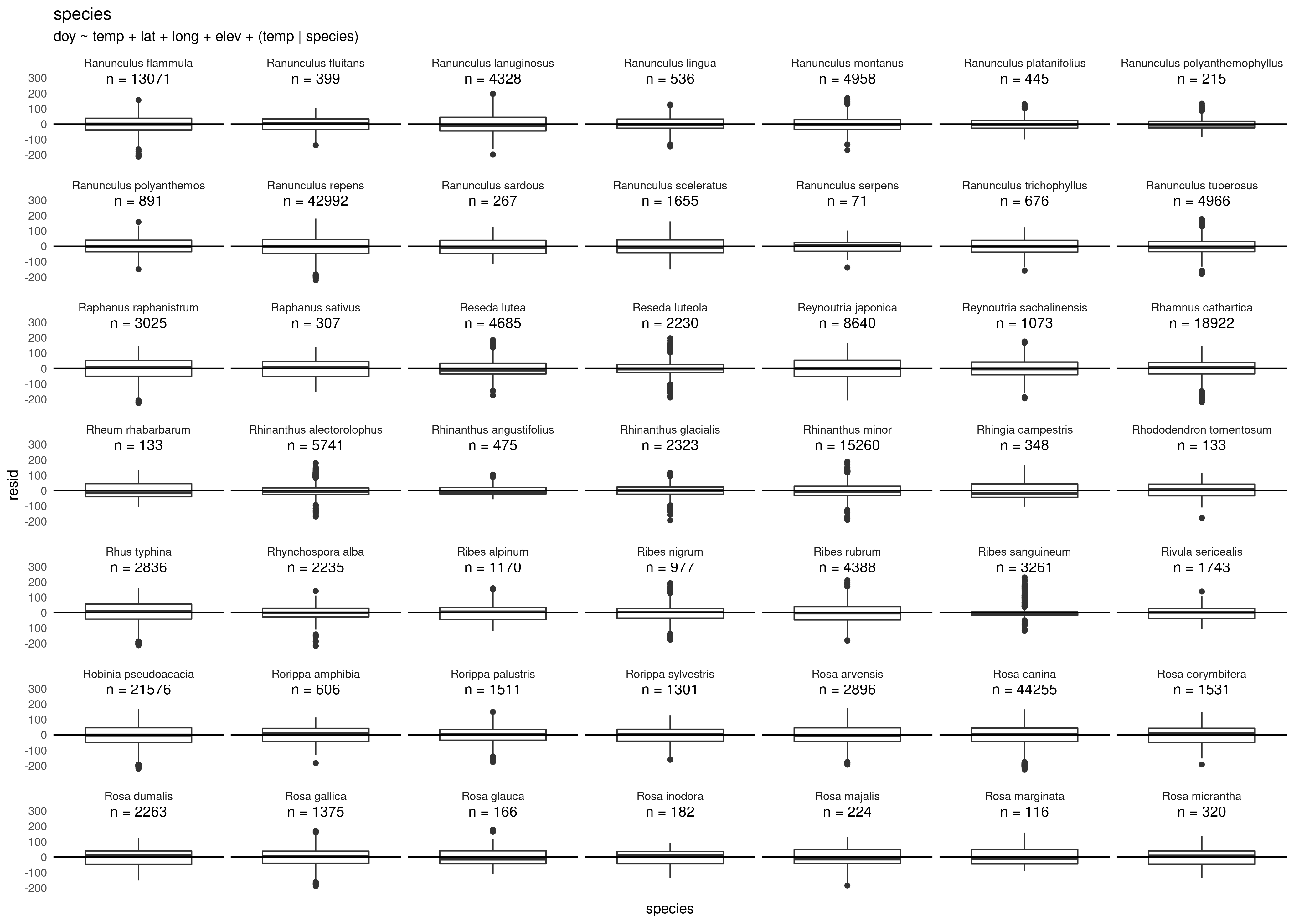

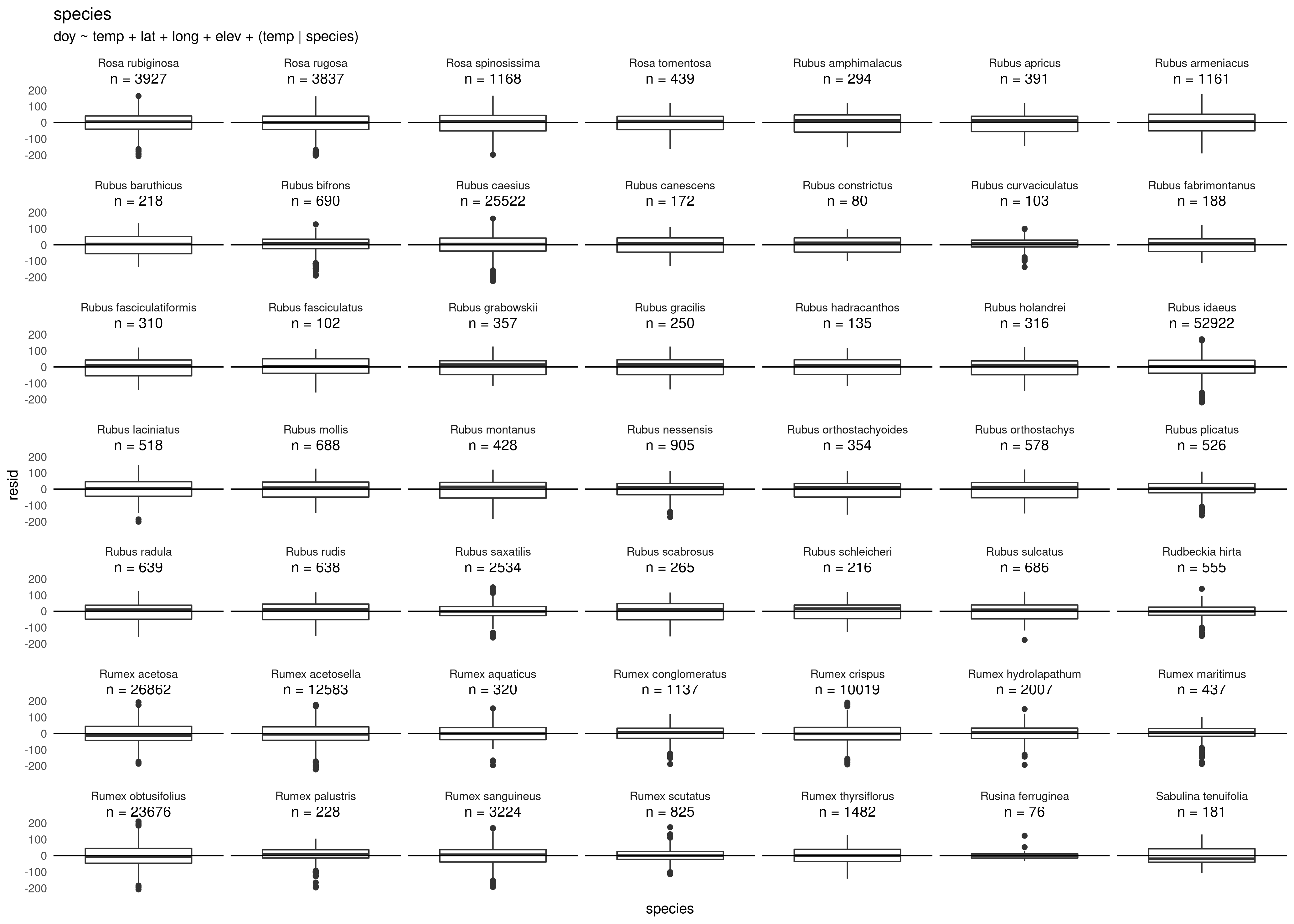

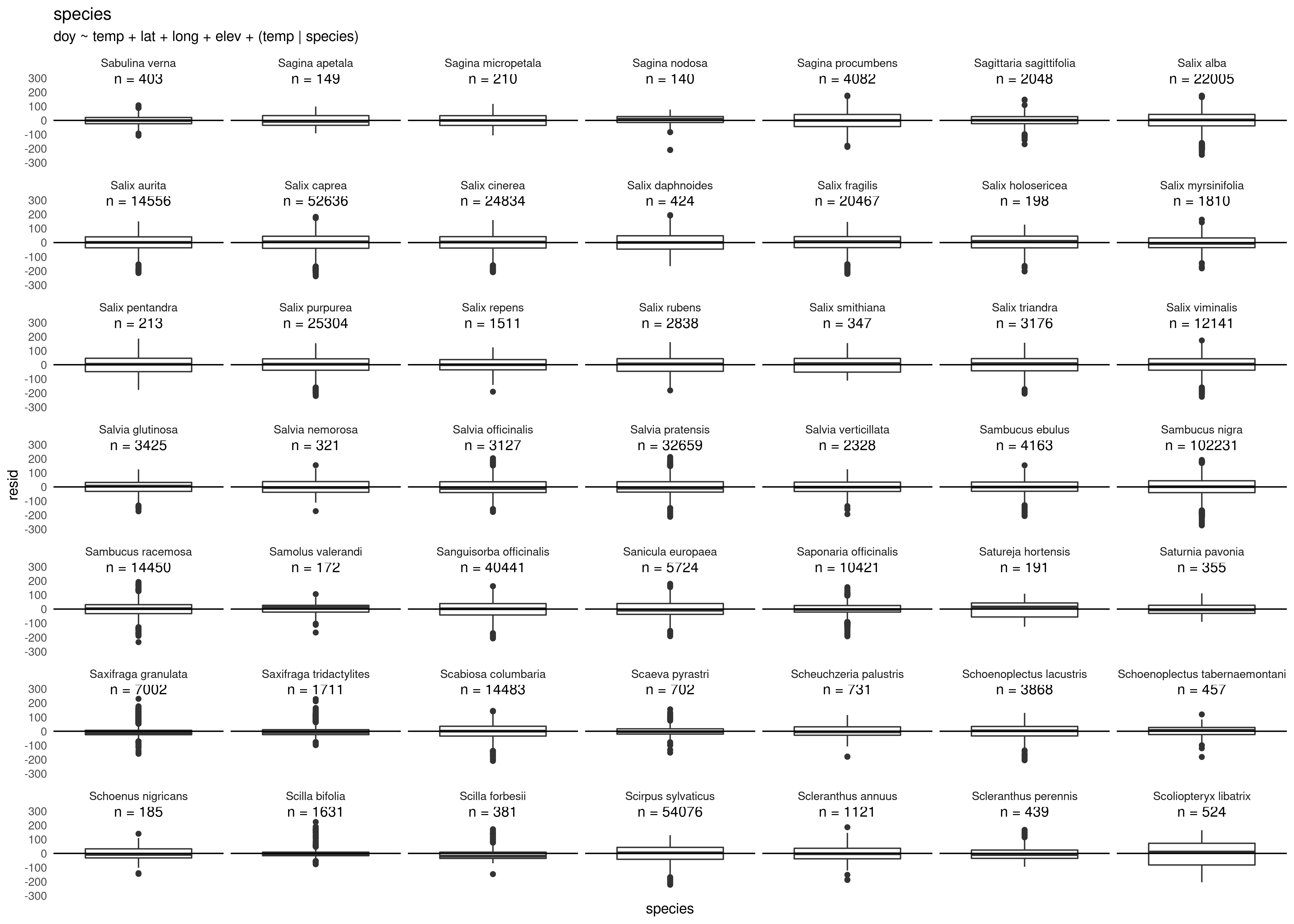

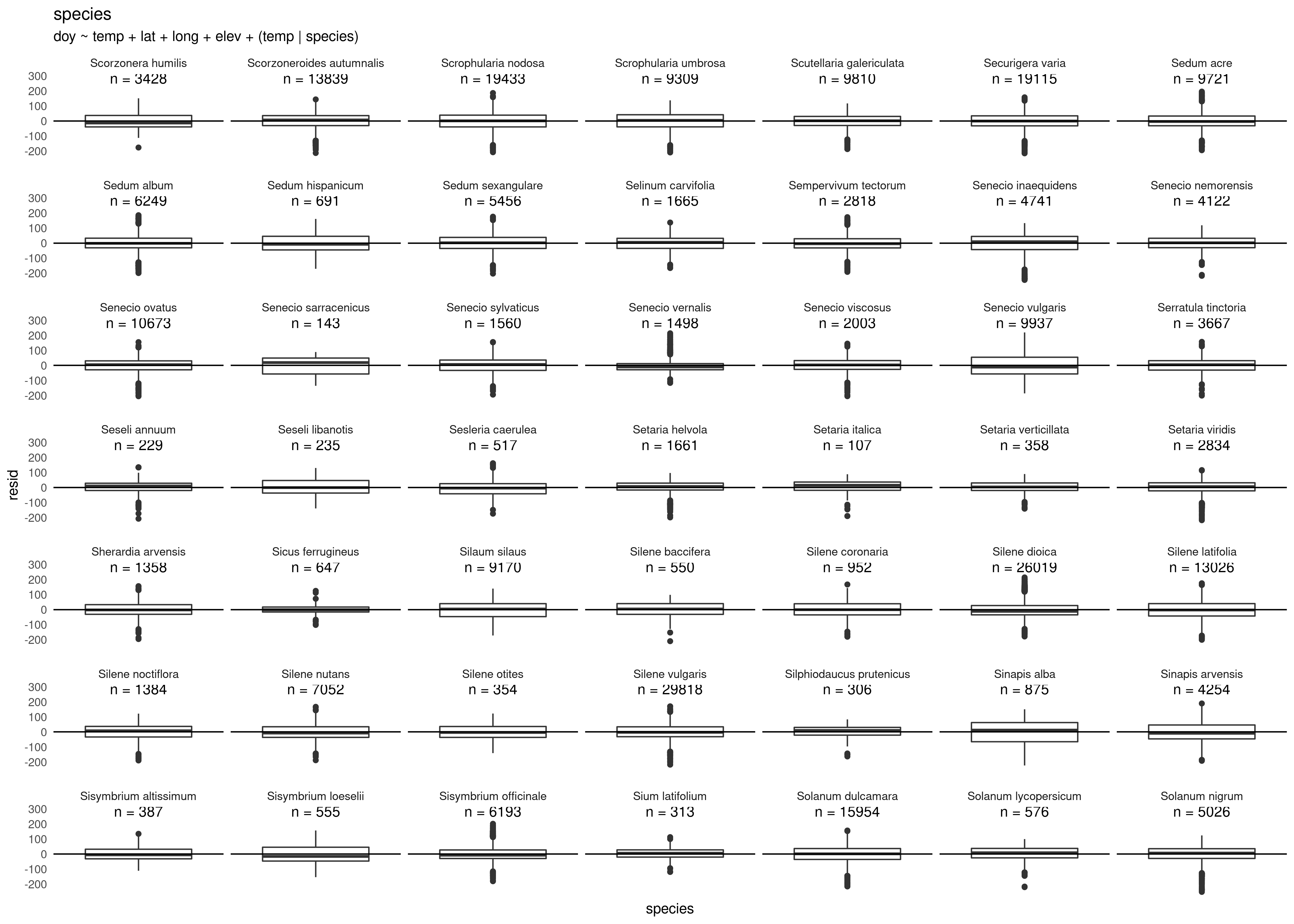

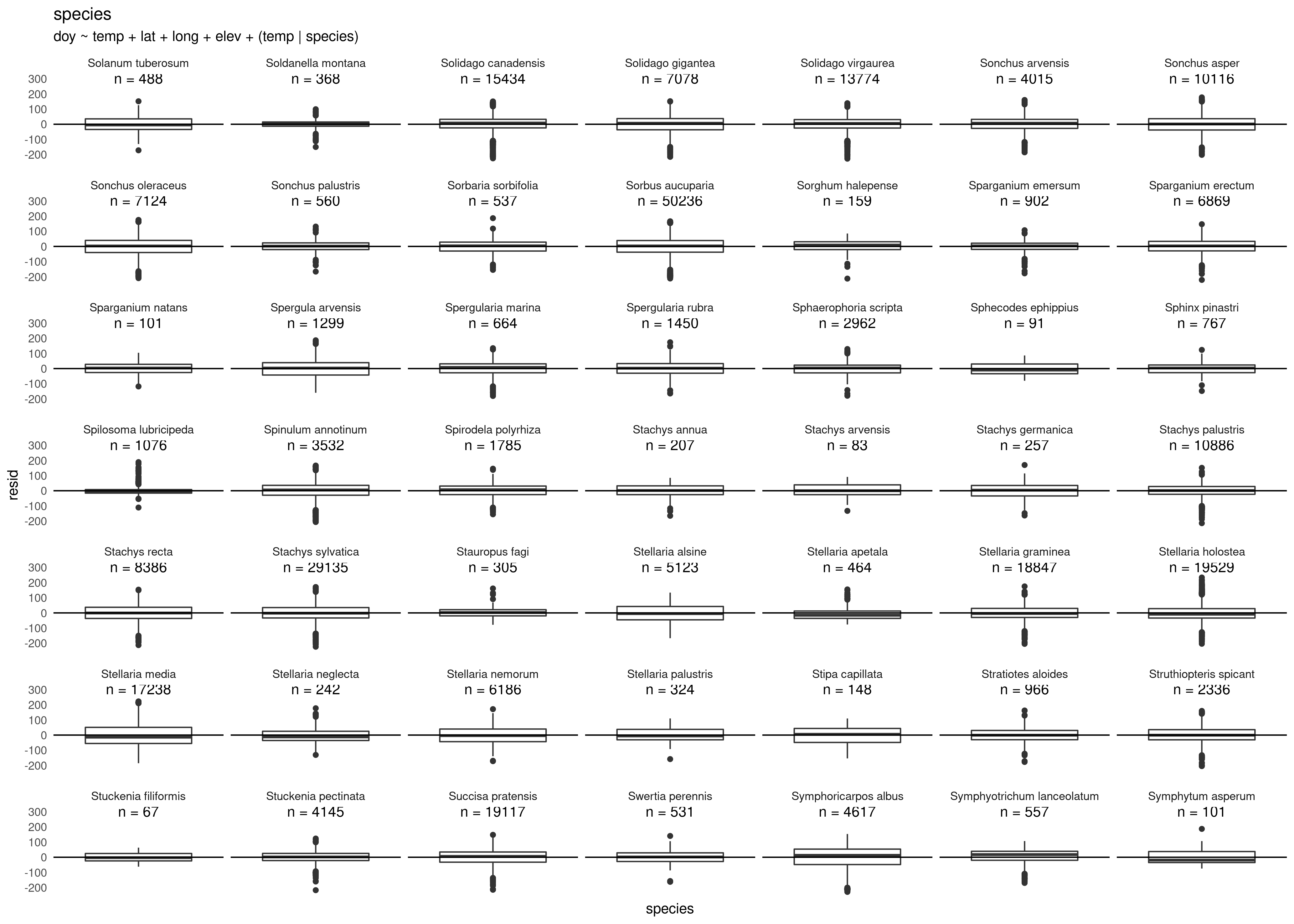

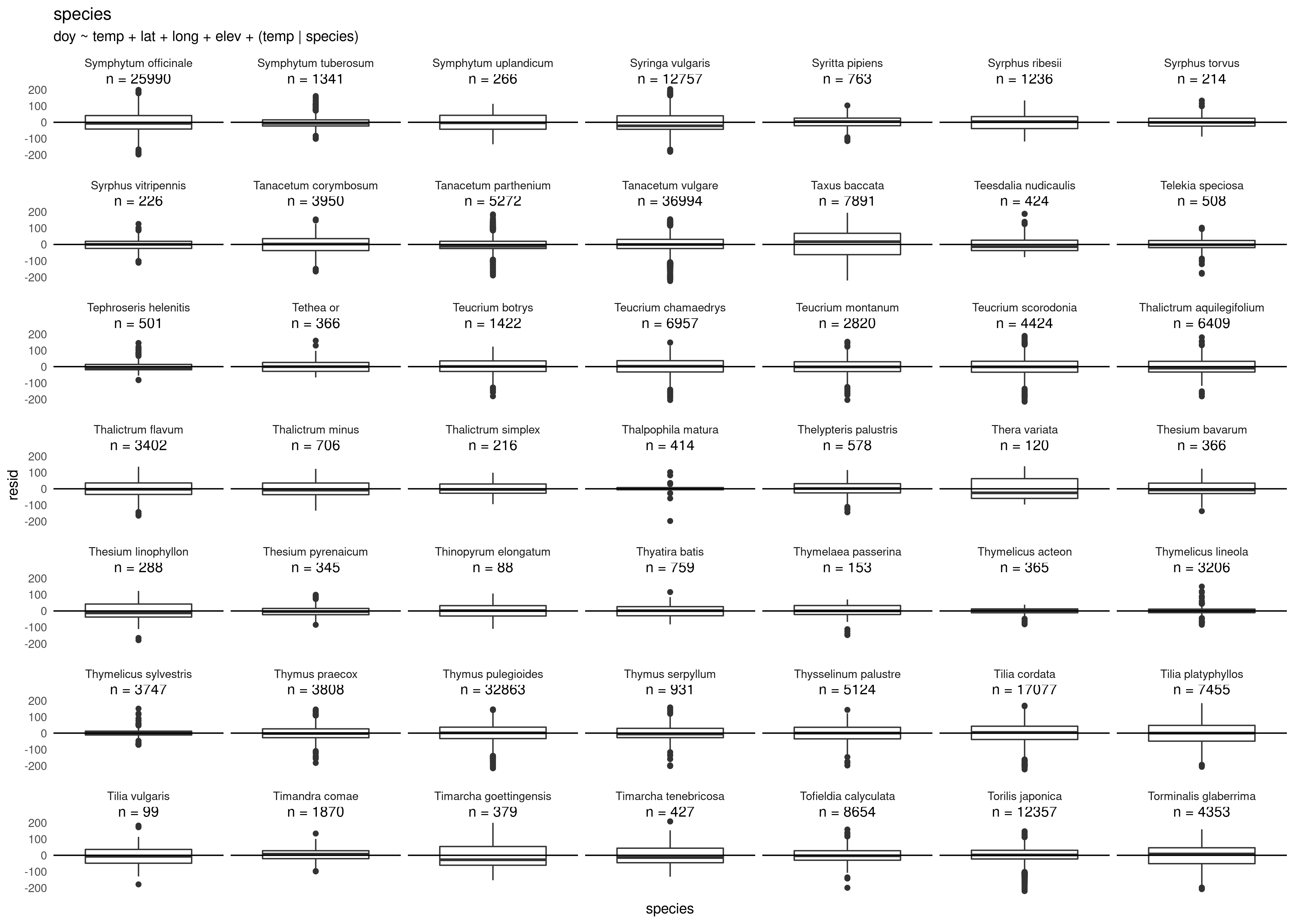

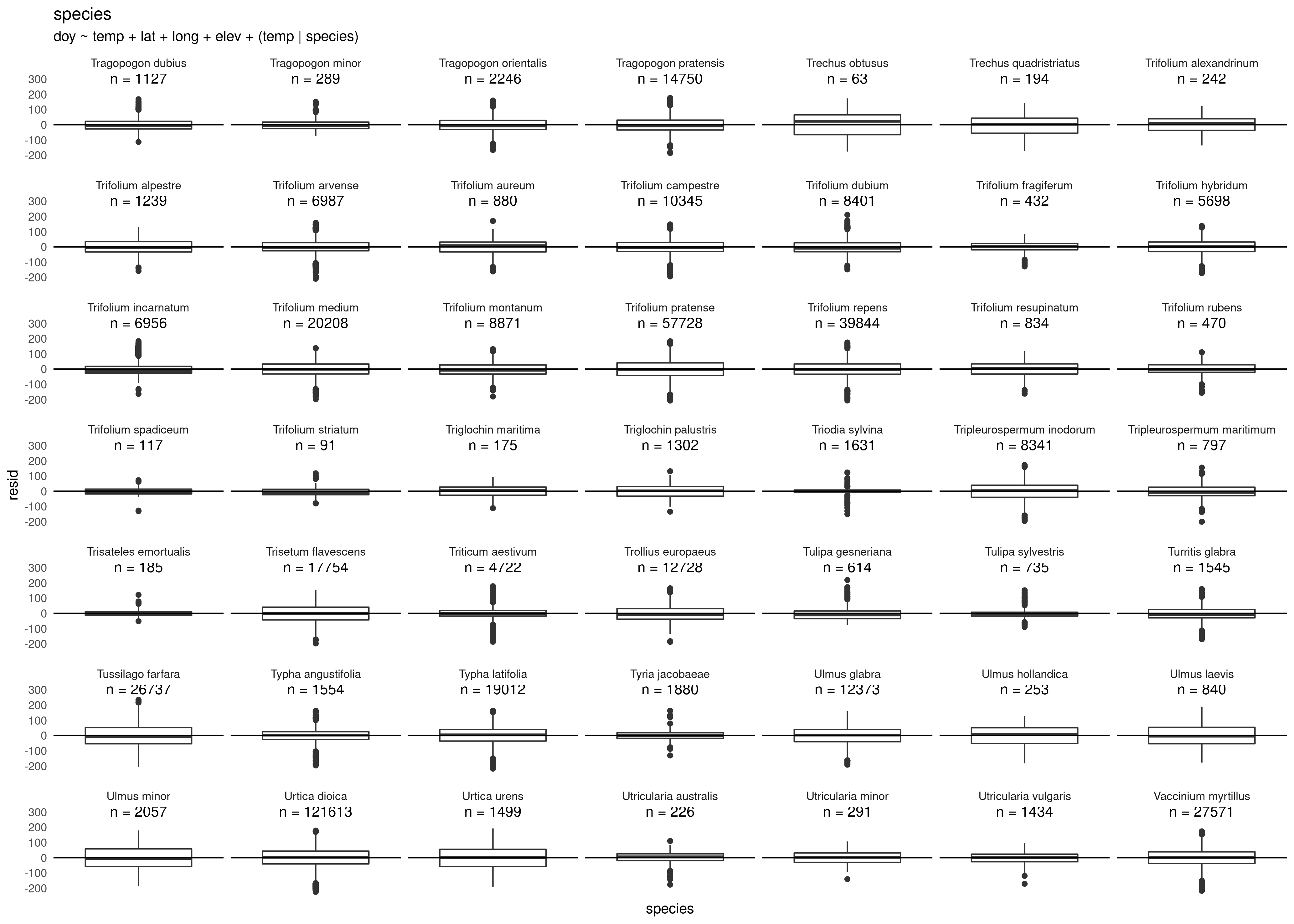

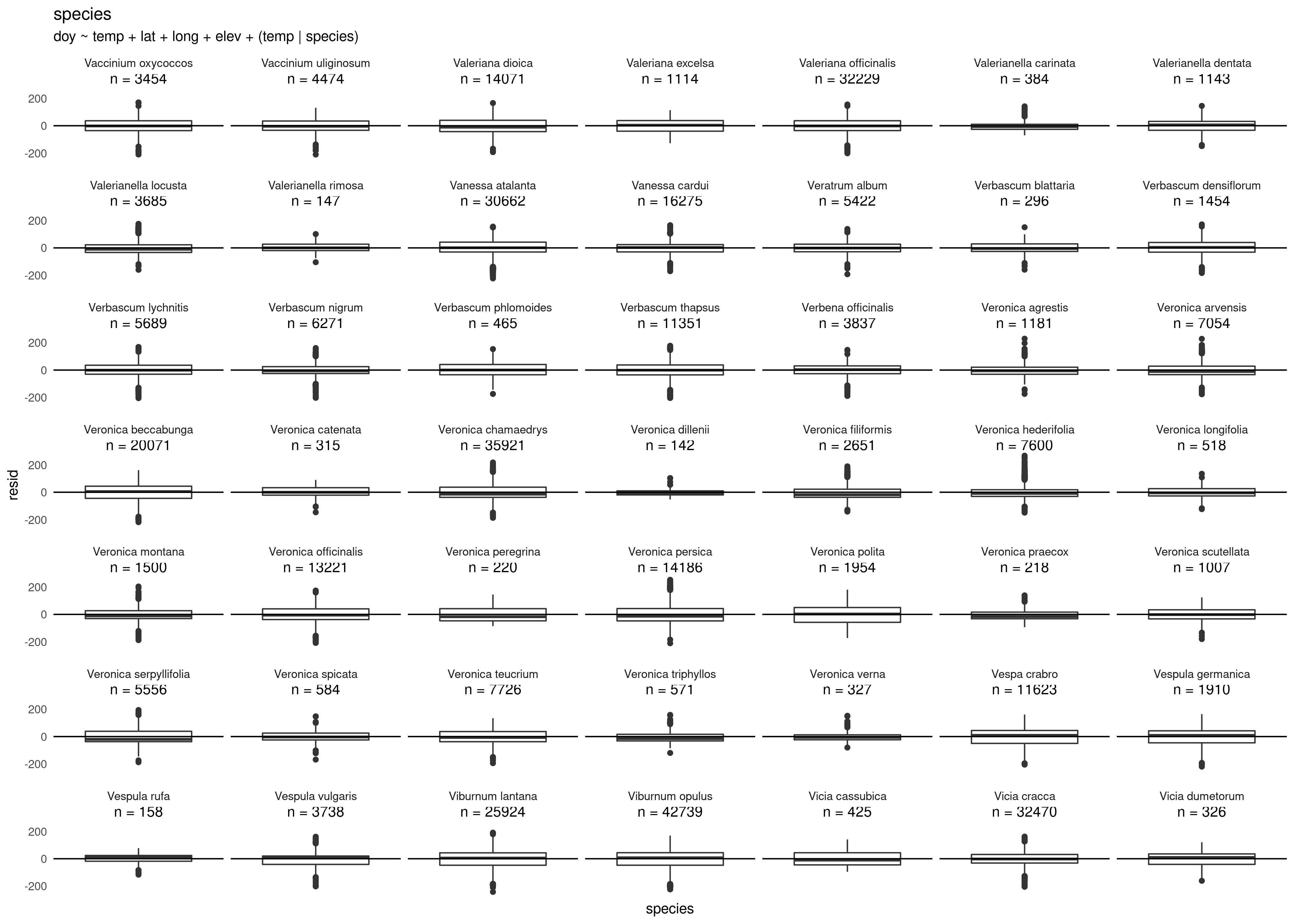

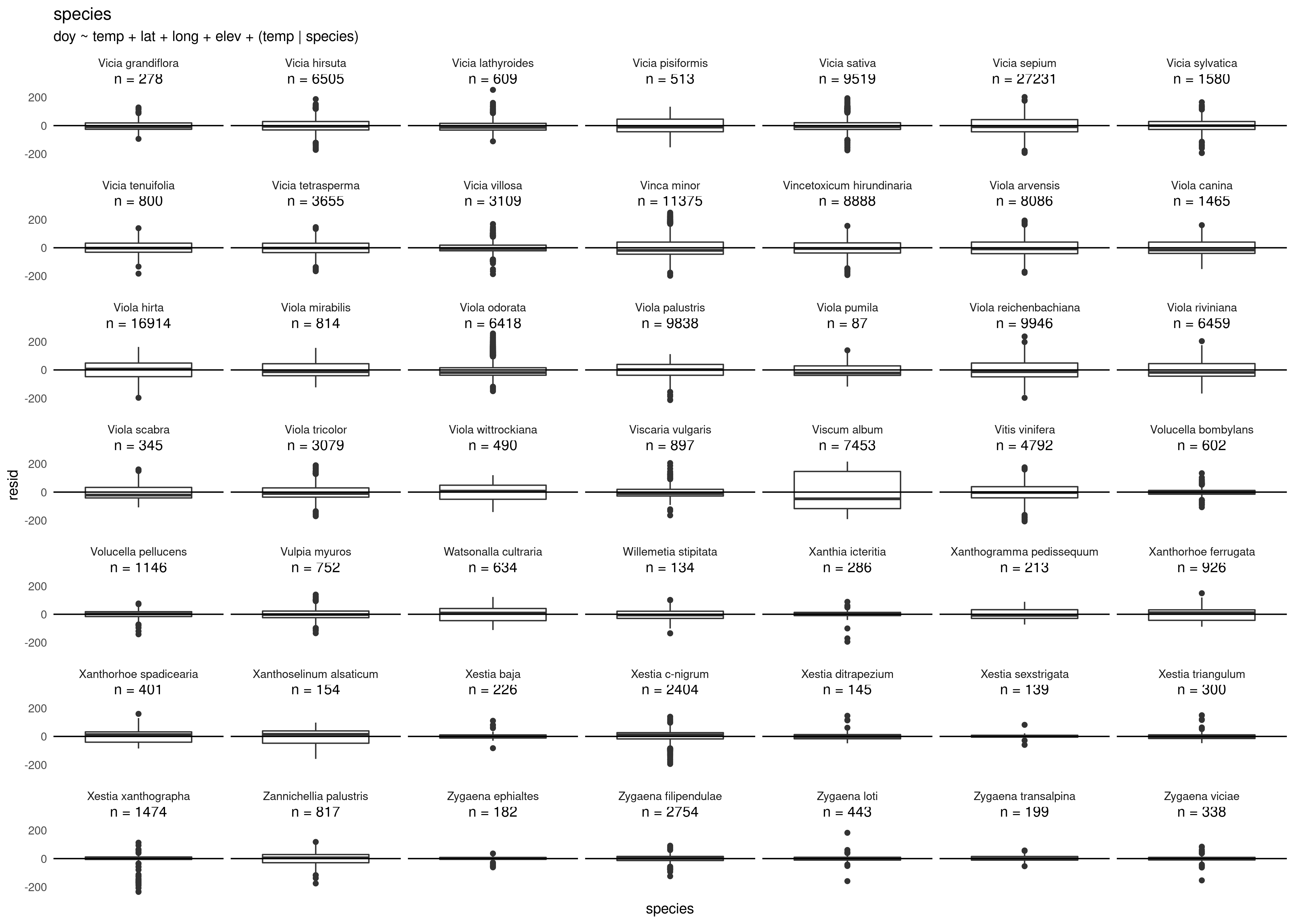

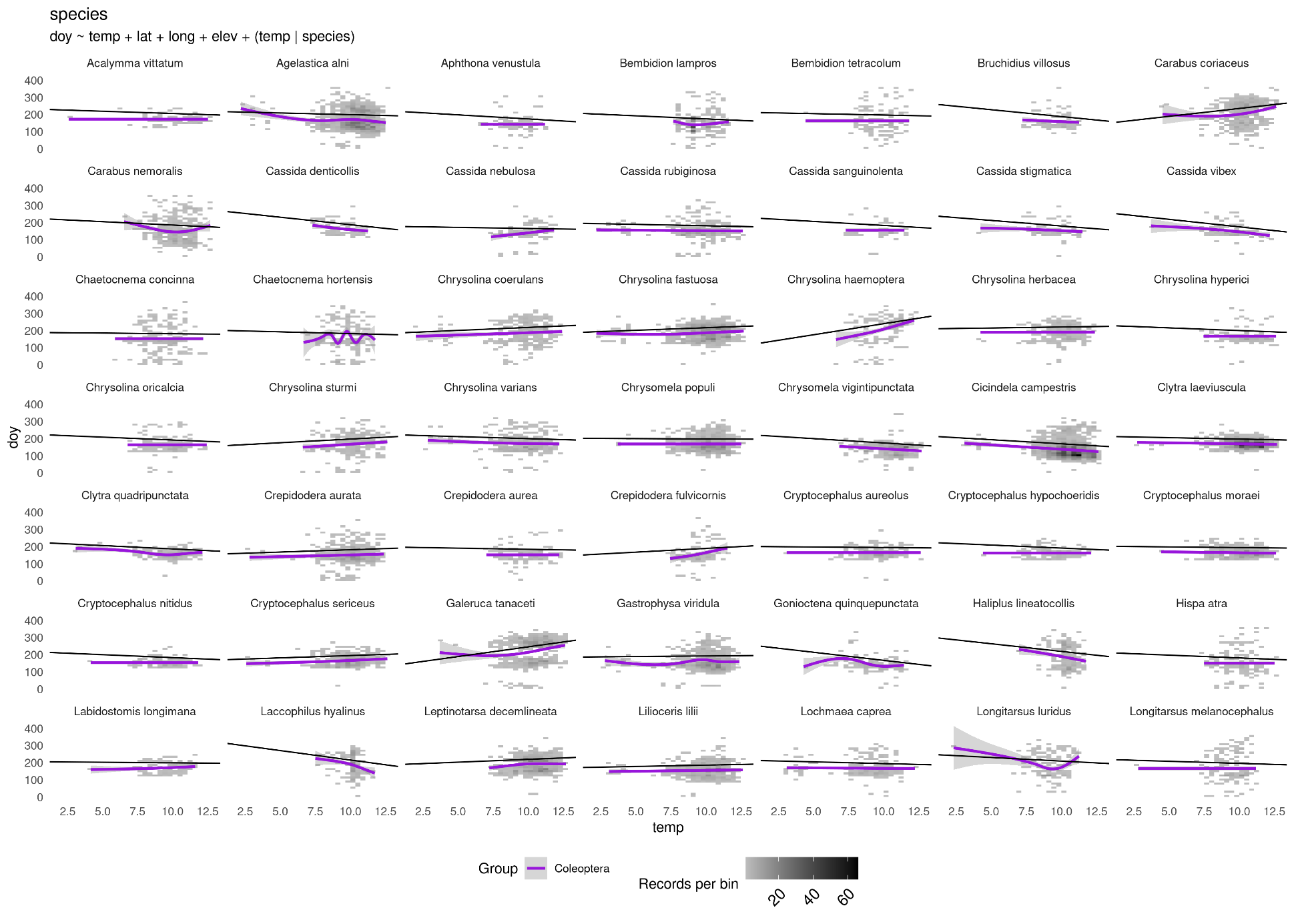

Diagnostic Plot 14 (and following) Species’ day of the year (DOY) distributions across record yearly mean temperatures in degrees Celsius. Coloured lines indicate generalized additive model curves fitted to the data; black lines indicate linear mixed effects temperature model slopes that were used in the analysis. Cell shading indicates point density.

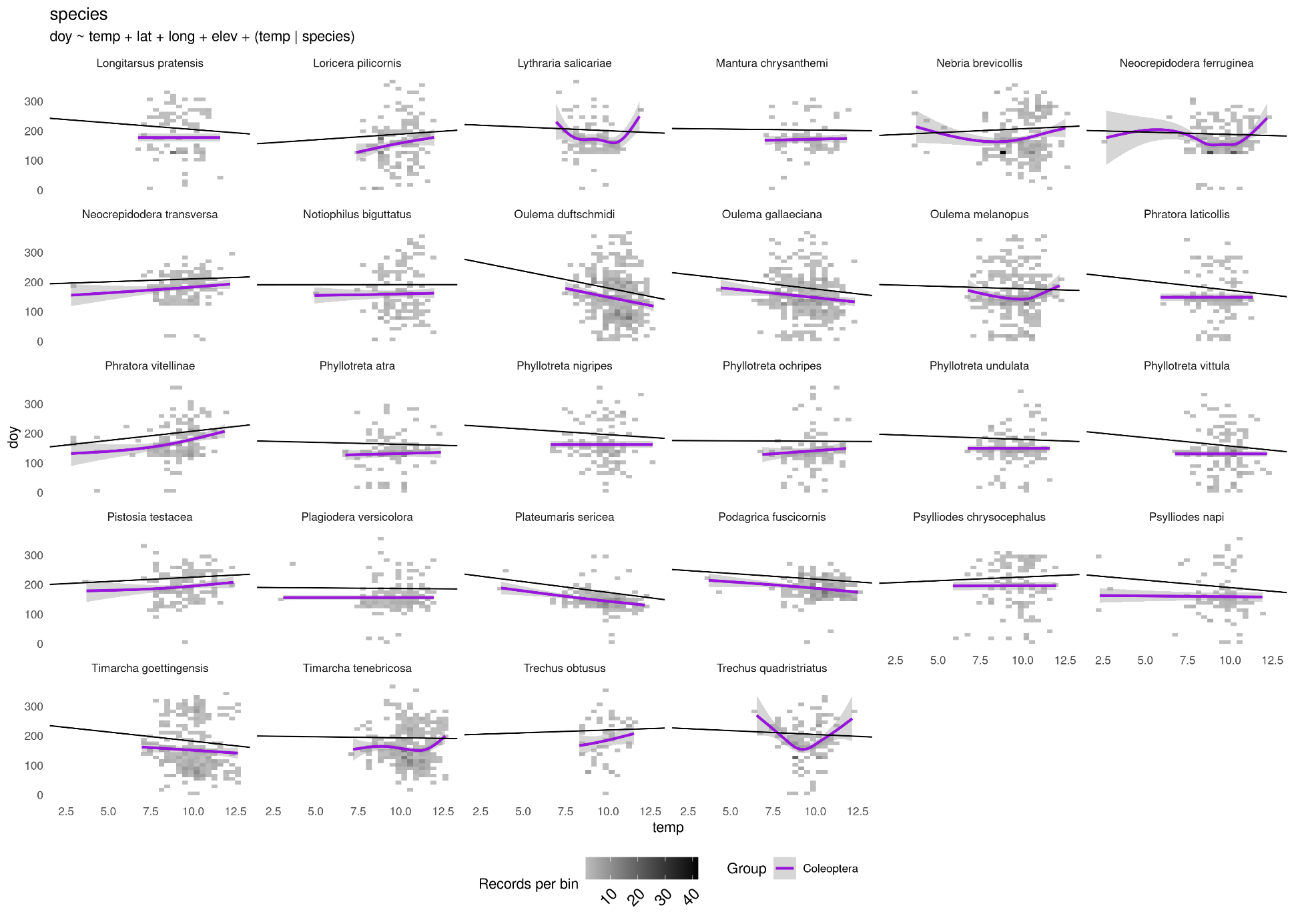

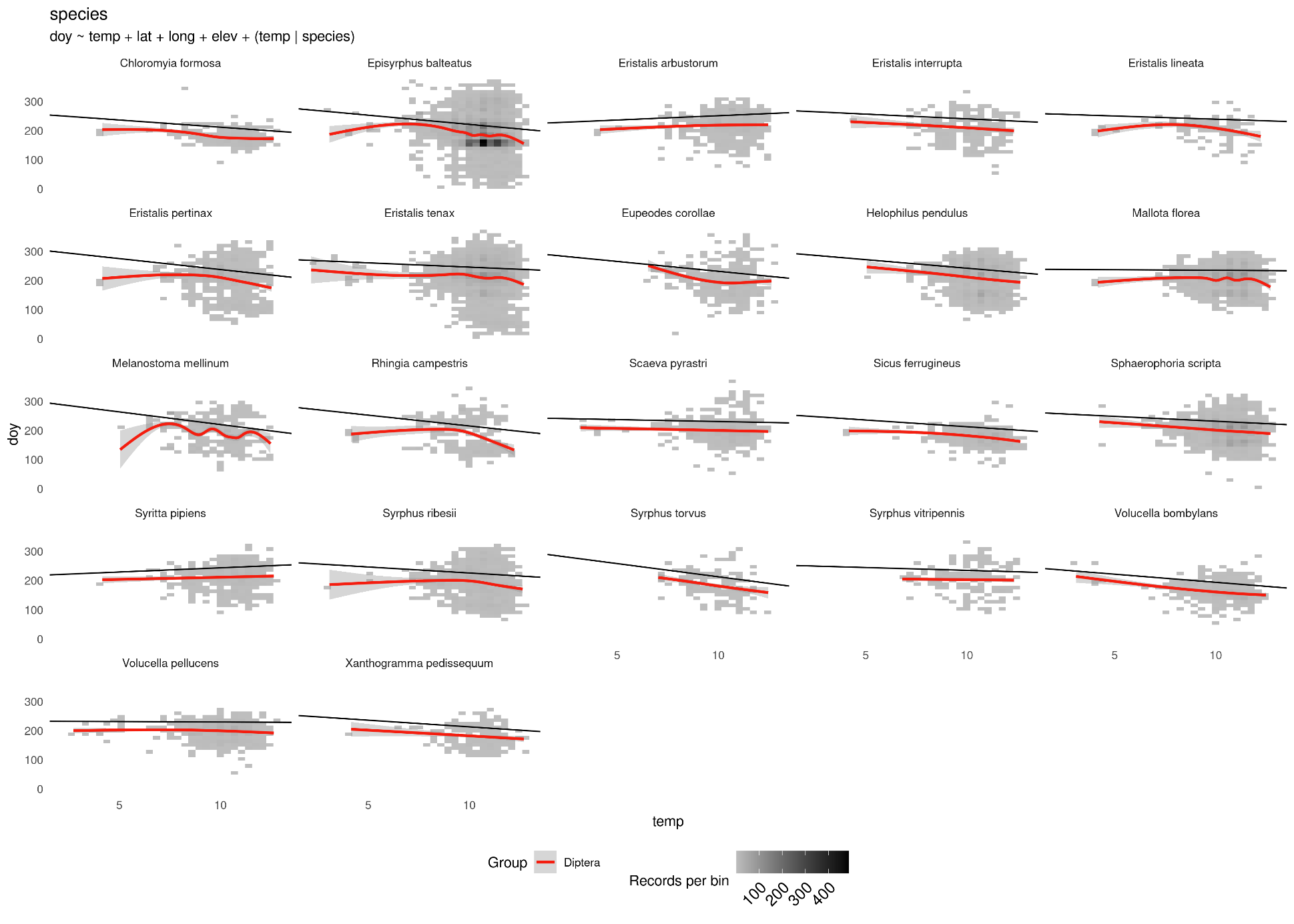

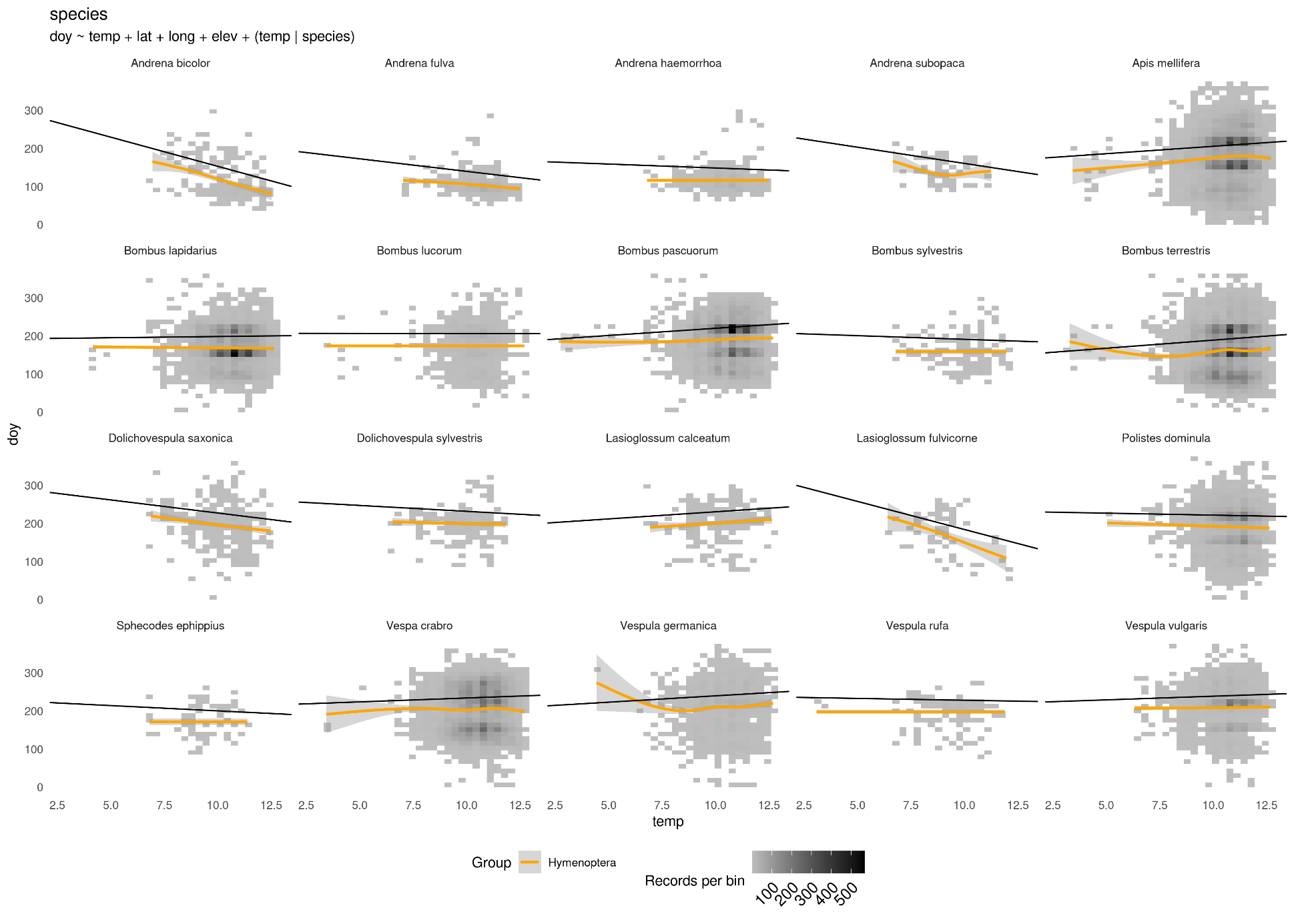

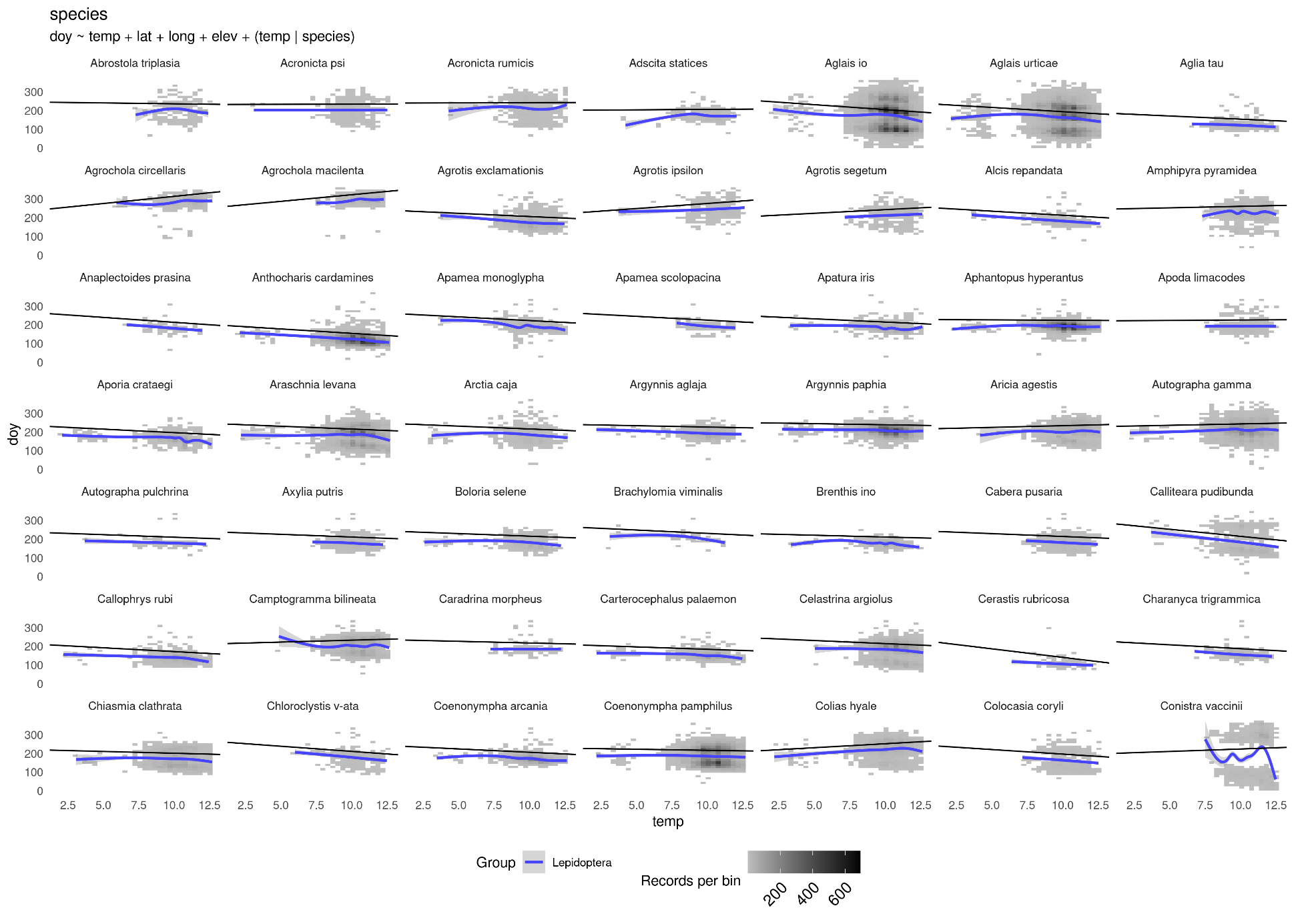

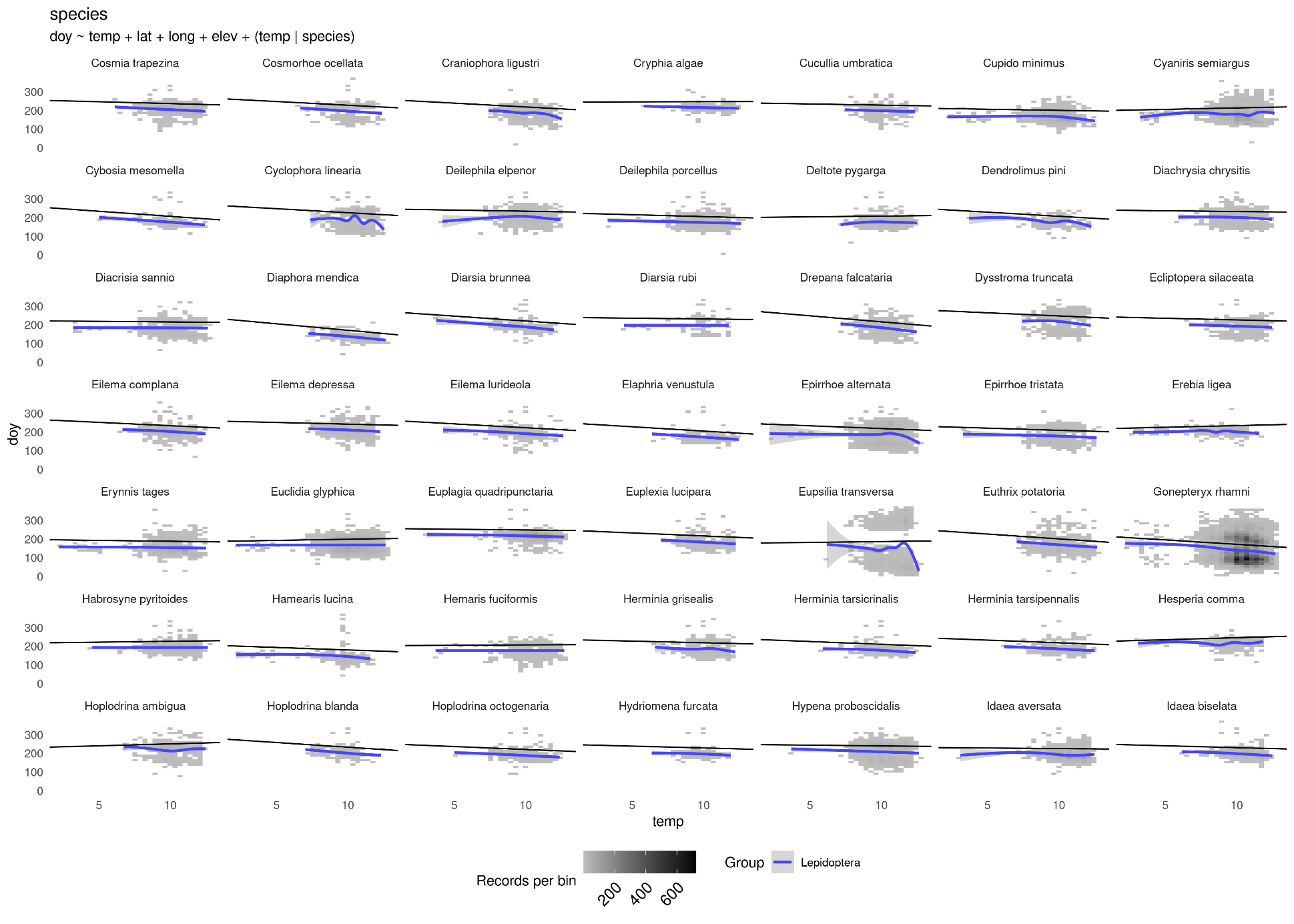

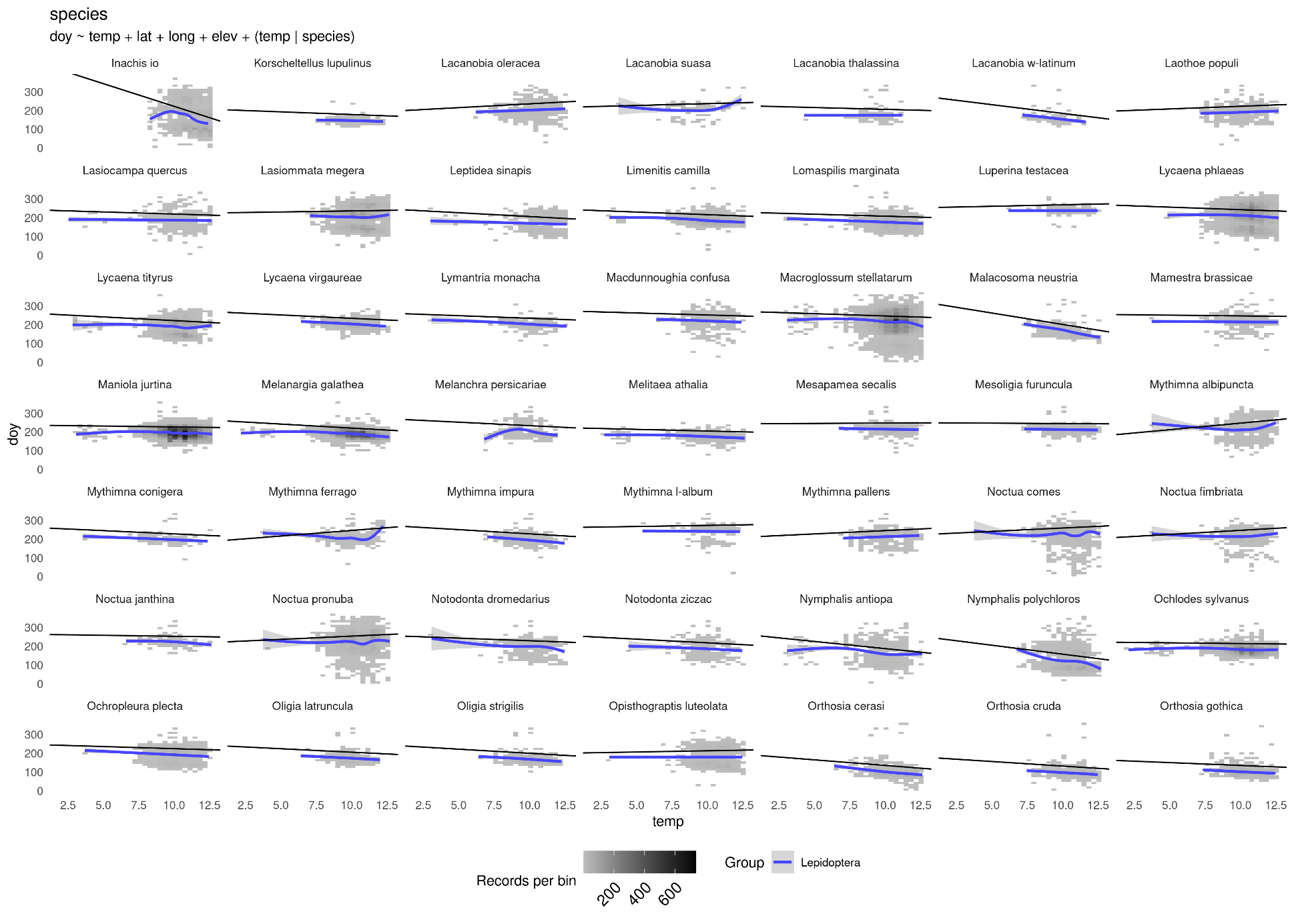

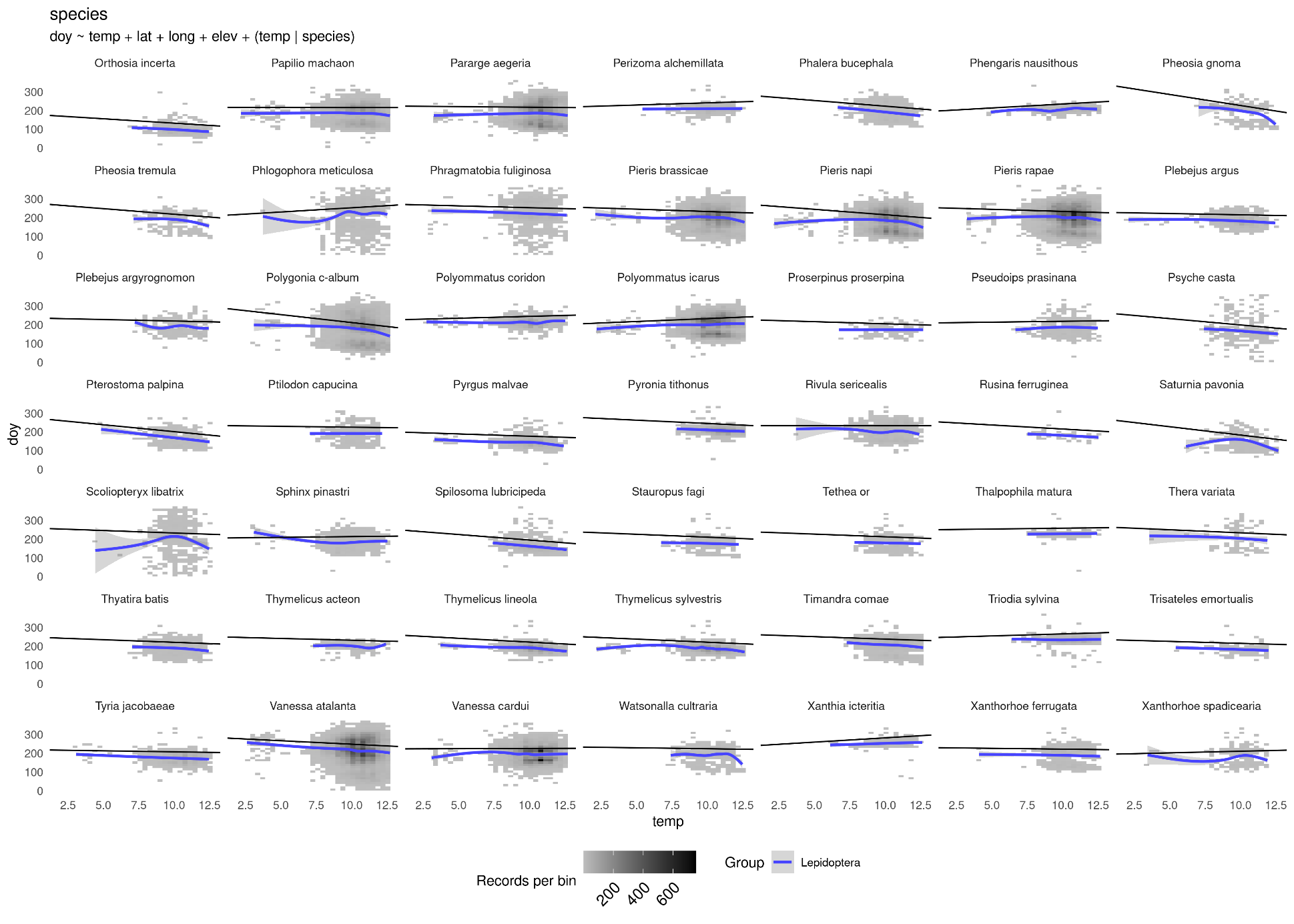

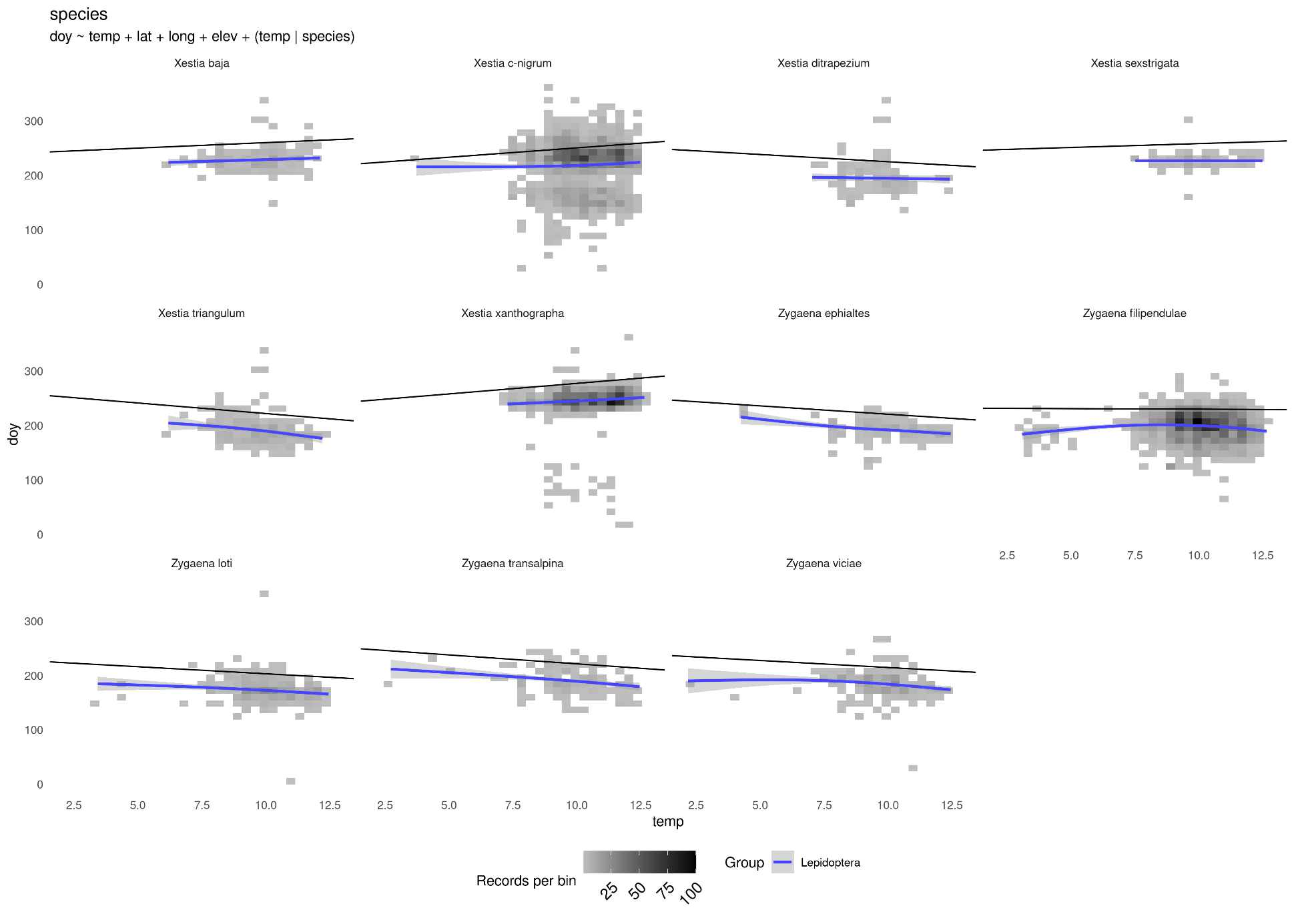

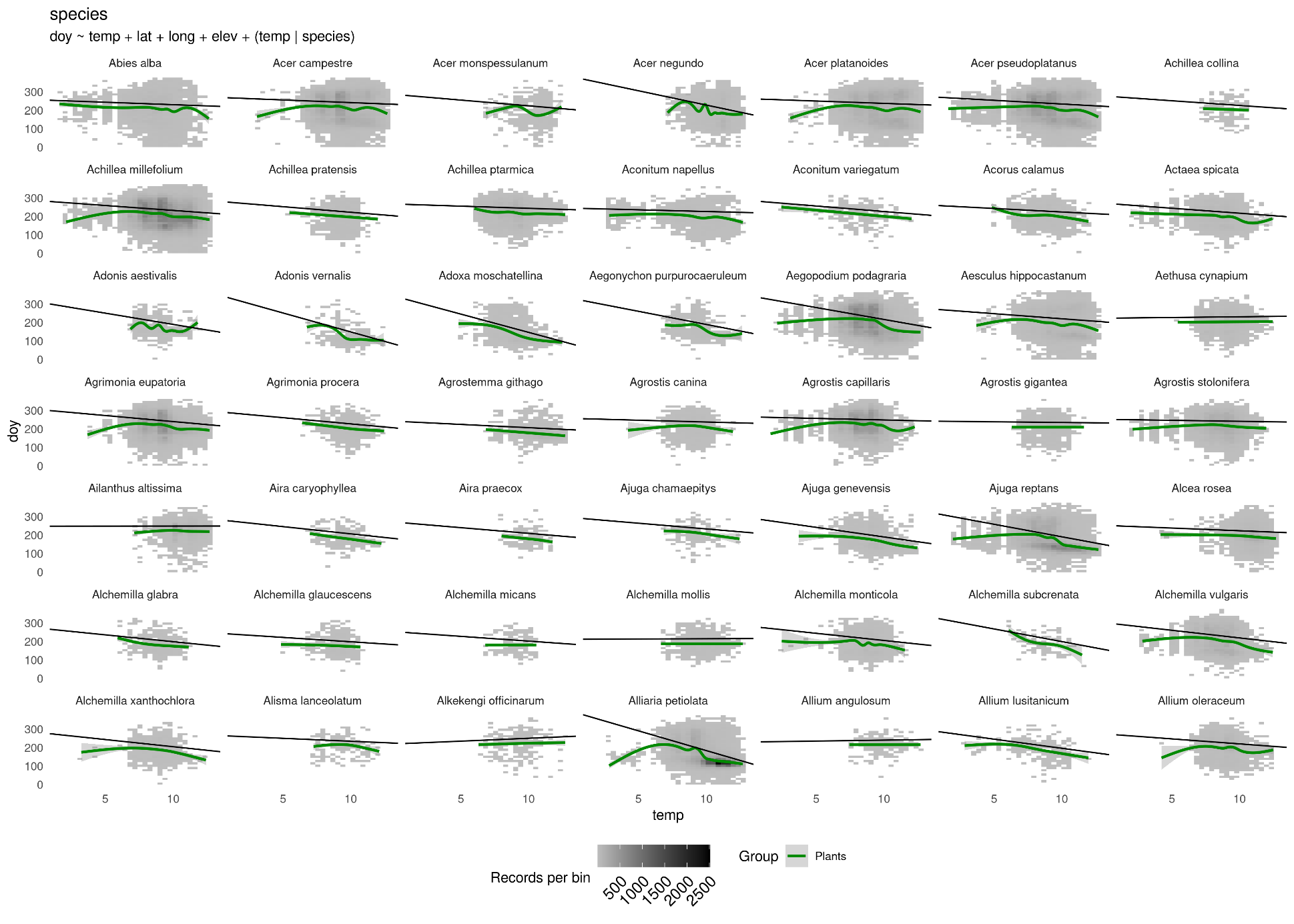

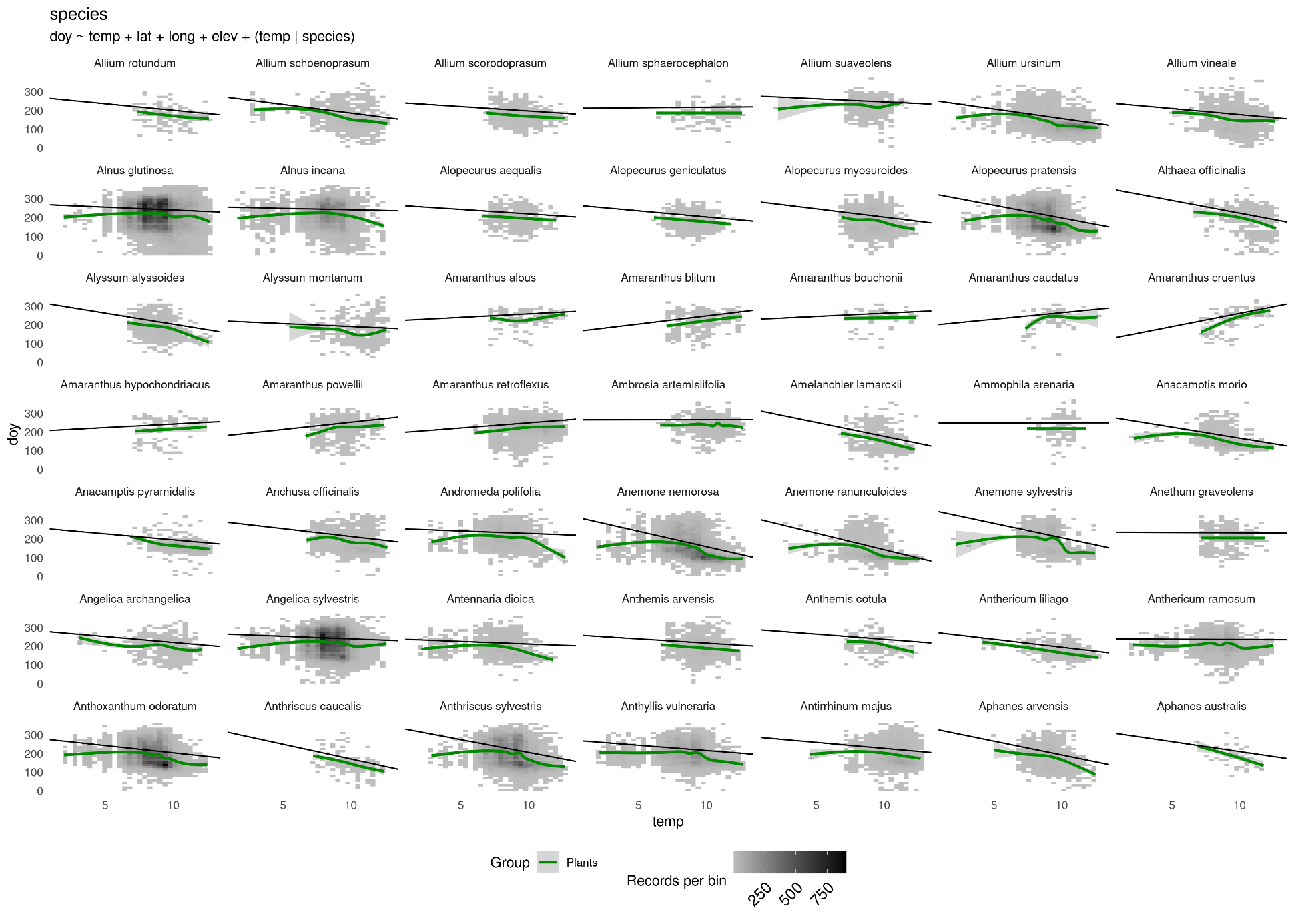

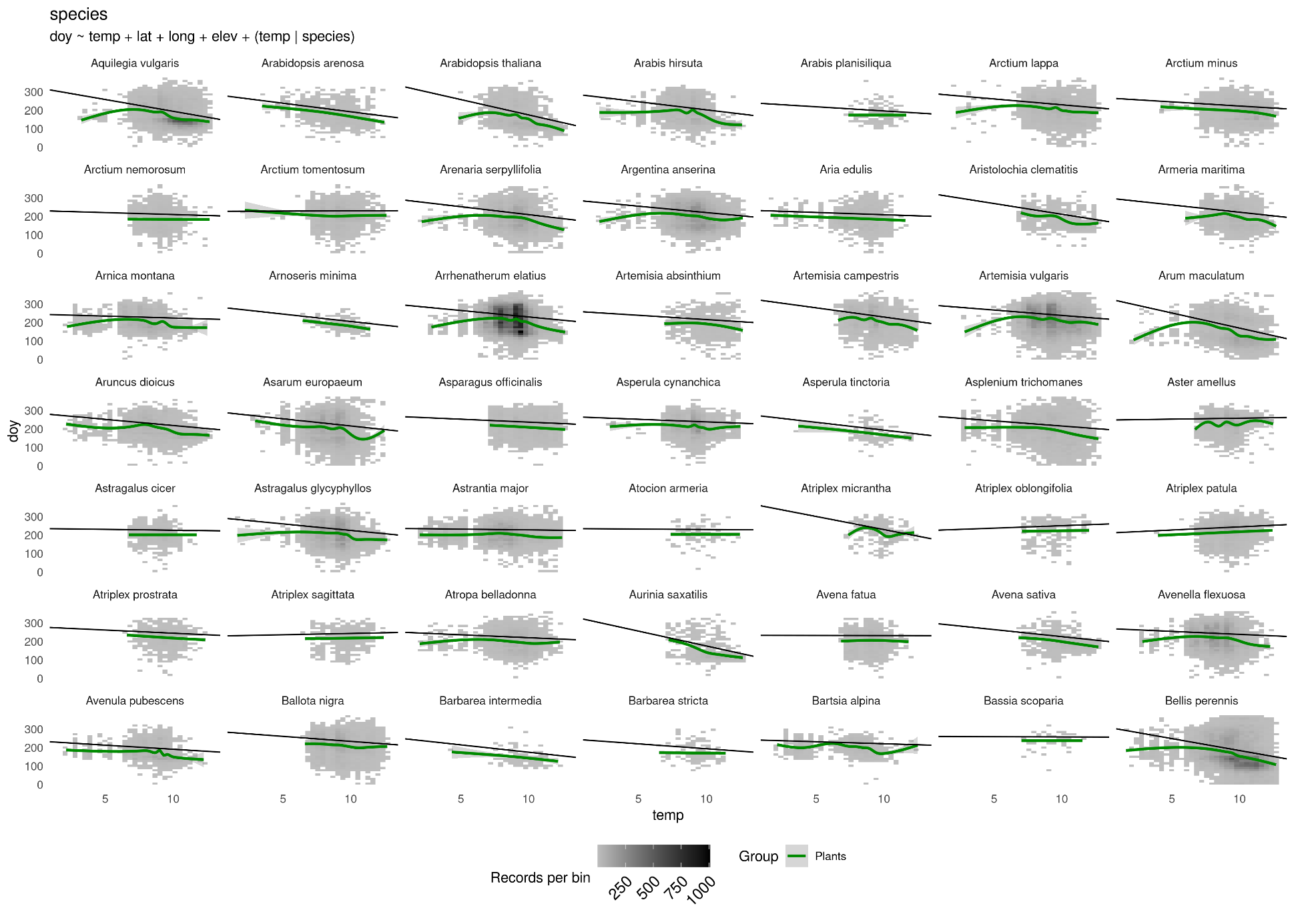

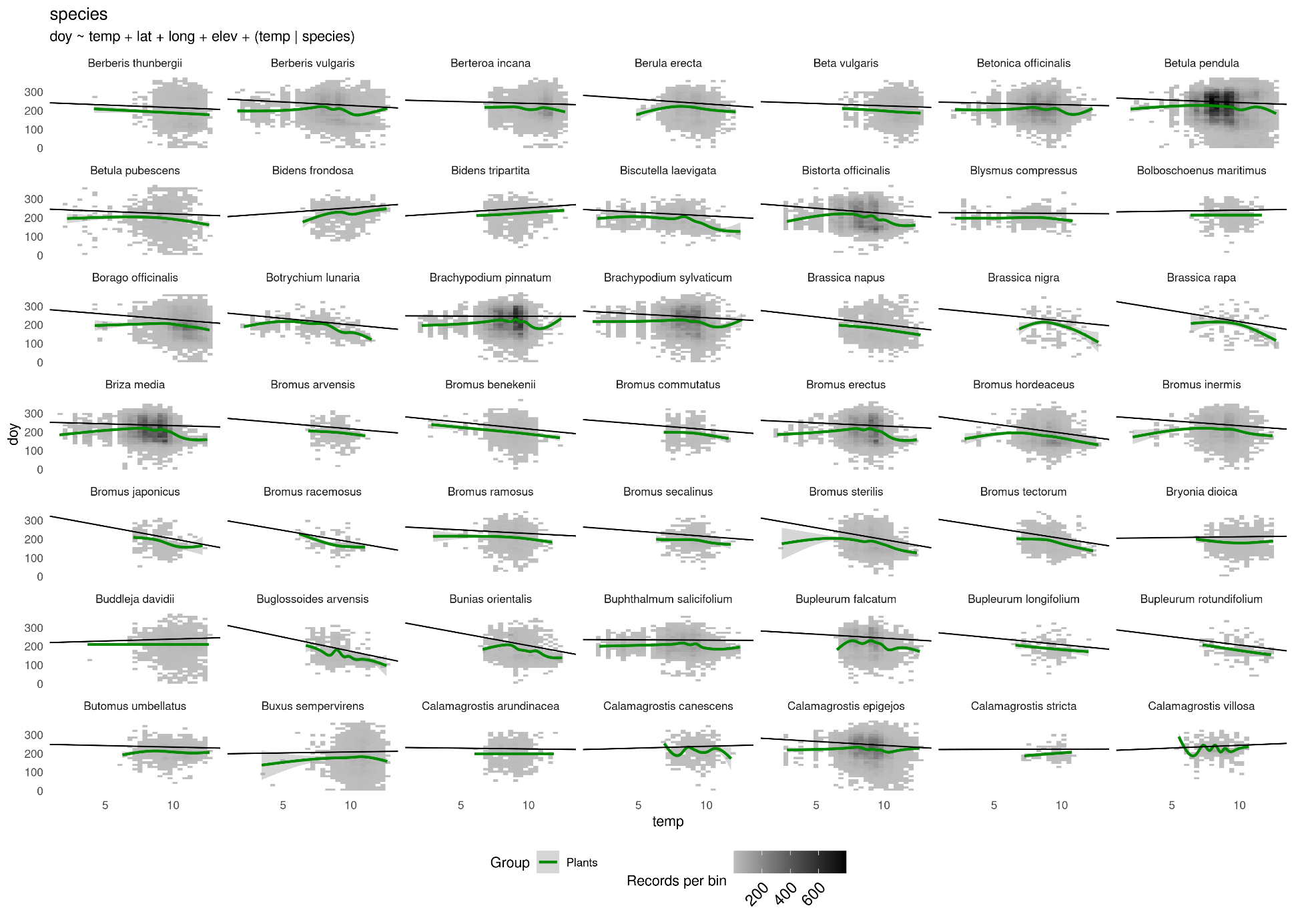

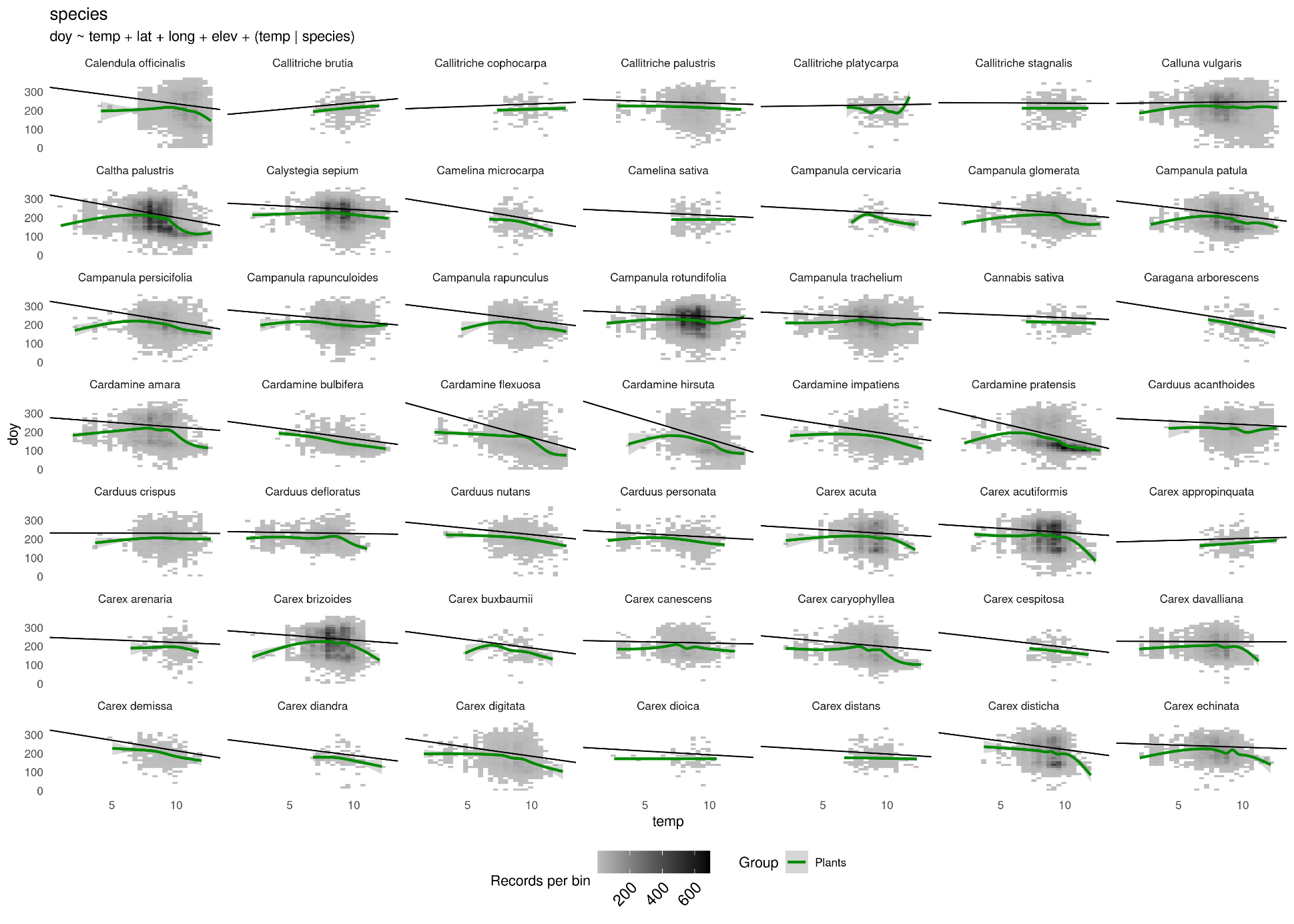

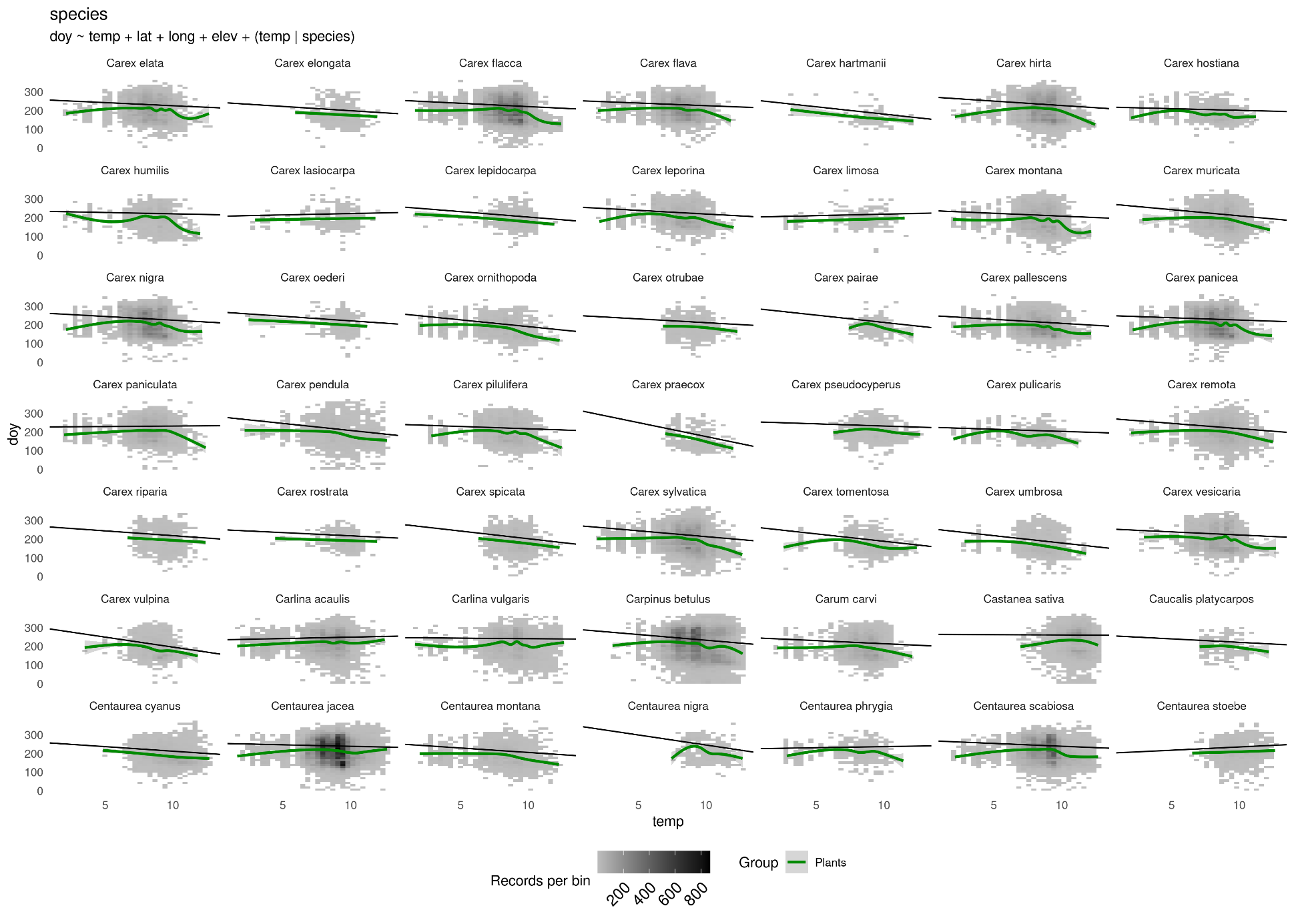

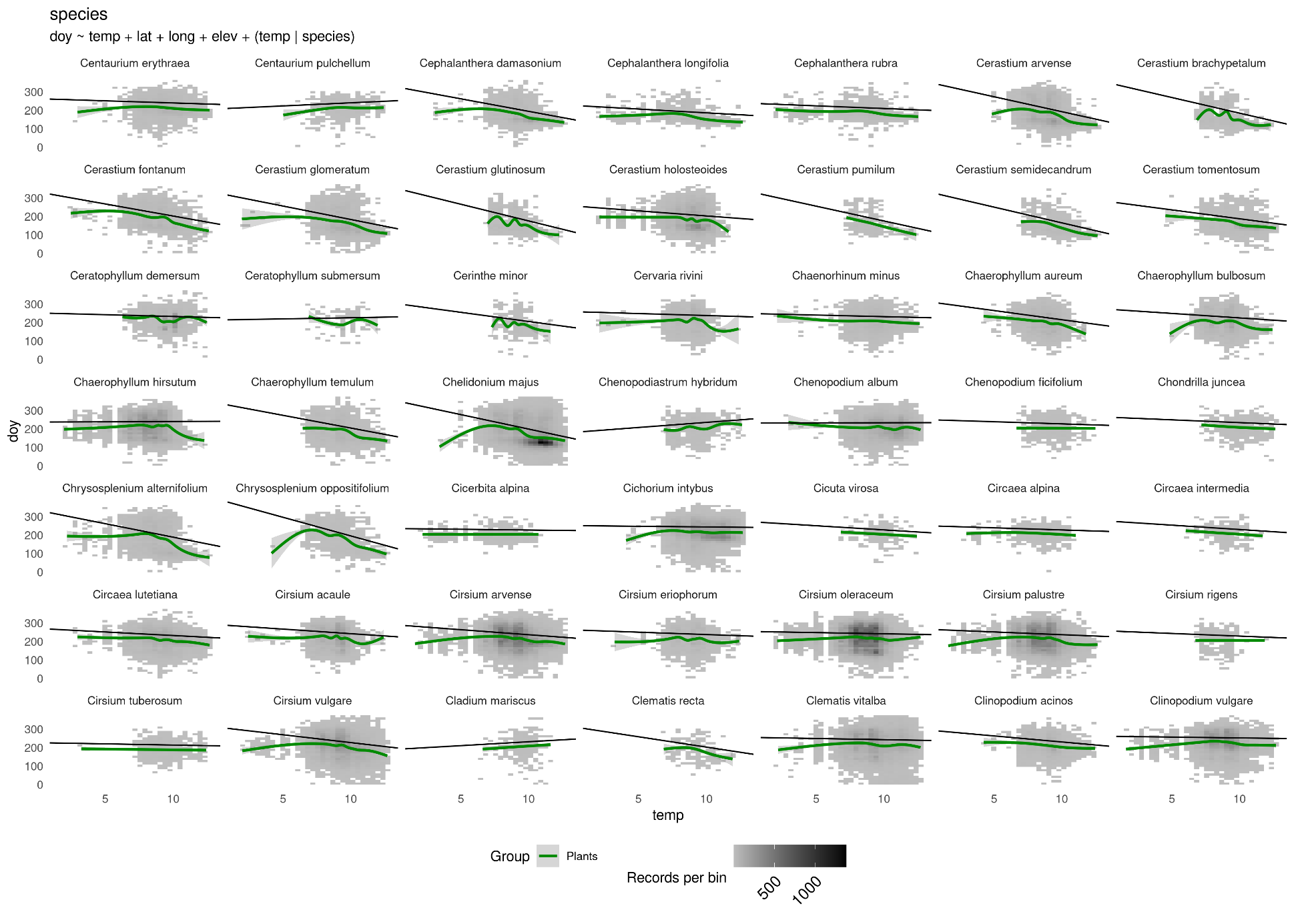

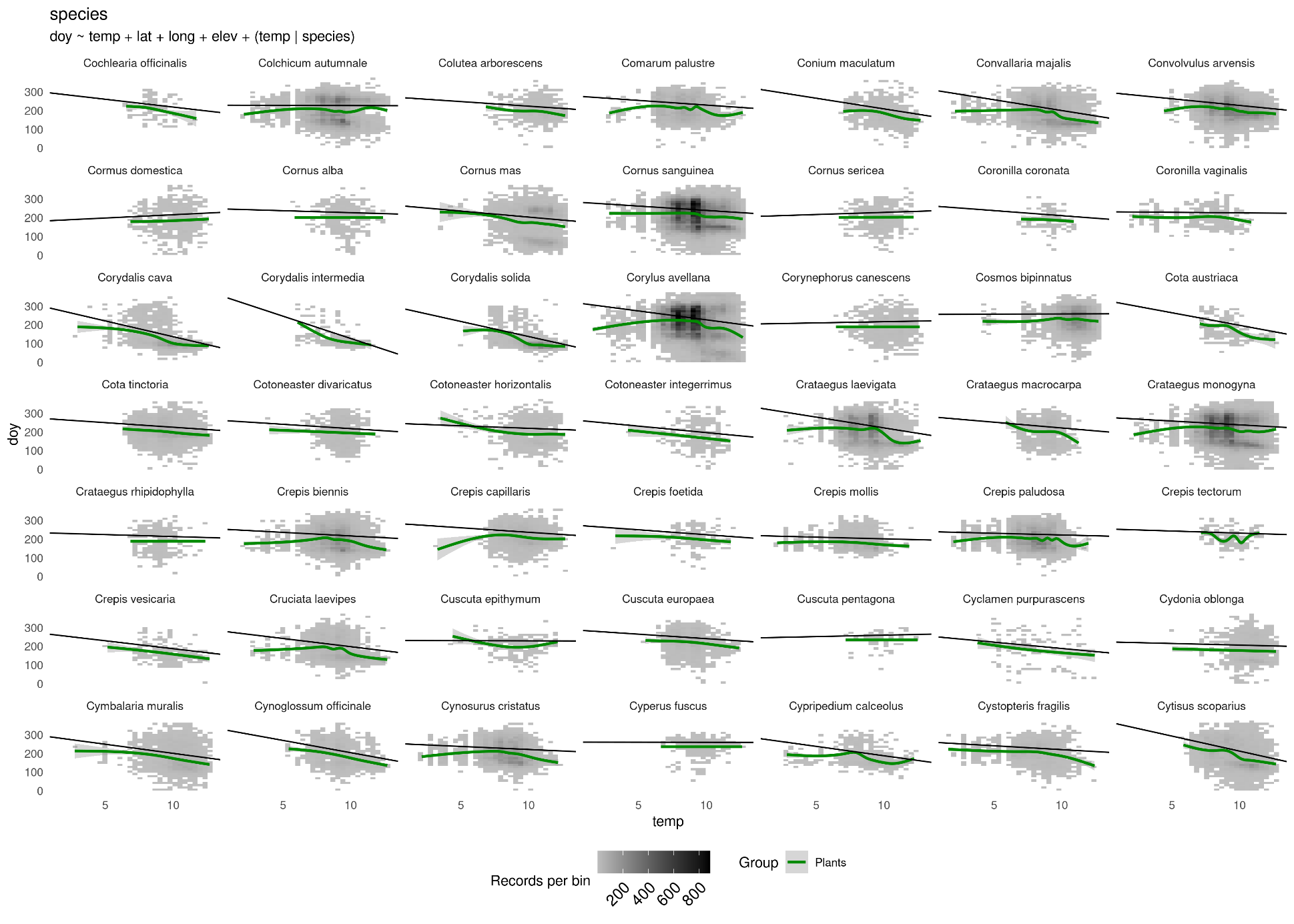

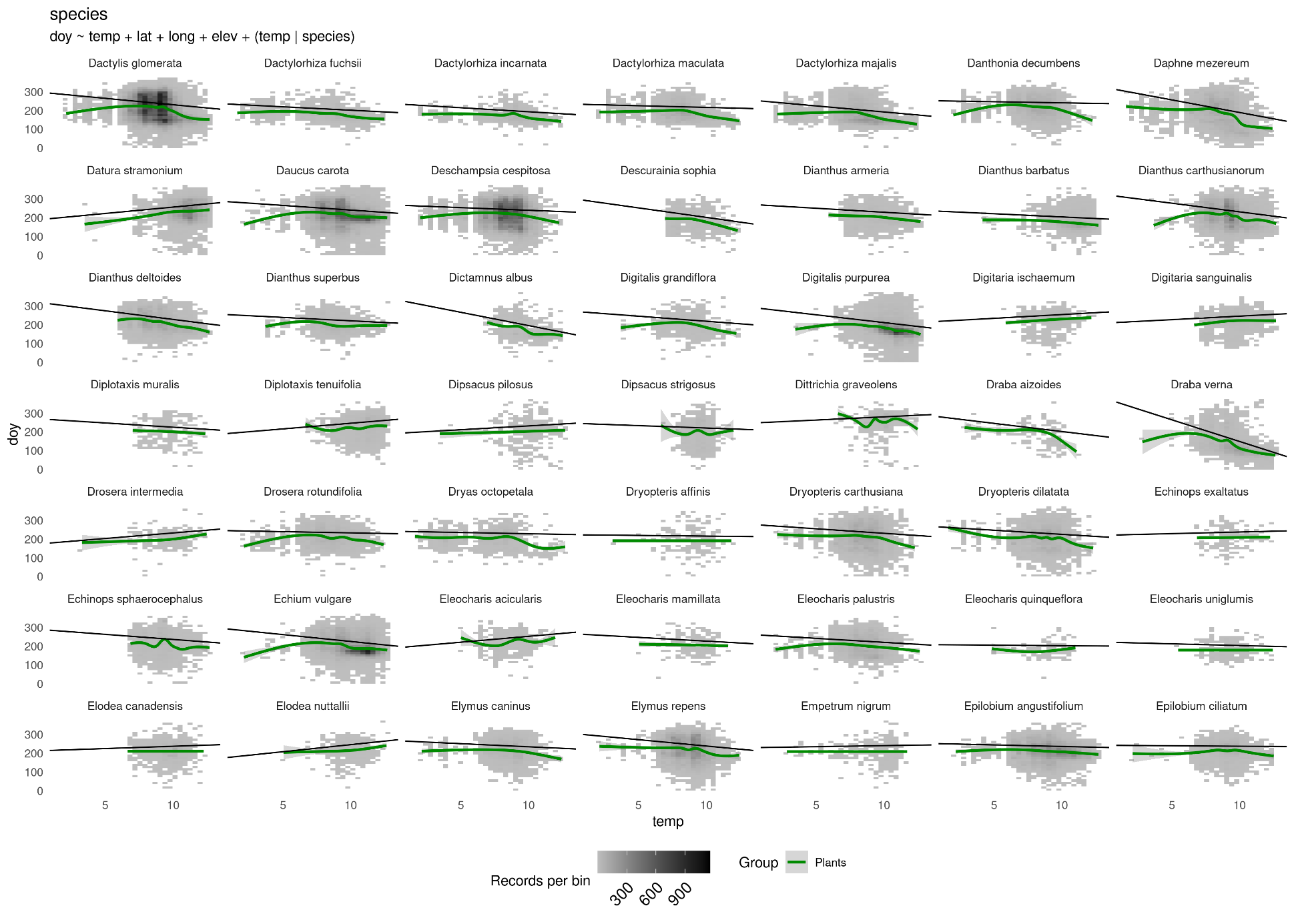

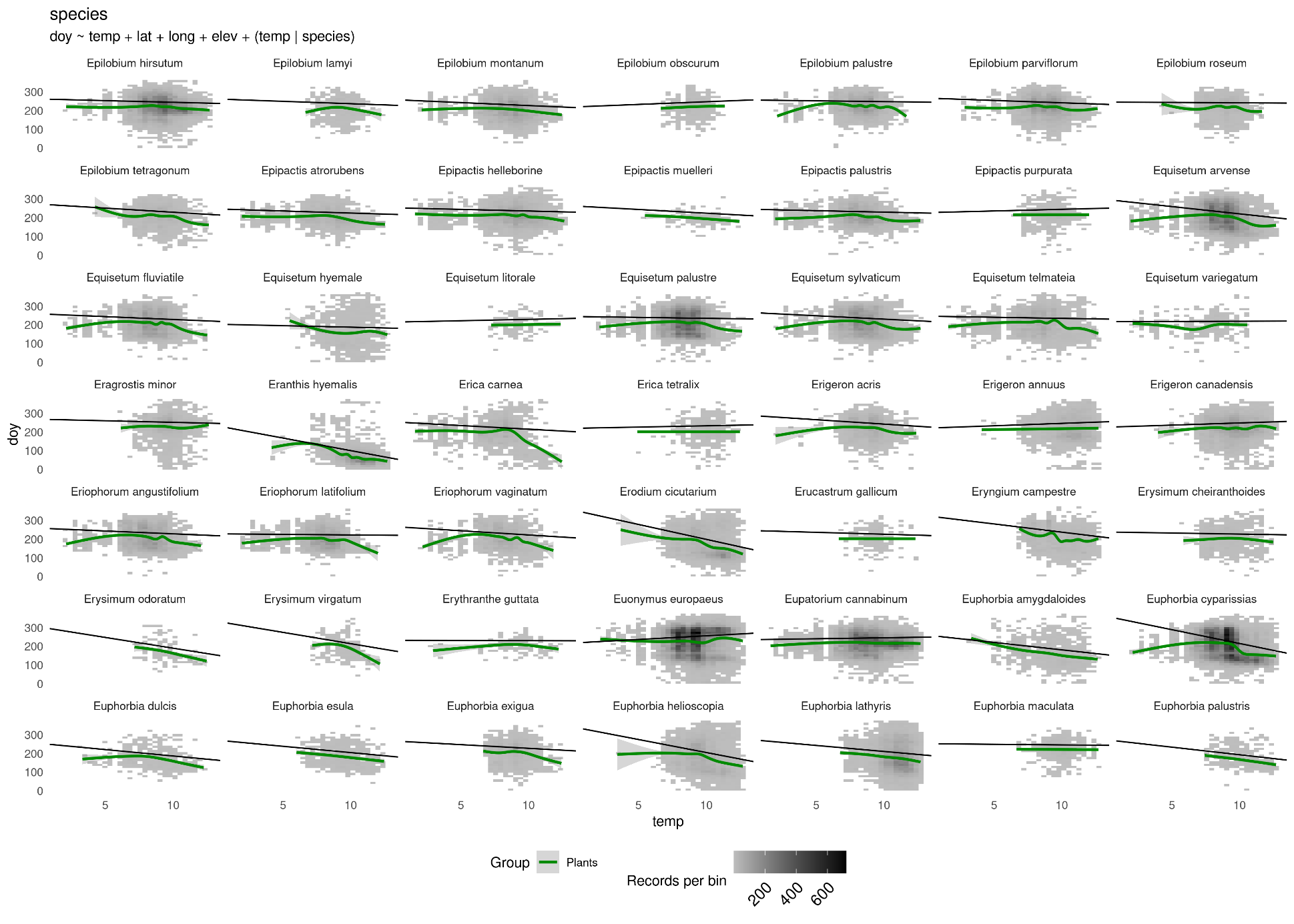

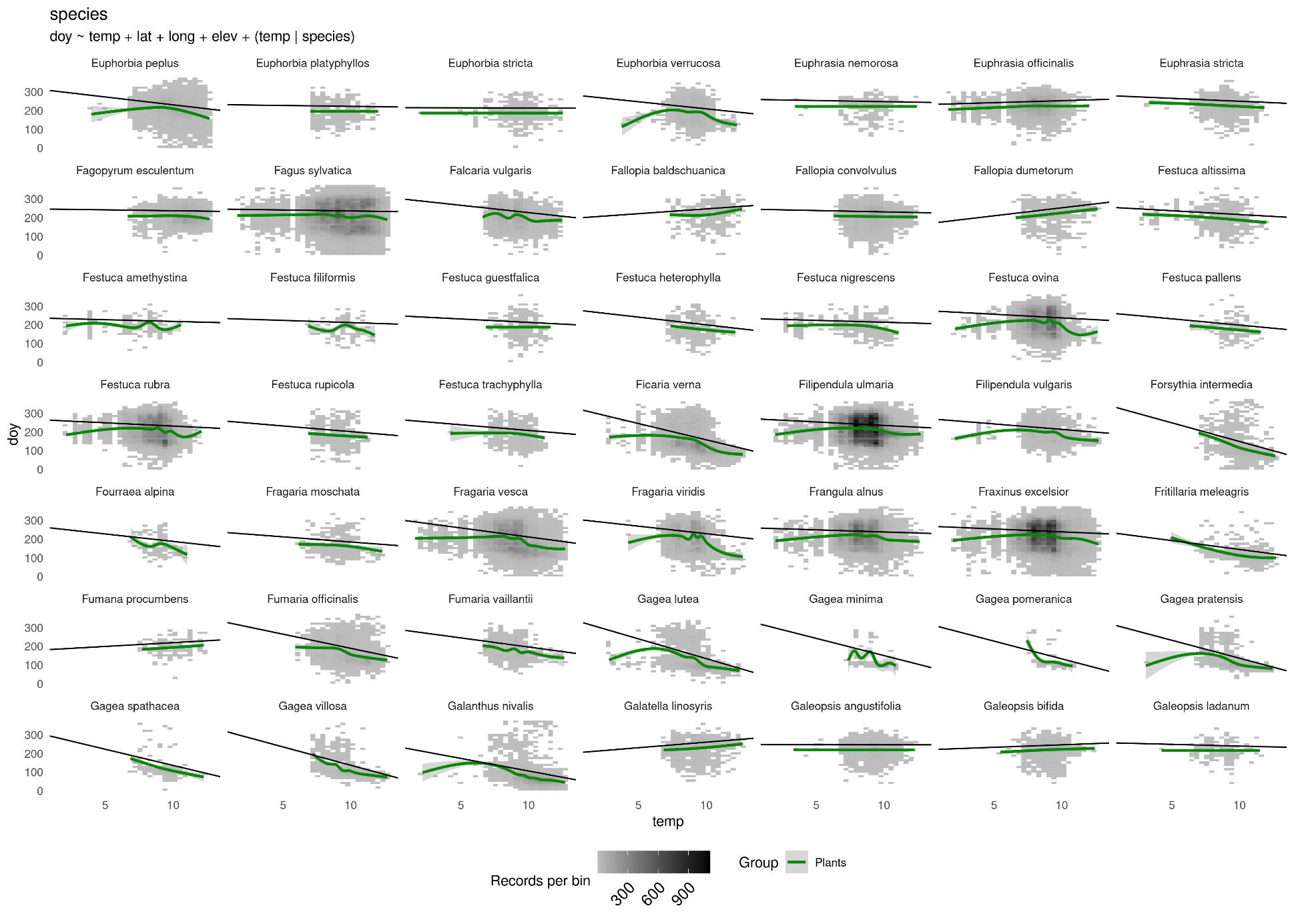

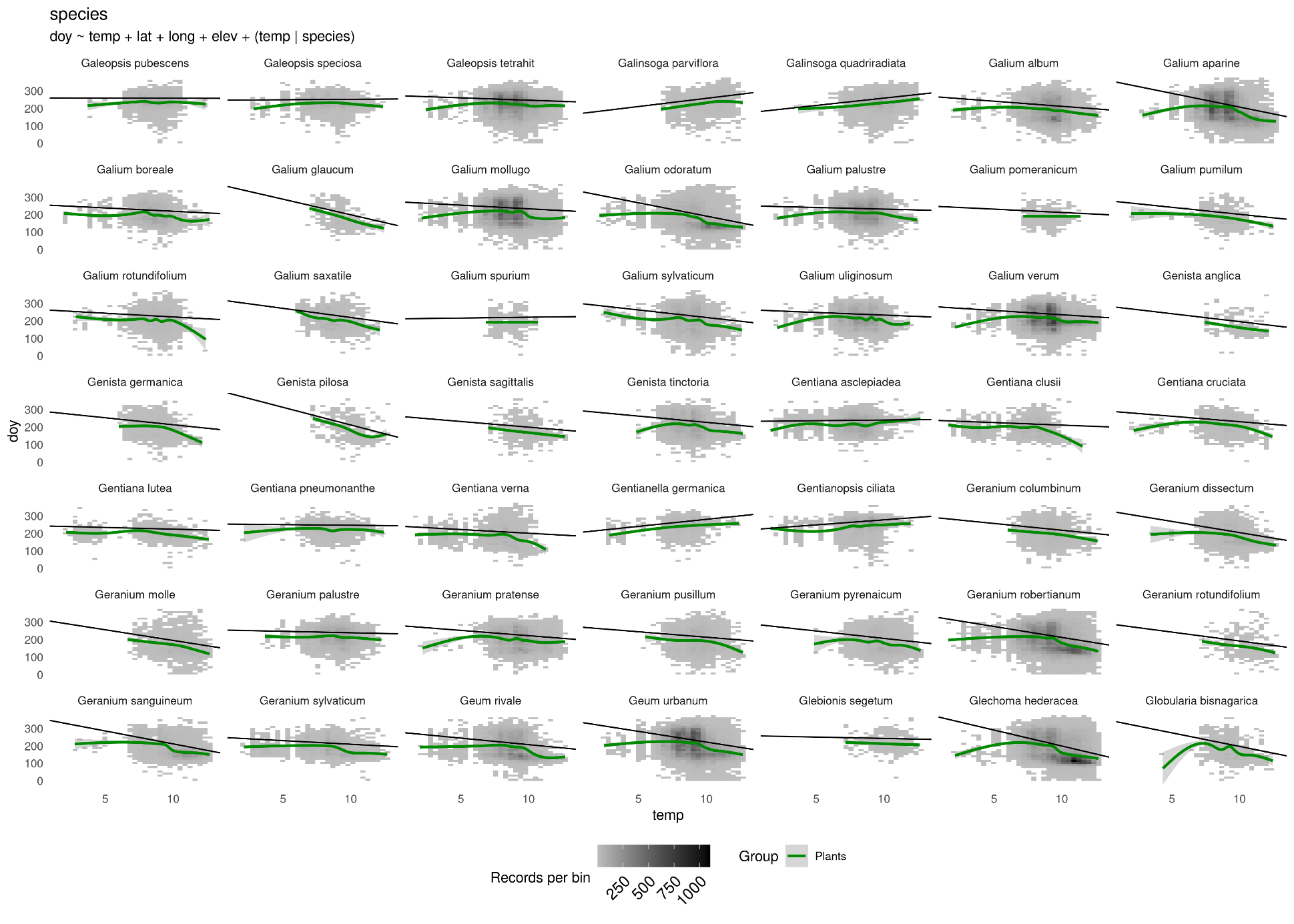

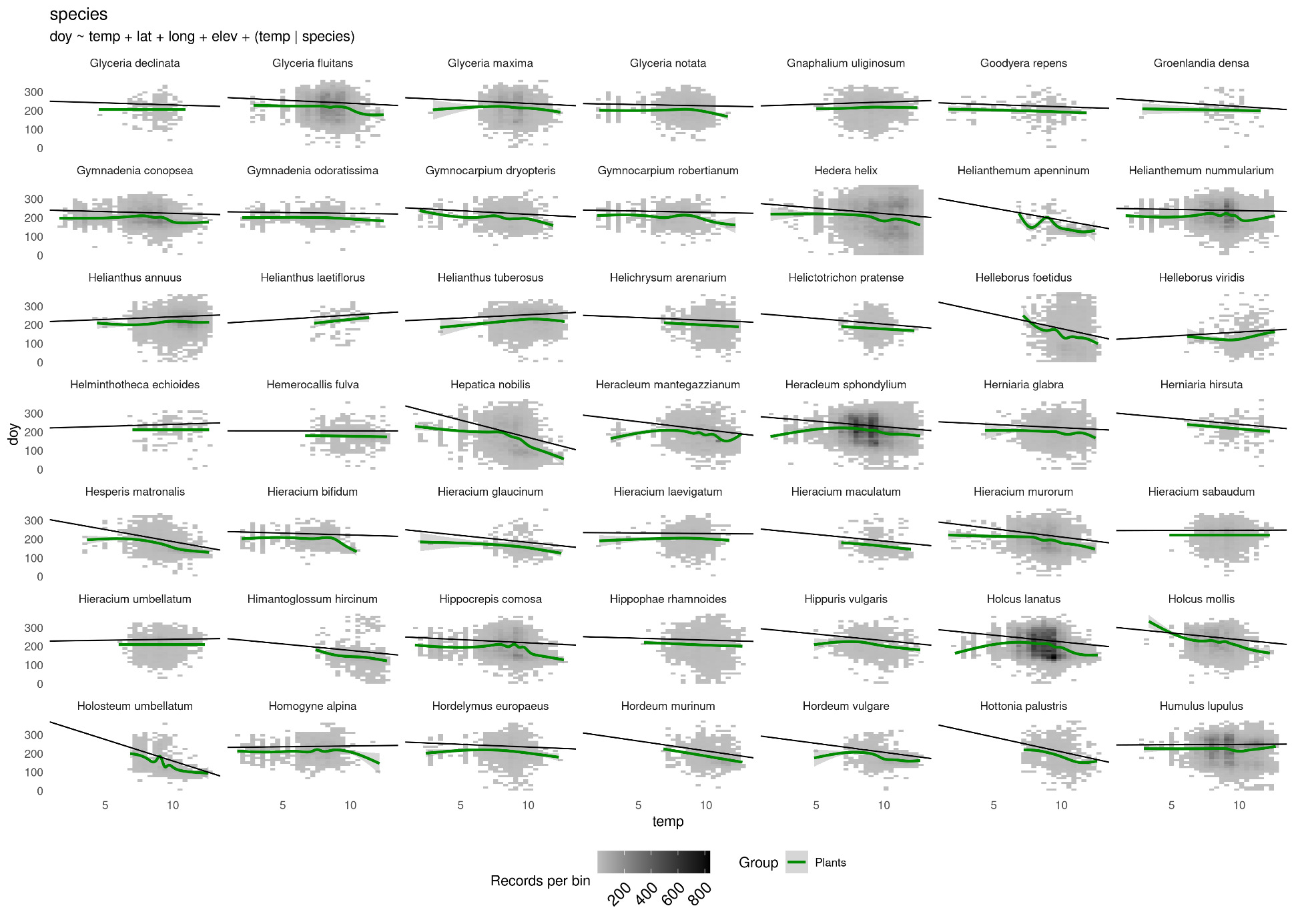

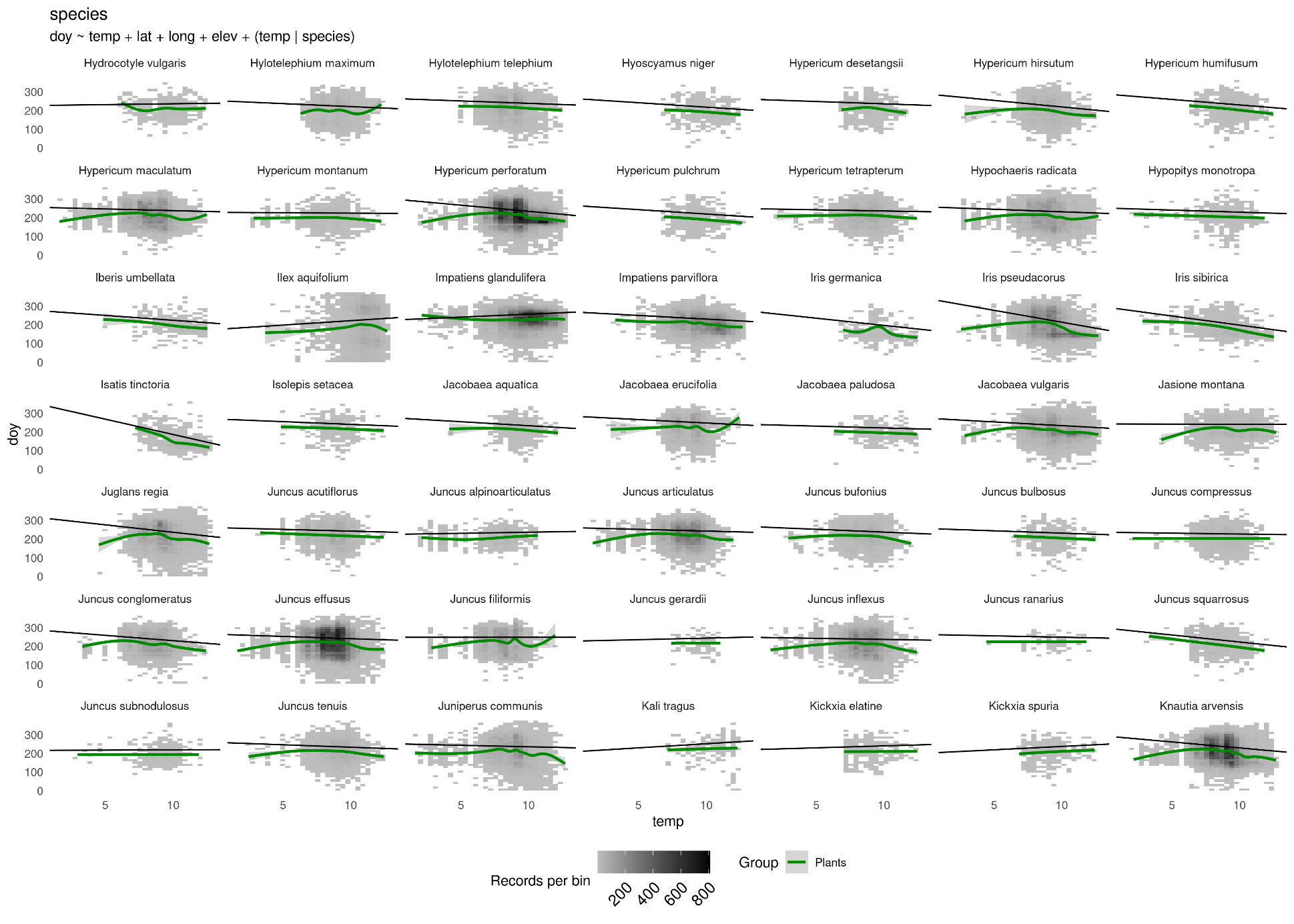

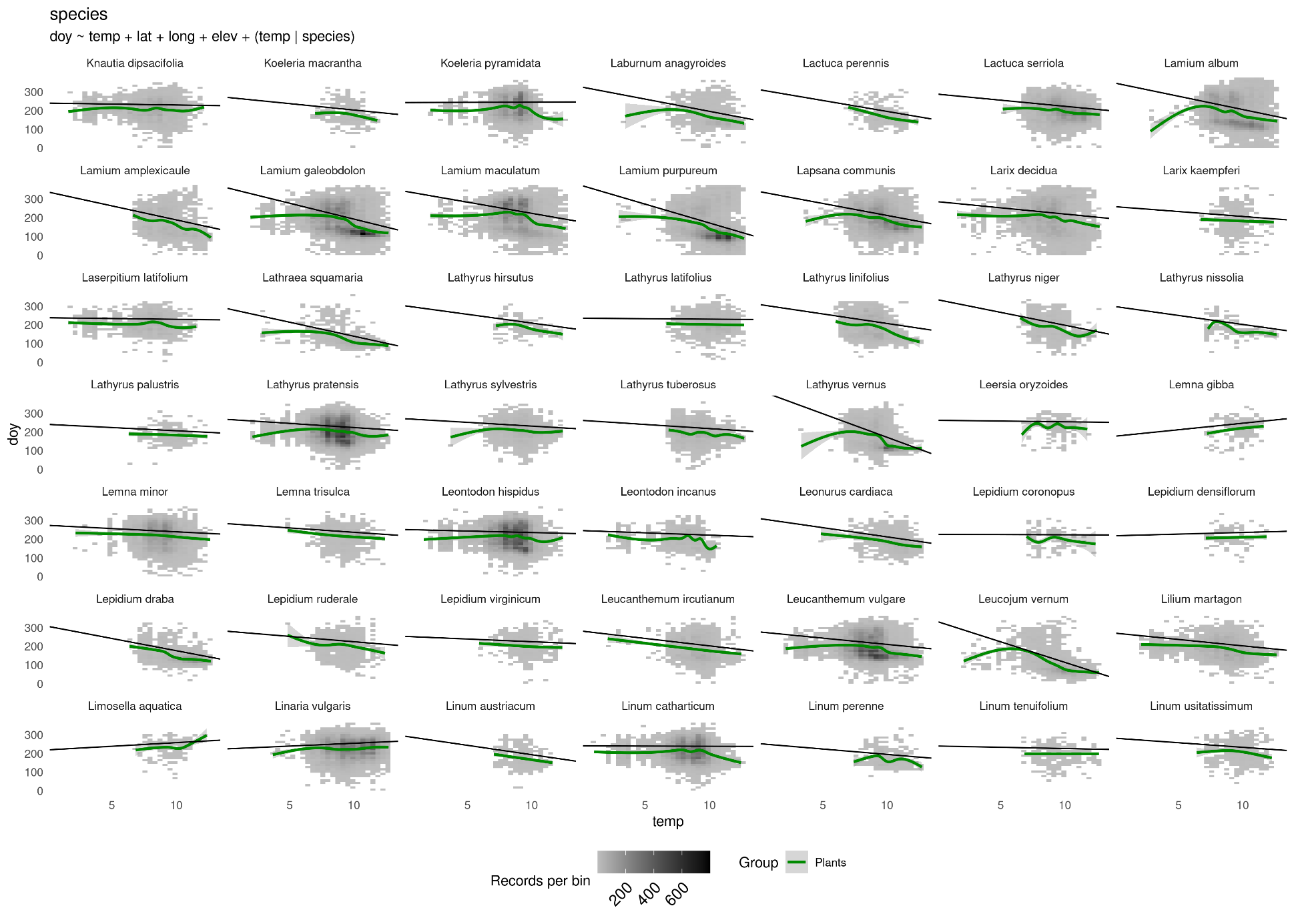

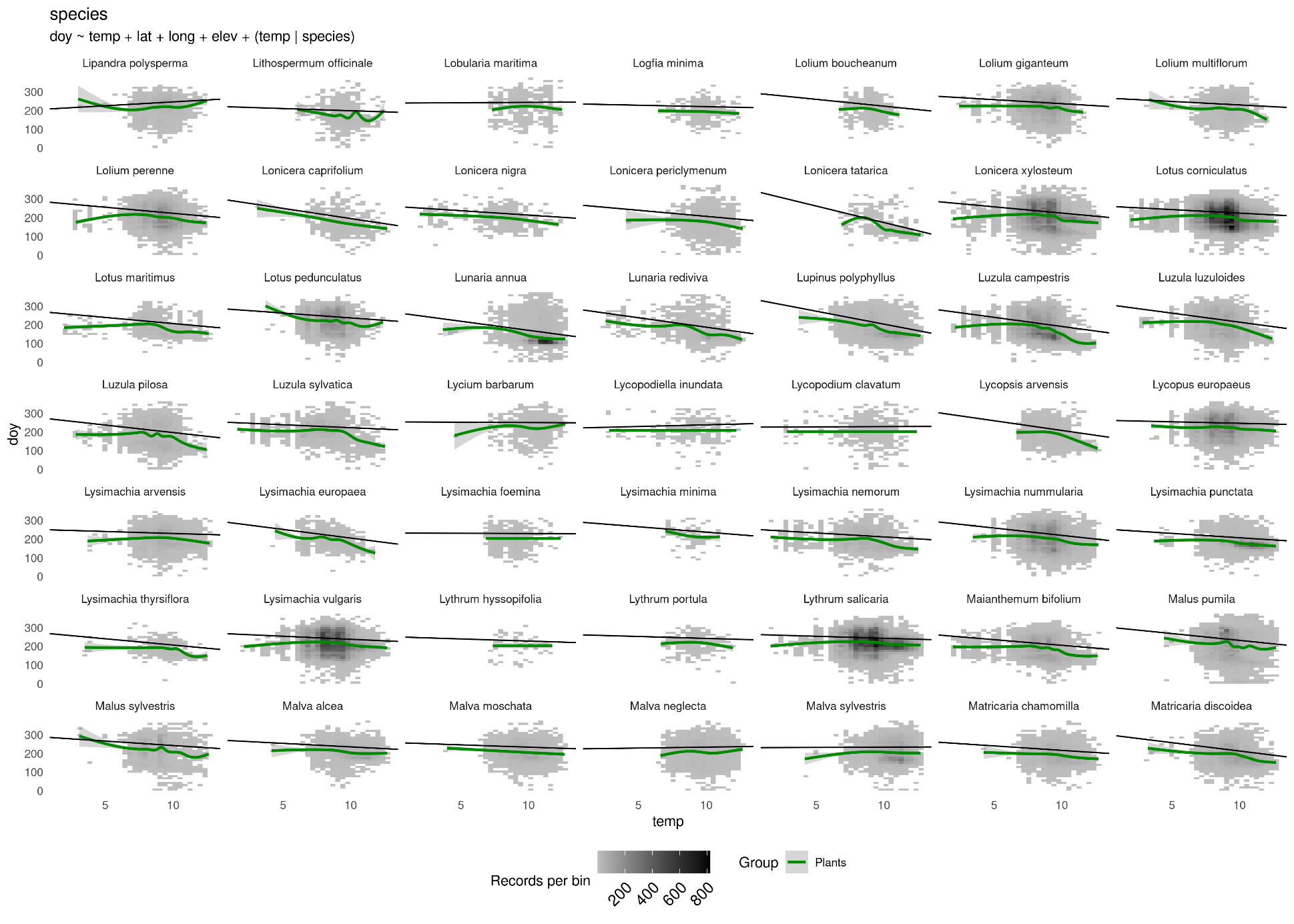

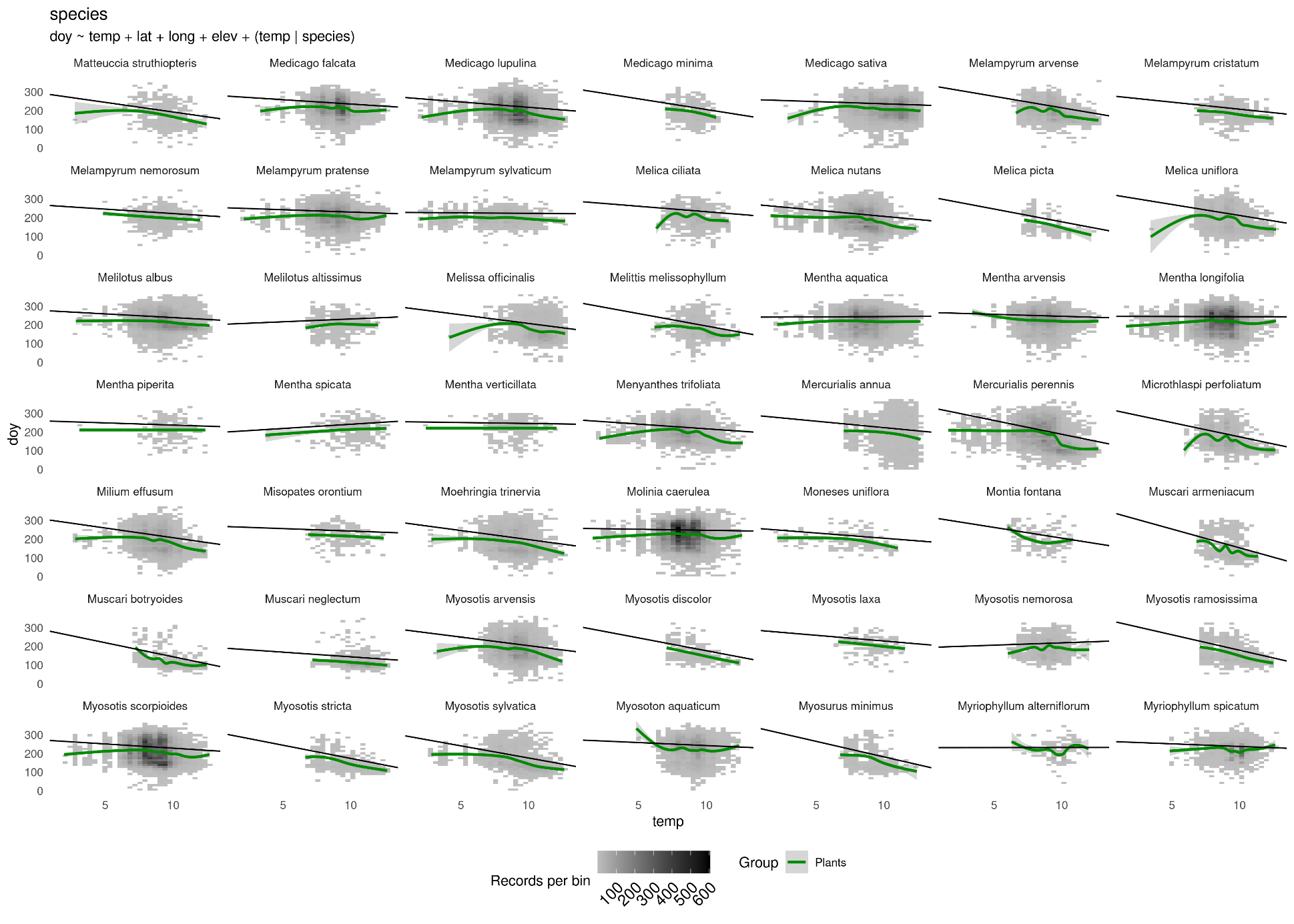

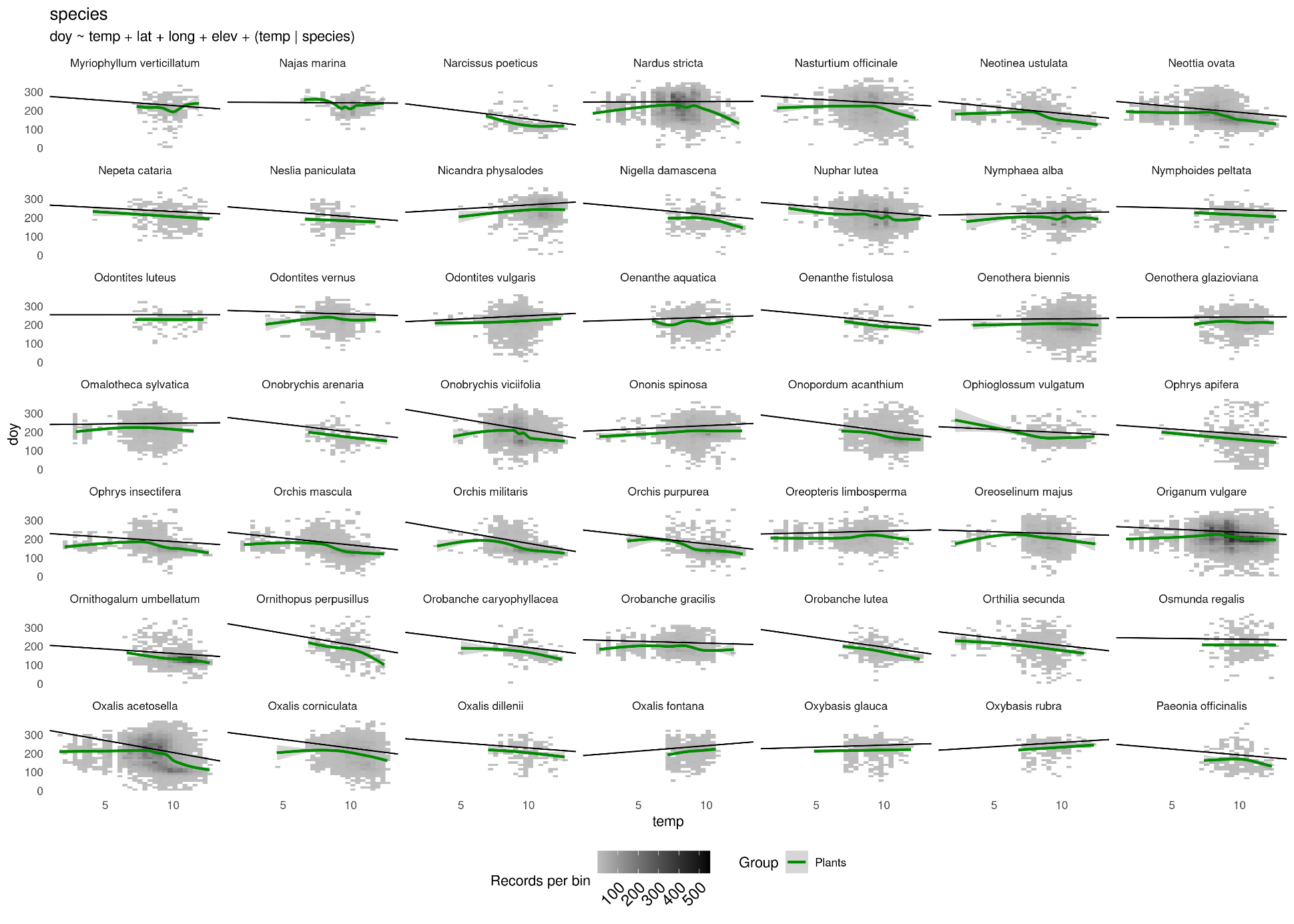

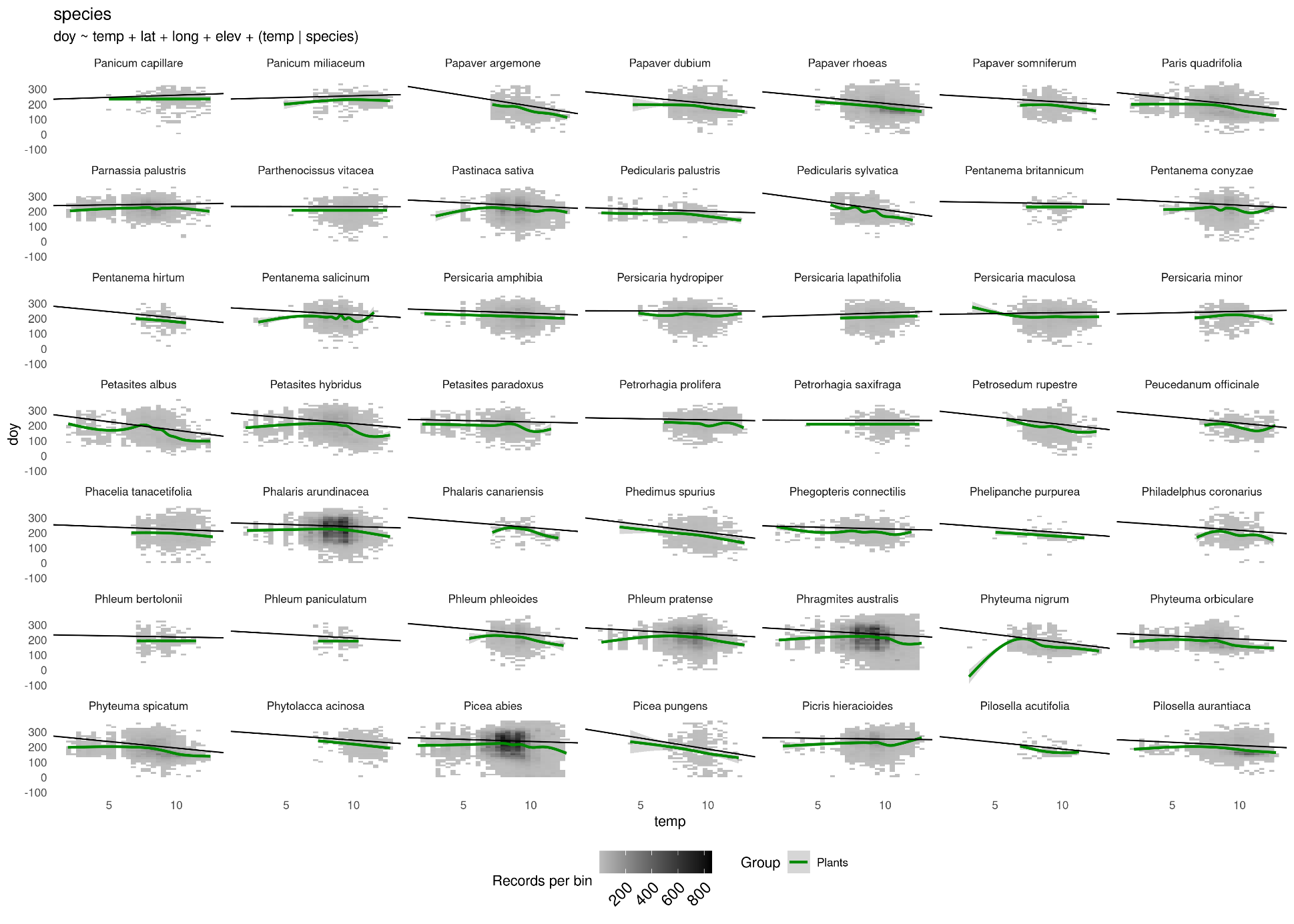

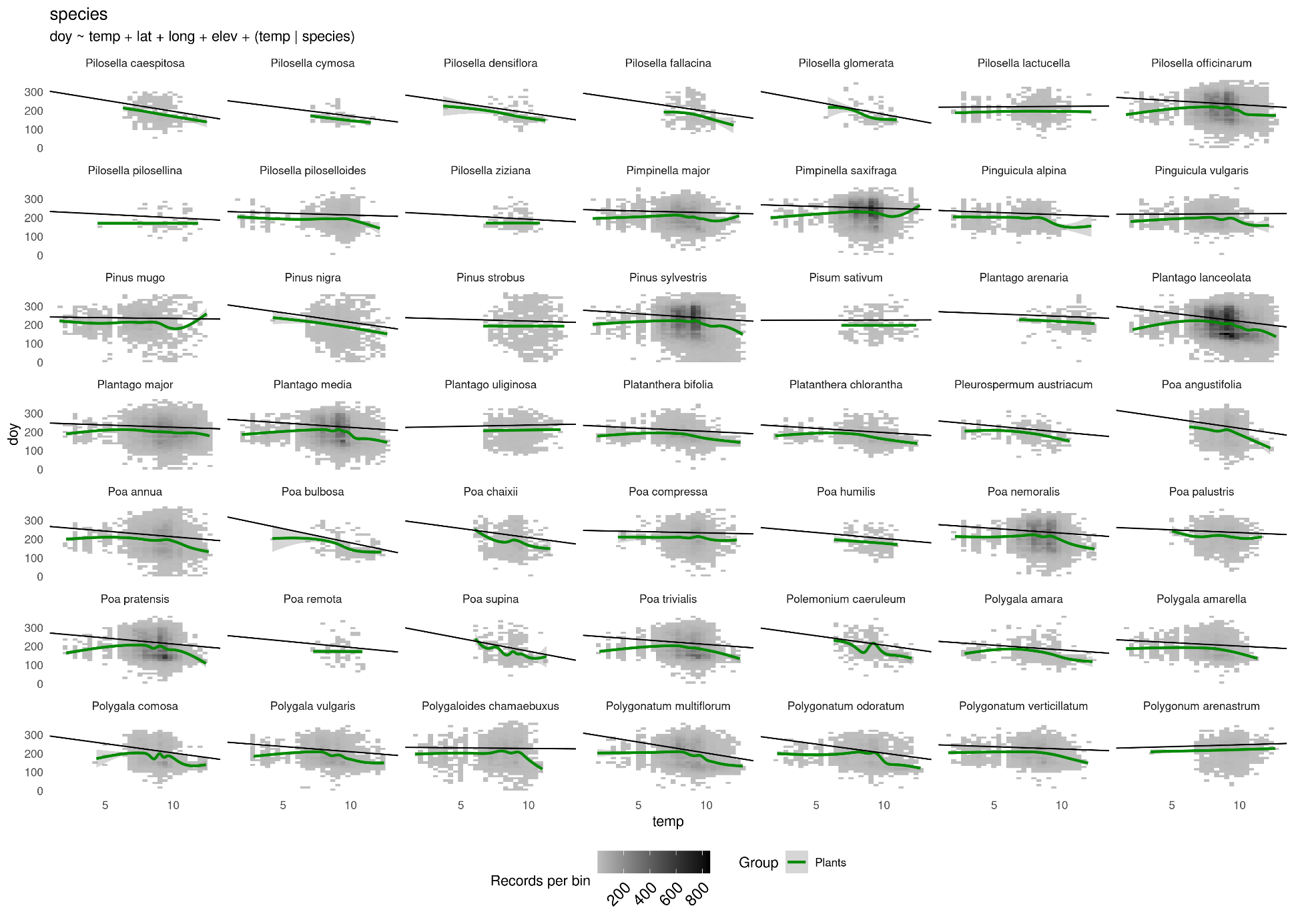

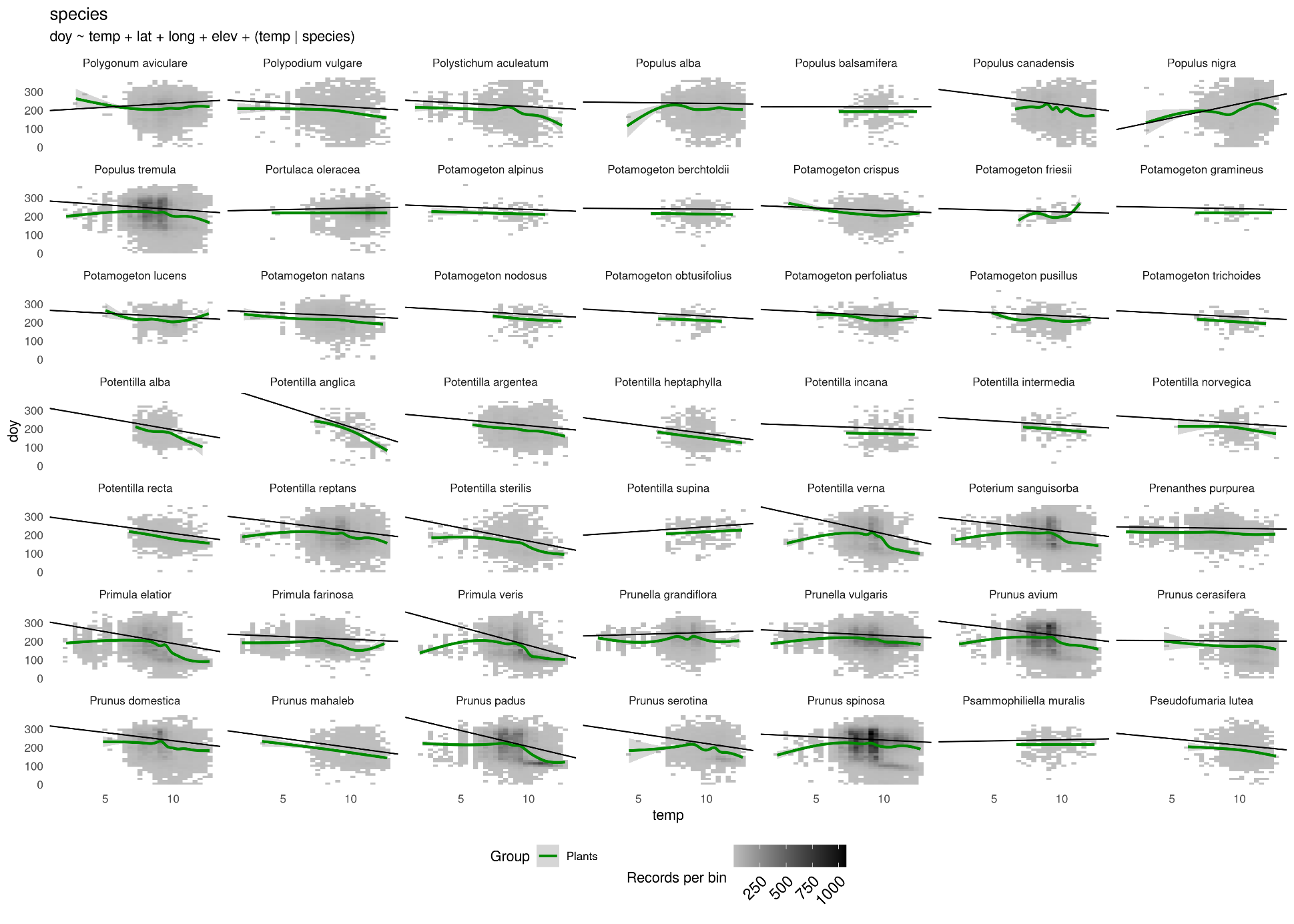

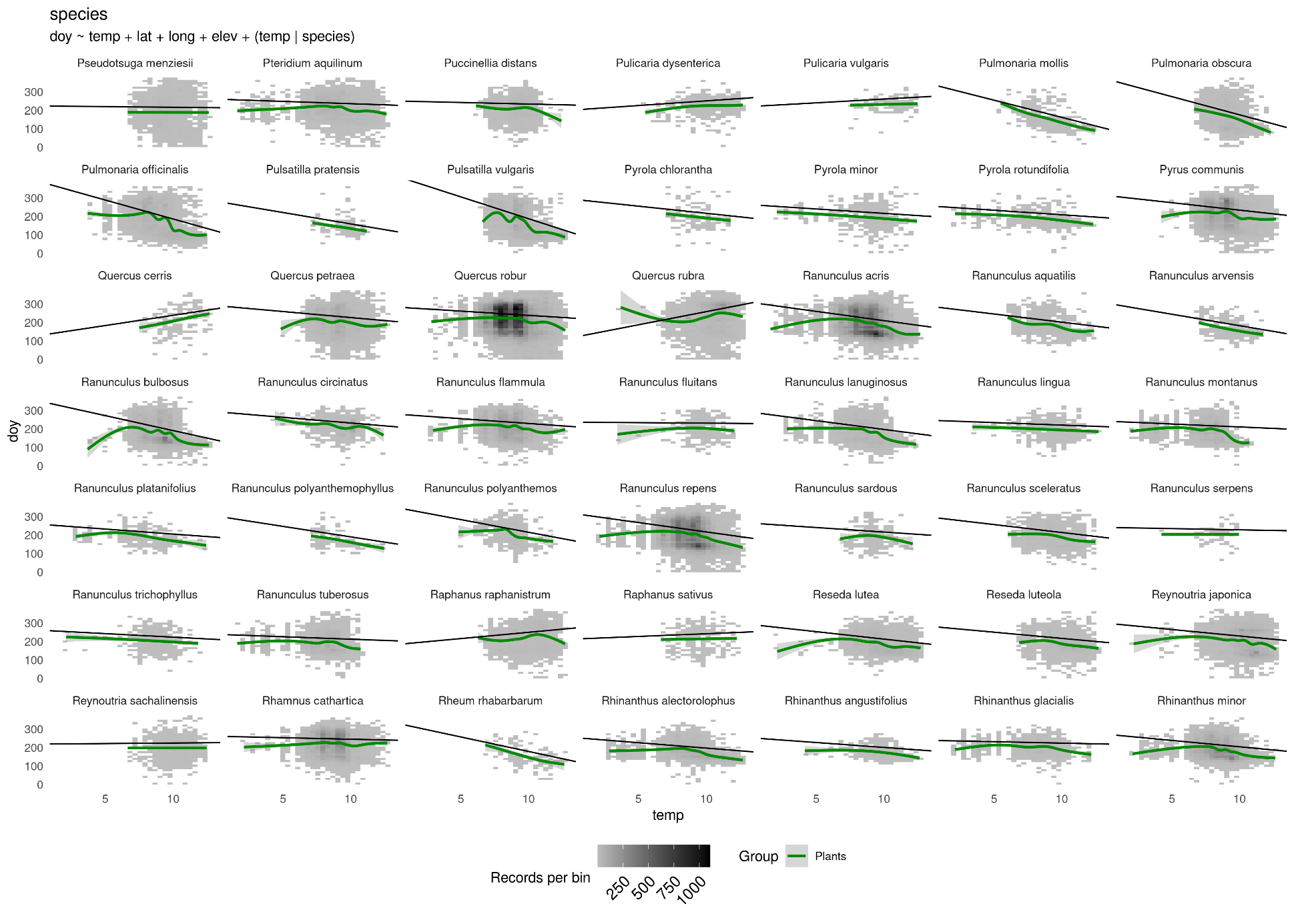

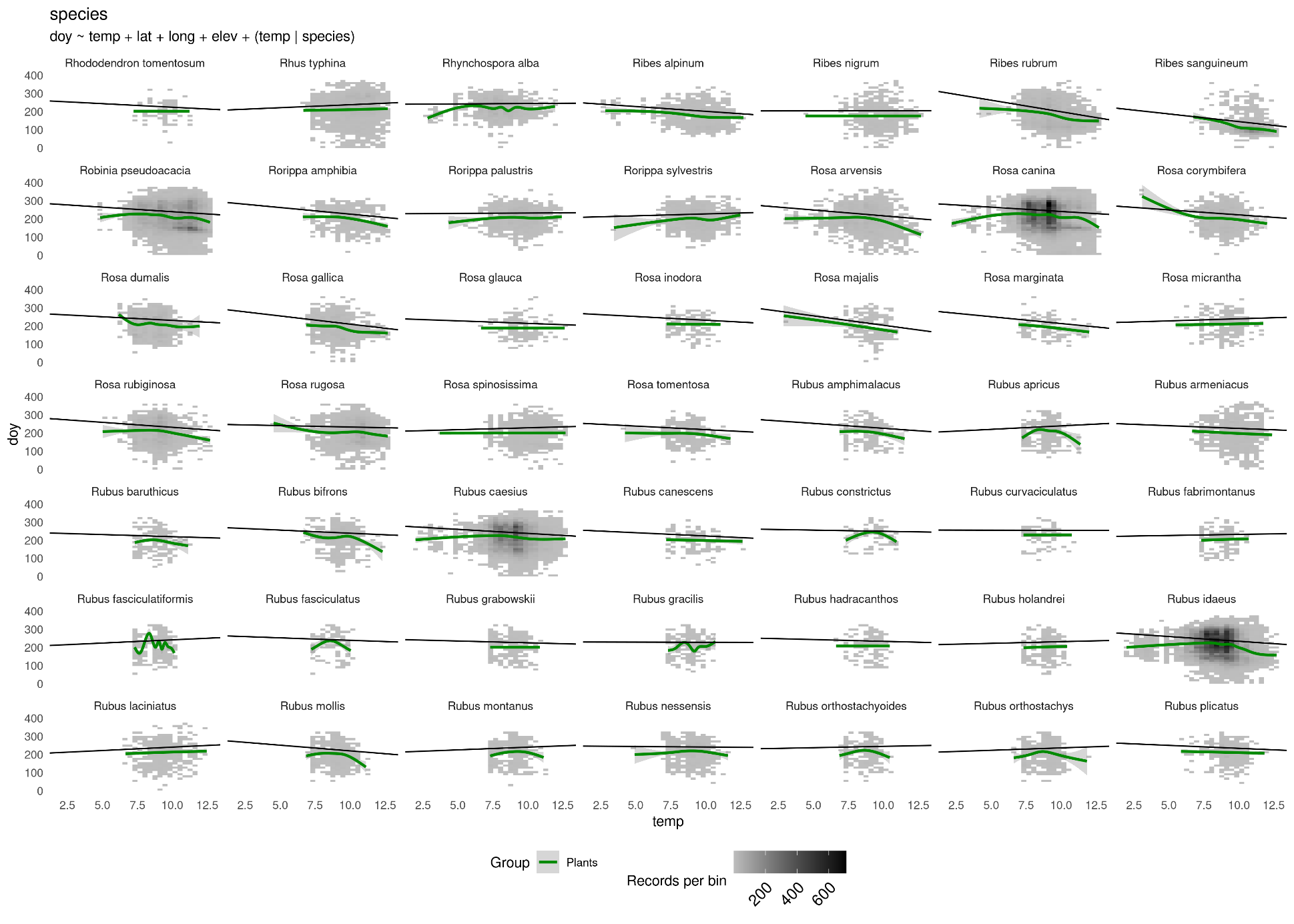

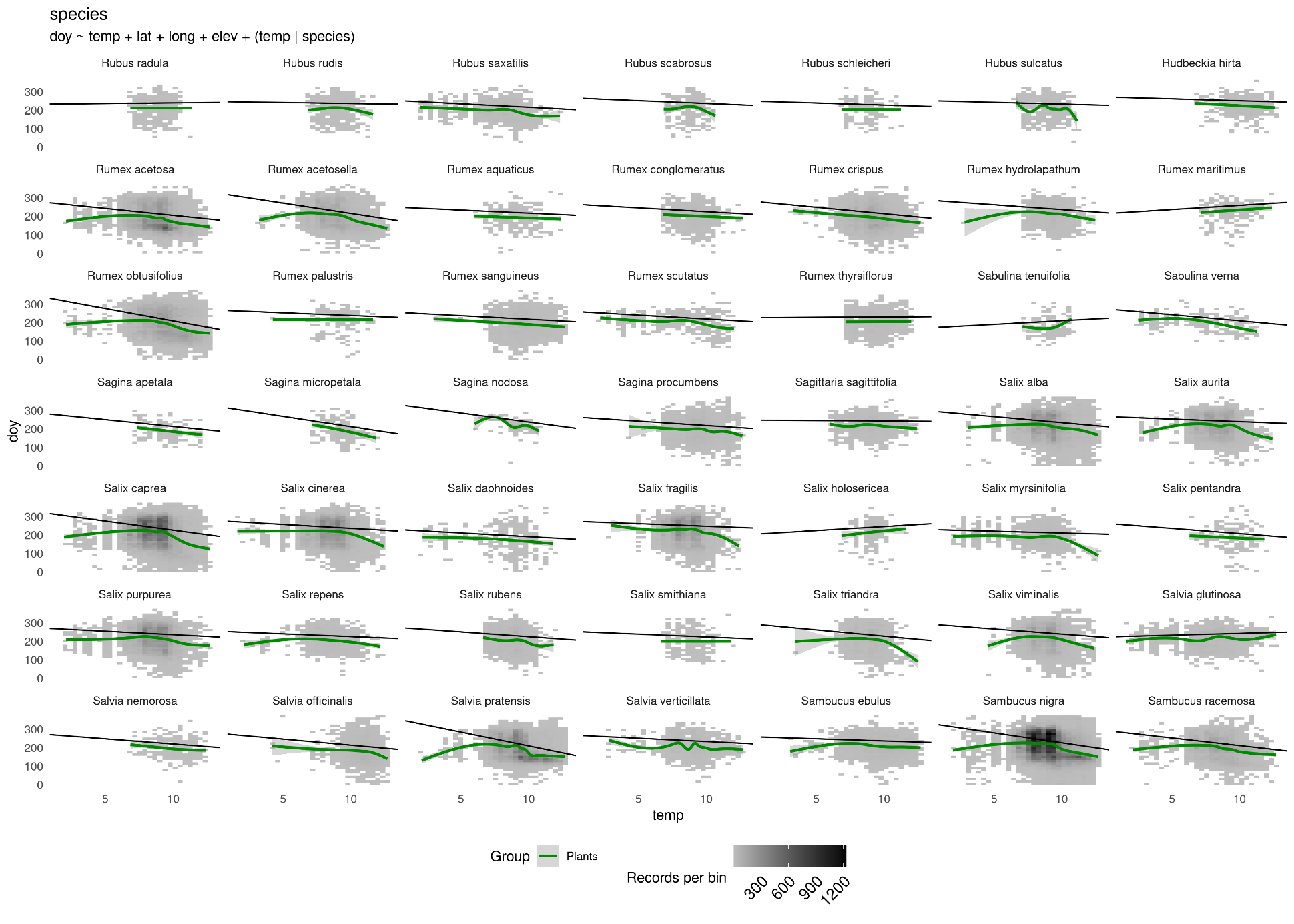

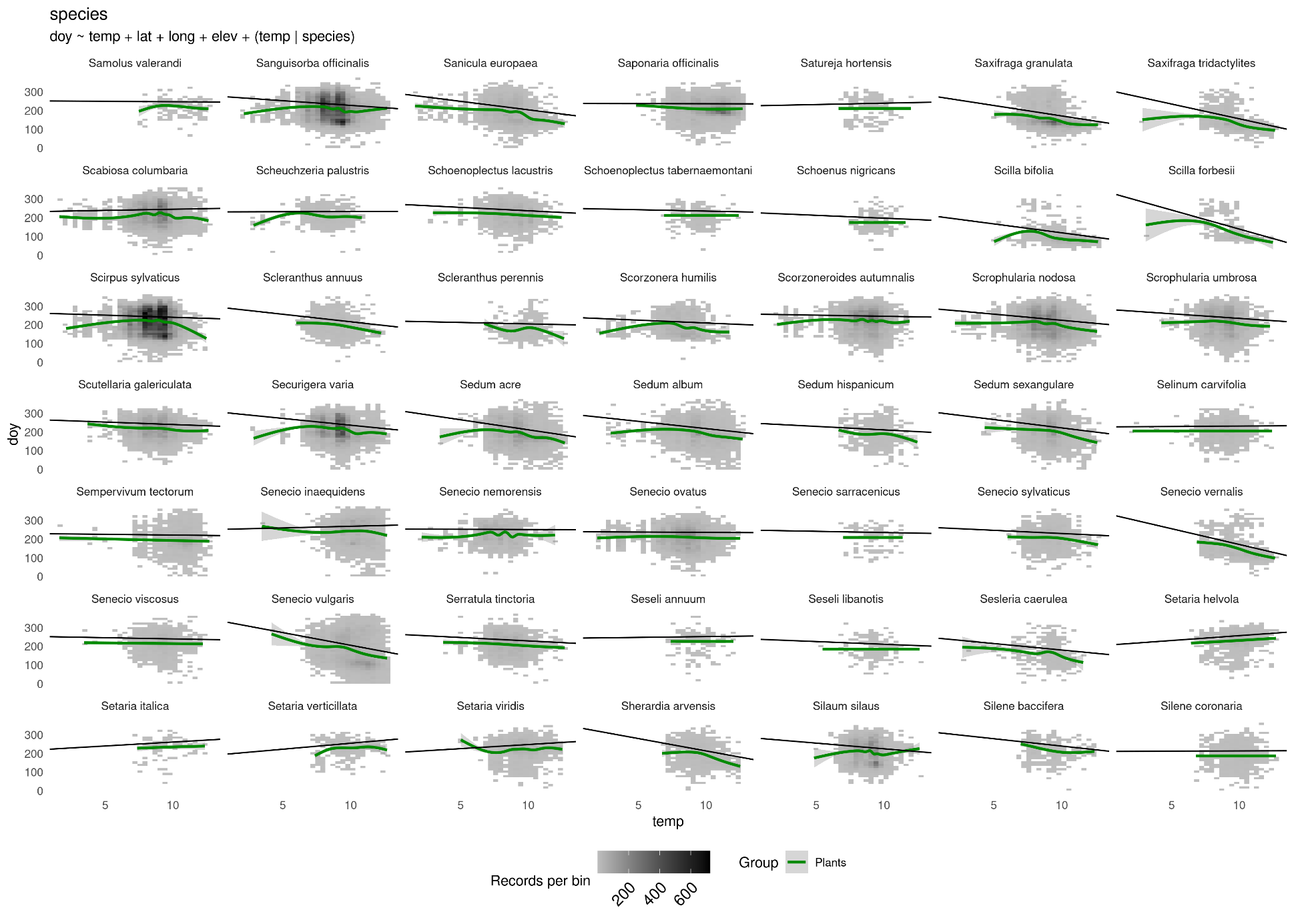

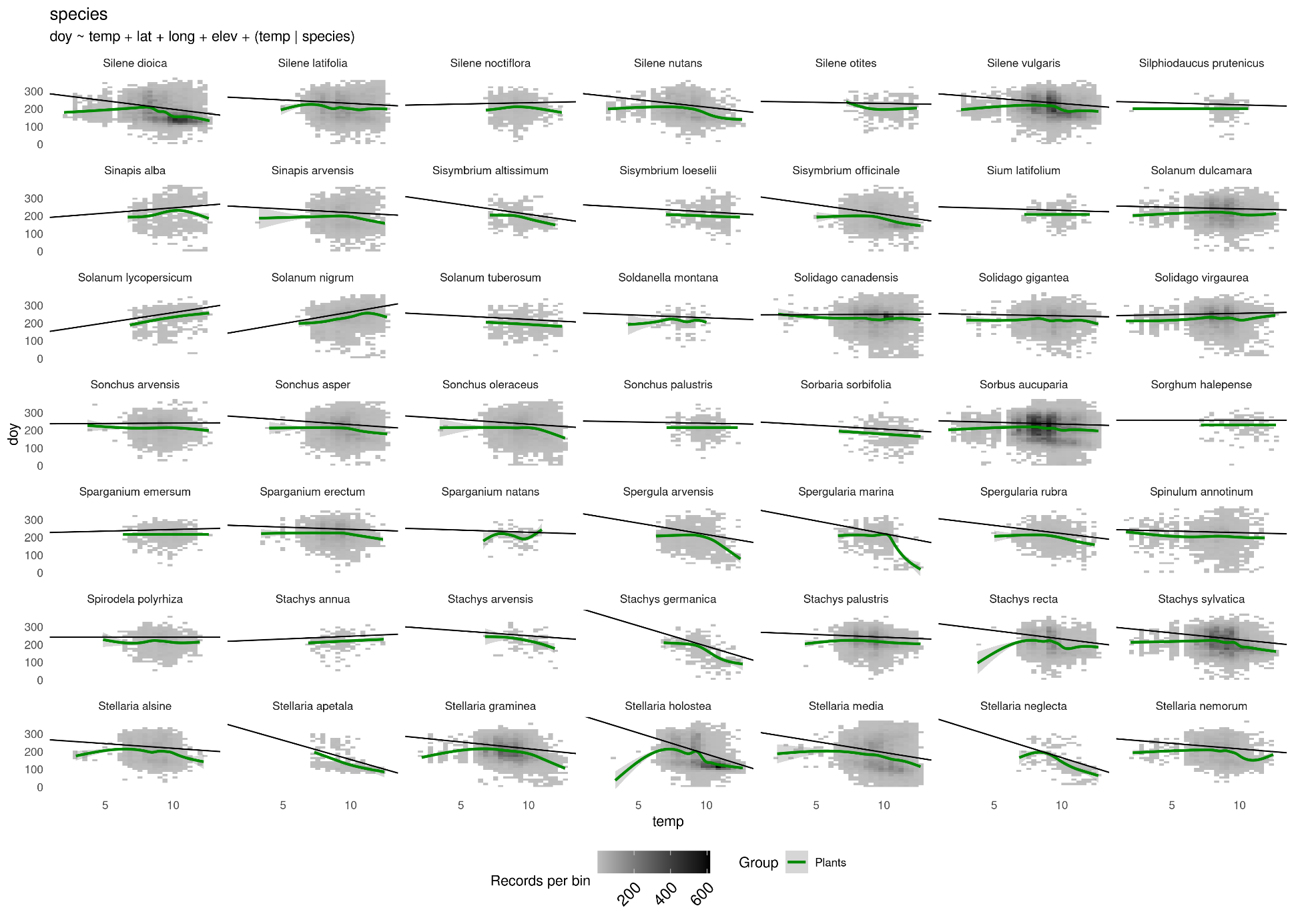

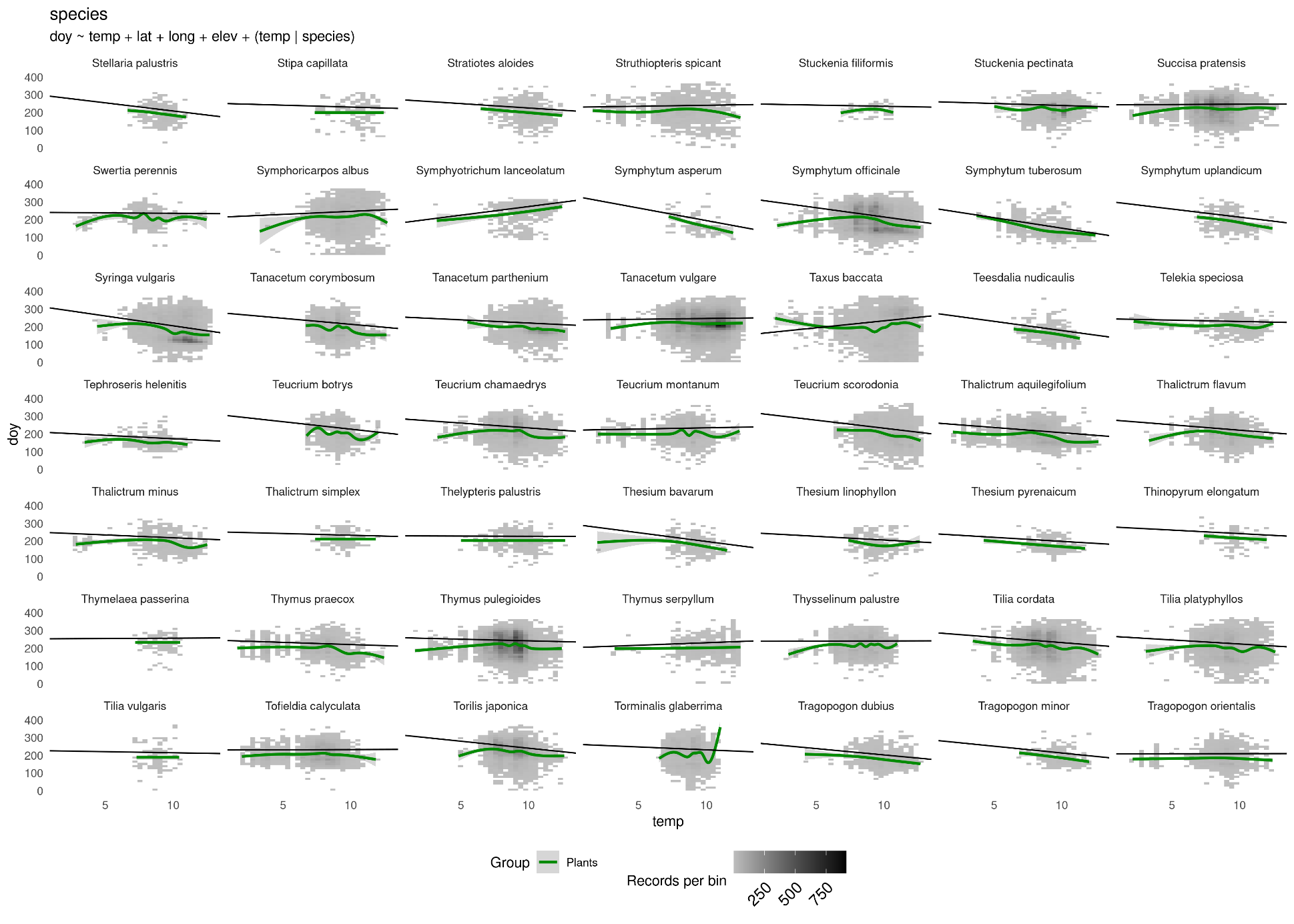

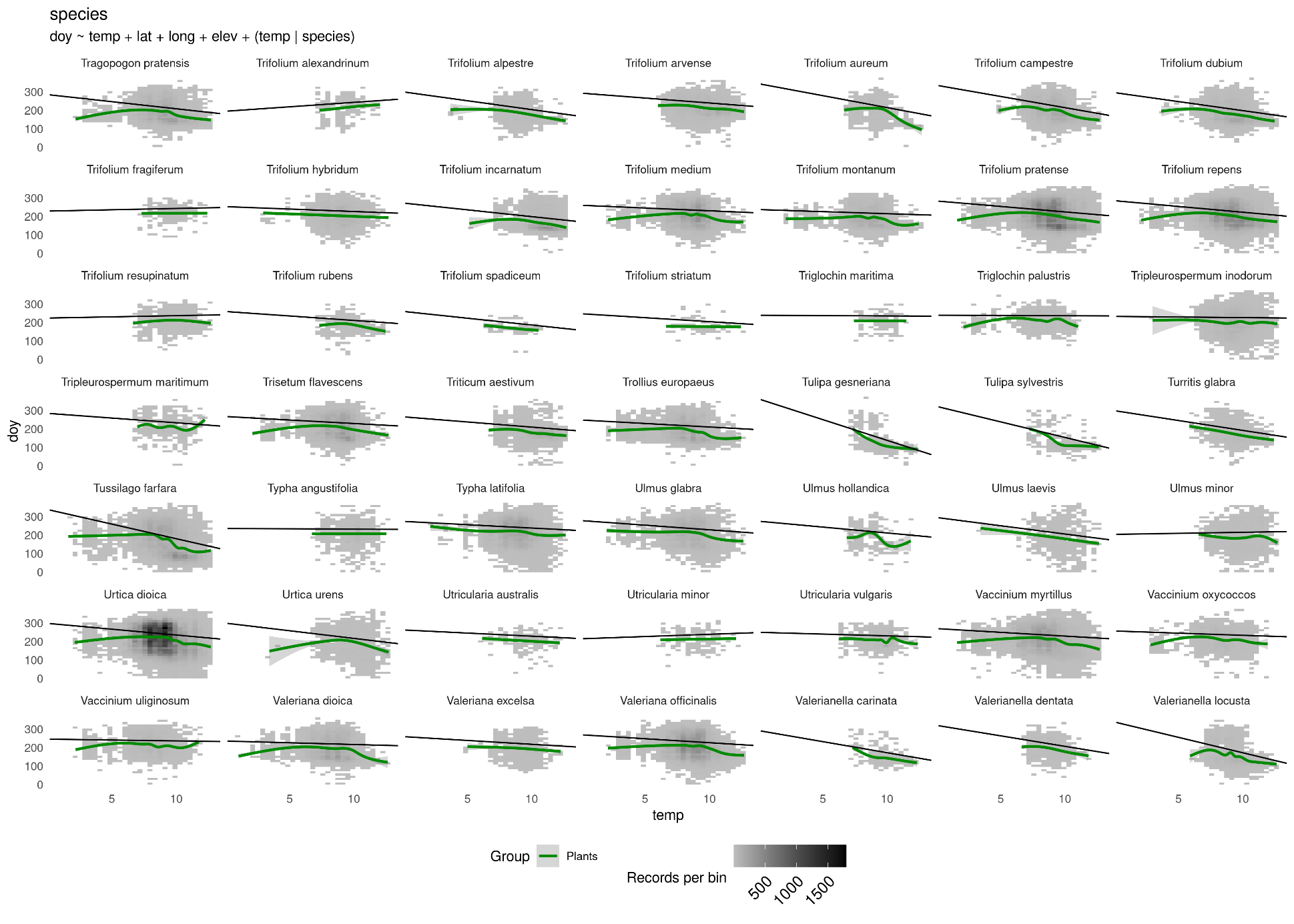

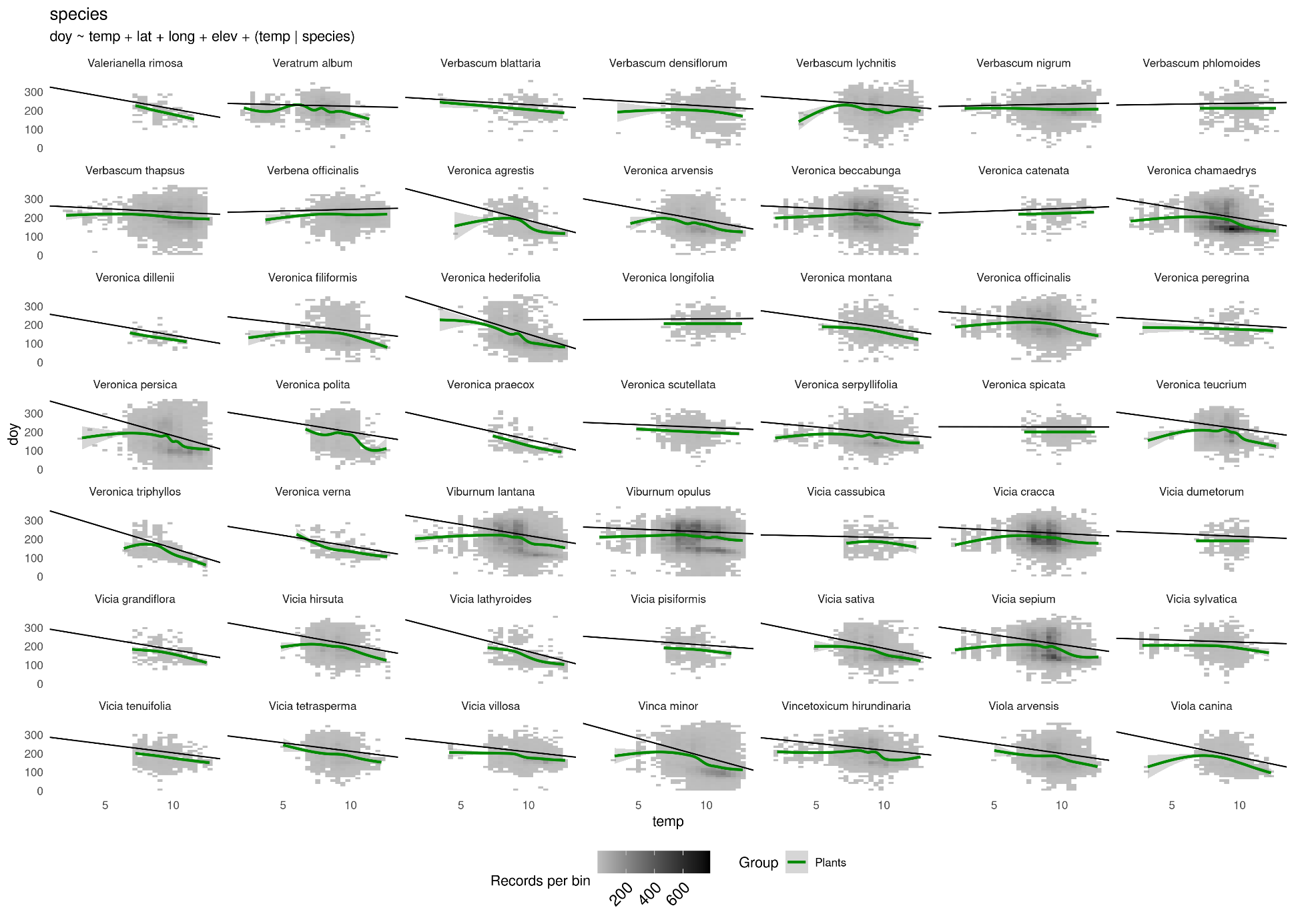

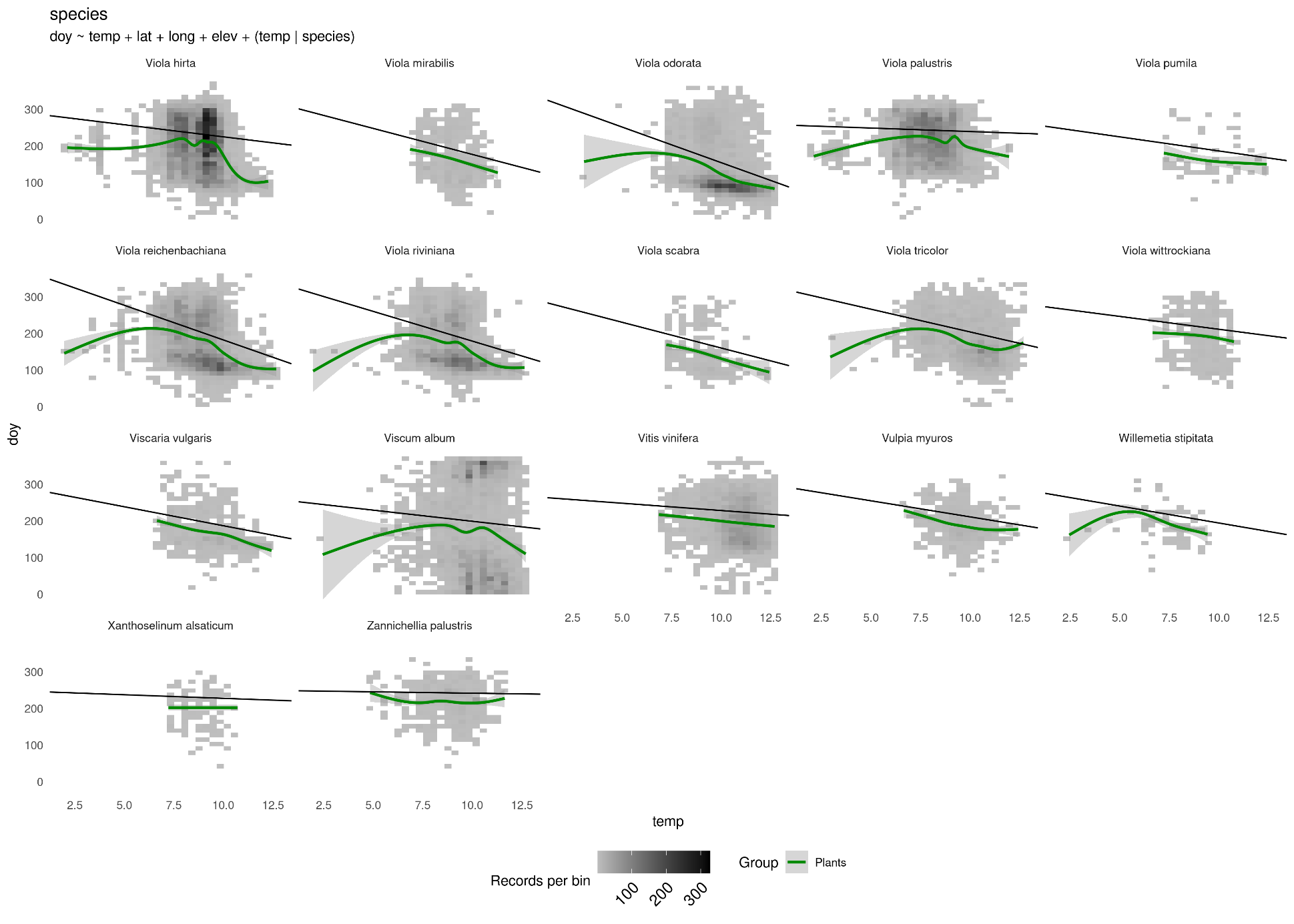
