## Supplementary diagnostic plots 2 for "Climate warming changes synchrony of plants and pollinators"

Supplementary Material - Supplementary plots

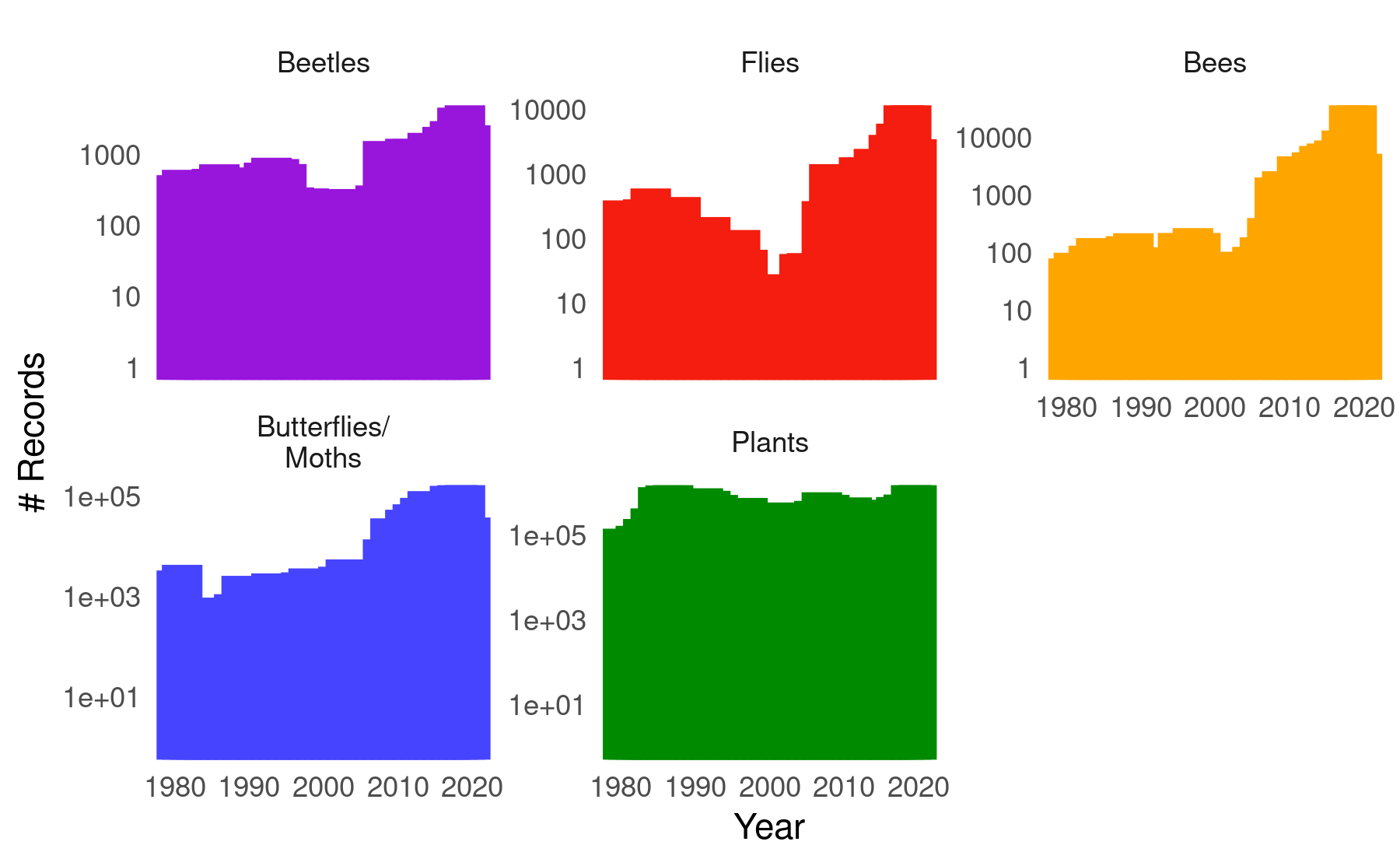

**Figure S 1** The numbers of occurrence records per year for each of the studied taxonomic groups. Note the different scales of the y-axes, and their log-transformation.

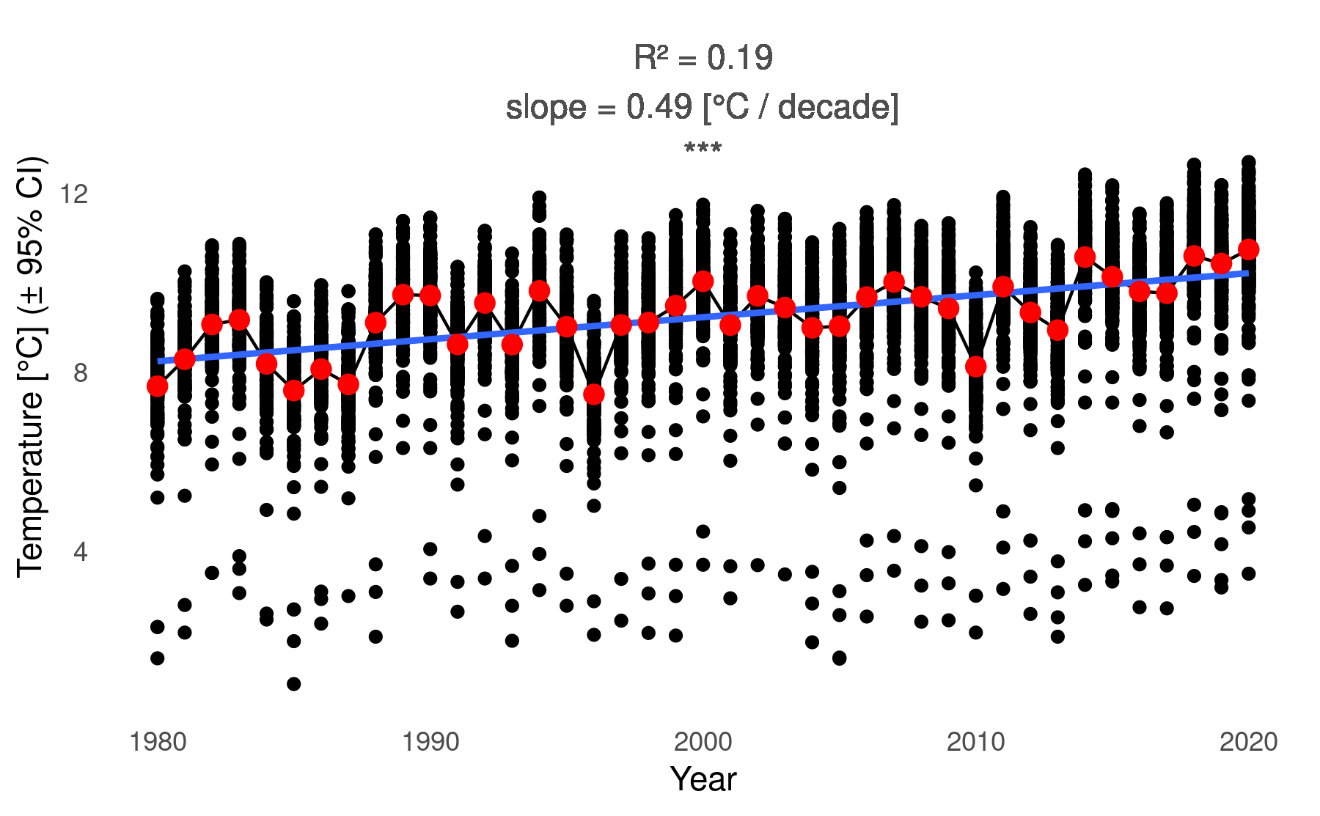

**Figure S 2** Changes in annual mean temperature across Germany. The black dots are individual climate tiles, red symbols are annual mean temperatures (across climate tiles), and the blue line a linear regression of individual temperatures across years (highly significant at $P<0.001$).
